# Beyond Cross-Sectional BARTharm: A Comprehensive Bayesian Framework for Longitudinal Harmonisation, Posterior Prediction, and Uncertainty Quantification^⋆^

**DOI:** 10.64898/2026.11.27.769670

**Authors:** Yasamin Ghassaei, Lawa Dilan, Emma Prevot, Thomas E. Nichols, Habib Ganjgahi

## Abstract

Multi-site magnetic resonance imaging (MRI) studies require harmonisation of image-derived phenotypes (IDPs) to reduce scanner-induced variability while preserving biological information to allow accurate analysis. In longitudinal studies, harmonisation must also account for the correlation between repeated measurements from the same individual, which cross-sectional methods do not explicitly model. In this paper, we extend the original cross-sectional formulation of BARTharm, a Bayesian nonparametric harmonisation tool which uses image quality metrics (IQMs) and Bayesian Additive Regression Trees (BART), to longitudinal data. Long-BARTharm incorporates subject-specific random intercepts and slopes to account for within-subject dependence of repeated observations. Taking advantage of the Bayesian formulation of BARTharm, we also further develop posterior predictive harmonisation and uncertainty-aware downstream inference for both cross-sectional BARTharm and Long-BARTharm. We use posterior samples to harmonise new observations without repeating computationally demanding MCMC estimation and to quantify uncertainty in harmonised measurements beyond point estimates. Through simulation studies we show that Long-BARTharm removes scanner-related variation more effectively than LongCom-Bat across all evaluated scenarios. In real-data analysis, Long-BARTharm reduces scanner effects and residual variance heterogeneity while preserving longitudinal cortical-thickness associations. The posterior predictive harmonisation closely reproduces full-data results, while accounting for harmonisation uncertainty yields stronger statistical evidence for biological associations than unweighted analyses. Together, these developments establish a flexible Bayesian framework for cross-sectional and longitudinal neuroimaging harmonisation that preserves biologically meaningful variation and leverages posterior inference to support predictive harmonisation and uncertainty-aware downstream analyses.

## 1 Introduction

The integration of neuroimaging data across multiple studies and institutions enables analyses of larger and more diverse populations, providing opportunities to investigate biological and clinical associations that may be difficult to characterise in individual cohorts. However, differences in scanner hardware and acquisition protocols introduce technical heterogeneity that can obscure biologically relevant variation or lead to spurious associations in downstream analyses (Bayer et al., 2022; Fortin et al., 2018). This is relevant for image-derived phenotypes (IDPs), such as regional brain volumes and cortical thickness, which provide quantitative measures of brain anatomy derived from magnetic resonance imaging (MRI), as these are sensitive to acquisition-related factors (Han et al., 2006). Neuroimaging harmonisation methods aim to mitigate these effects while preserving biologically meaningful variation. Conventional approaches, such as ComBat, typically rely on categorical scanner identifiers and parametric location–scale adjustments to account for acquisition-related variability (Fortin et al., 2018). Although extensions have introduced greater modelling flexibility, they generally retain these assumptions, potentially limiting their ability to capture complex scanner effects and within-scanner variability (Pomponio et al., 2020). To address these limitations, we previously introduced BARTharm, a Bayesian nonparametric harmonisation framework that uses Bayesian Additive Regression Trees (BART) to separately model biological and acquisition-related contributions to IDPs (Prevot et al., 2025). By incorporating image quality metrics (IQMs), which provide continuous, observation-level descriptors of MRI characteristics, BARTharm captures acquisition-related variability beyond categorical scanner effects, while the flexibility of BART allows both linear and nonlinear relationships and interactions to be modelled without imposing predefined functional forms (Chipman et al., 2010; Esteban et al., 2017). Furthermore, its fully Bayesian formulation provides posterior distributions over biological and acquisition-related effects, establishing a foundation for further methodological developments in harmonisation and downstream statistical inference.

One important setting for extending this framework is longitudinal neuroimaging, where repeated structural MRI measurements enable the characterisation of within-subject changes in brain anatomy over time. Such measurements provide information about individual ageing trajectories that may not be adequately captured by cross-sectional age-related associations (Di Biase et al., 2023). However, repeated observations introduce within-subject dependence, while scanner-related differences across follow-up measurements may complicate the interpretation of longitudinal changes. Cross-sectional harmonisation methods do not explicitly account for this dependence, treating repeated observations as independent and potentially failing to preserve subject-specific trajectories. Longitudinal harmonisation methods must therefore account for individual differences in baseline measurements and rates of change over time, while distinguishing acquisition-related variability from biological effects (Beer et al., 2020; Gaiser et al., 2024). Despite the increasing use of longitudinal neuroimaging data, harmonisation methods specifically designed to accommodate repeated measurements have received less methodological attention than their cross-sectional counterparts. To address this, longitudinal ComBat (LongComBat) extends the conventional ComBat framework by incorporating subject-specific random intercepts within a linear mixed-effects formulation. However, it retains the categorical scanner-based location–scale adjustments of the original method (Beer et al., 2020). Building on the original BARTharm framework, our first contribution is the longitudinal extension of BARTharm (Long-BARTharm), which introduces subject-specific random intercepts and slopes to account for individual differences in baseline measurements and rates of change over time. The extended model retains the flexibility of BART to capture both linear and nonlinear biological and acquisition-related effects, while continuing to leverage IQMs to characterise technical variability beyond categorical scanner identifiers. Long-BARTharm also preserves the heteroskedastic formulation of the original framework, allowing residual variability to differ across scanners.

Beyond accommodating longitudinal data, the fully Bayesian formulation of BARTharm provides opportunities to address broader challenges in neuroimaging harmonisation. Both cross-sectional BARTharm and Long-BARTharm rely on Bayesian backfitting Markov chain Monte Carlo (MCMC) sampling for posterior inference (Chipman et al., 2010; Hastie & Tibshirani, 2000). Although computationally demanding, particularly for large neuroimaging datasets, MCMC provides posterior samples that characterise uncertainty in model estimates rather than only point estimates. In this work, we exploit these samples to introduce two additional contributions. First, we develop posterior predictive harmonisation for both cross-sectional BARTharm and Long-BARTharm, enabling previously unseen observations to be harmonised without repeating the full MCMC inference procedure. By evaluating previously fitted BART ensembles across retained posterior draws, acquisition-related effects can be estimated for new observations, reducing the computational burden associated with repeated model fitting. This is relevant for large or evolving neuroimaging studies, including longitudinal settings where additional follow-up measurements become available over time. Second, beyond providing harmonised point estimates, posterior sampling enables uncertainty associated with individual harmonised measurements to be quantified, an aspect typically not explicitly considered in downstream analyses following conventional harmonisation. Importantly, this uncertainty can vary across observations and brain regions, resulting in differences in the precision of harmonised estimates (Reynolds et al., 2023). Such variation raises broader concerns about treating estimated quantities as known in subsequent statistical analyses, as uncertainty introduced during an initial estimation stage can have implications for downstream inference (Murphy & Topel, 1985). To investigate the potential relevance of harmonisation uncertainty to downstream analyses, we use posterior draws from BARTharm and Long-BARTharm to estimate observation-specific harmonisation uncertainty and incorporate inverse-variance weights into downstream regression analyses, allowing measurements to contribute according to their estimated precision.

Taken together, these developments build upon the original BARTharm methodology to establish a more comprehensive Bayesian nonparametric framework for neuroimaging harmonisation. The proposed framework encompasses three complementary methodological contributions. First, we introduce Long-BARTharm, extending the original cross-sectional formulation to longitudinal neuroimaging data through subject-specific random intercepts and slopes, while retaining the flexibility of BART to model both linear and nonlinear biological and acquisition-related effects and the use of IQMs to characterise technical variability. Second, we develop posterior predictive harmonisation for both cross-sectional BARTharm and Long-BARTharm, enabling previously unseen observations to be harmonised using posterior samples from an existing fitted model without repeating the computationally demanding MCMC estimation procedure. Third, we leverage posterior samples to quantify observation-specific harmonisation uncertainty and investigate its incorporation into downstream statistical analyses through inverse-variance weighted regression.

We evaluate these methodological developments using fully synthetic and semi-synthetic simulations, alongside real-world neuroimaging data from the Alzheimer’s Disease Neuroimaging Initiative (ADNI) (Petersen et al., 2010). Our evaluation examines harmonisation performance, the preservation of biological associations and longitudinal trajectories, posterior predictive agreement, and the implications of uncertainty weighting for downstream statistical inference.

## 2 Materials & Methods

We first describe LongComBat and Long-BARTharm, the two methods we compare for harmonising repeated MRI measurements. Both aim to reduce scanner effects while accounting for dependence between visits and preserving biological changes within subjects over time. For a given IDP, let *y_ij_*denote its value for subject *i* at visit *j*, where *i* = 1*, . . . , N* and *j* = 1*, . . . , n_i_*. We denote the scanner used at that visit by *s*(*i, j*).

### 2.1 LongComBat

LongComBat is a harmonisation method that extends ComBat to repeated measurements using a linear mixed-effects model (Beer et al., 2020). Like ComBat, it adjusts for scanner differences in mean and residual scale while retaining the fitted effects of biological covariates. The longitudinal extension adds a subject-specific random intercept to account for dependence between repeated scans of the same subject. For a scan acquired on scanner *s* = *s*(*i, j*), the model is

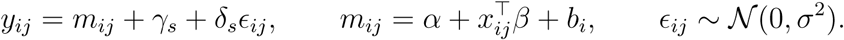

Here, *α* is the fixed intercept, *x_ij_* is the covariate vector comprising the biological predictors and the time of the visit, and *β* is the corresponding vector of fixed-effect coefficients. The random intercept *b_i_* is shared across visits for subject *i*. Each scanner *s* has an additive shift *γ_s_* and a residual standard deviation multiplier *δ_s_ >* 0.

The estimated harmonised IDP is

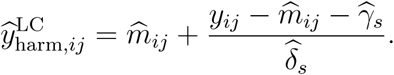

This adjustment removes the estimated scanner shift and rescales the residual while retaining the fitted covariate and subject-specific effects. When several IDPs are available, empirical Bayes estimation pools information across IDPs within each scanner.

Longitudinal ComBat inherits two limitations from ComBat. It requires a scanner ID for each observation and specifies a linear structure for the effects of biological covariates. The longitudinal model also includes only a subject-specific random intercept, so it does not estimate subject-specific slopes.

### 2.2 Long-BARTharm

#### 2.2.1 BART

Bayesian additive regression trees (BART) express the conditional mean of a continuous outcome *y* as a sum of *m* decision trees,

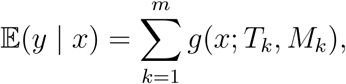

where *T_k_*contains the splitting rules for tree *k* and *M_k_* contains its terminal-node values (Chipman et al., 2010). Splits on a single predictor can capture its main effect, while splits on different predictors along the same path can capture an interaction by allowing the effect of one predictor to depend on another. By summing the piecewise-constant contributions of individual trees, BART can capture both linear and non-linear relationships that vary in complexity across the covariate space. In Long-BARTharm, this flexibility allows us to estimate the scanner-related and biological effects without specifying the functional forms or interactions within either component in advance.

#### 2.2.2 Longitudinal model and harmonisation

Cross-sectional BARTharm separates scanner-related and biological variation using two independent tree ensembles (Prevot et al., 2025). We extend this model by adding subject-specific random intercepts and slopes to account for repeated measurements. The model hierarchy is

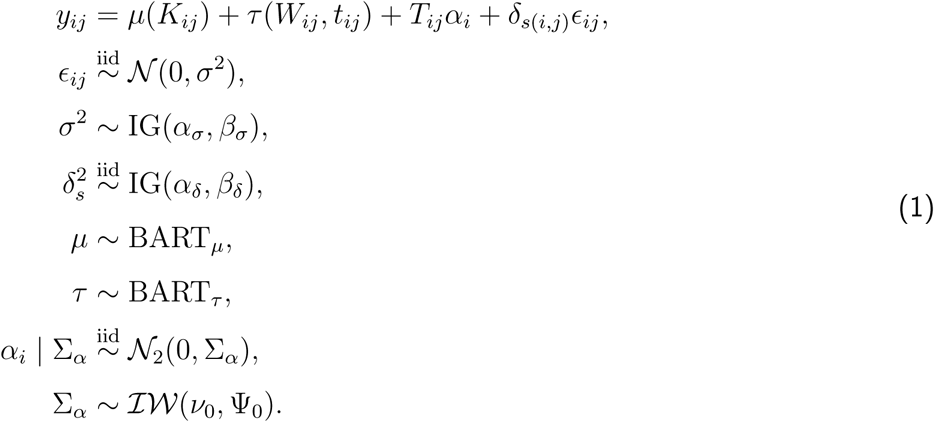

Here, *K_ij_* is the vector of scanner-related predictors for the scan acquired from subject *i* at visit *j*, *W_ij_*is the vector of biological covariates, and *t_ij_* is the time of the observation. We separately model the effects of the biological covariates and IQMs on the observation *y_ij_* by placing independent nonparametric BART priors on the scanner-related component *µ* and the biological component *τ* . The scanner-specific multiplier *δ_s_ >* 0 allows residual variability to differ across scanners, with residual standard deviation *σδ_s_* for scanner *s*.

We use 200 trees for the scanner component *µ* and 50 for the biological component *τ* . The prior probability that a node at depth *d* splits is *γ*(1 + *d*)*^−β^*. We set (*γ, β*) = (0.95, 2) for the scanner trees and (0.25, 3) for the biological trees. These settings favour shallower biological trees, resulting in stronger shrinkage towards homogeneous effects (Hahn et al., 2020).

The term *T_ij_α_i_* represents the subject-specific random effects, with design vector *T_ij_* = (1*, t_ij_*) and coefficient vector *α_i_* = (*α_i_*_1_*, α_i_*_2_)*^⊤^*, where *α_i_*_1_ and *α_i_*_2_ are the random intercept and slope, respectively. These effects capture individual deviations from the population-level trajectory and account for within-subject dependence. They are independent of the residual errors. We assign the coefficients a multivariate Gaussian prior, and an inverse-Wishart prior on their variance-covariance matrix Σ*_α_* (Spanbauer & Sparapani, 2021). In our analysis, we set *ν*_0_ = 2 and Ψ_0_ = *I*_2_.

We fit the model to standardised IDPs, *y_ij_* = (*Y_ij_ − ̄Y*)*/S_Y_* , where *Y_ij_* is the observed IDP and *̄Y* and *S_Y_* are its training-data mean and standard deviation.

The global residual variance *σ*^2^ and the scanner-specific variance multipliers *δ_s_*^2^ have inverse-gamma priors. We set *α_σ_*= *β_σ_* = *α_δ_* = *β_δ_* = 0.01, as we intend the priors to be non-informative. Observations from scanners with greater residual variability receive less weight when estimating the tree ensembles and subject-specific effects, with weights proportional to 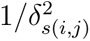.

The likelihood identifies only the products σ^2^*δ_s_*^2^, so the global variance *σ*^2^ and the scanner-specific multipliers *δ_s_*^2^ are not separately identifiable. We sample in this expanded parameter space and impose an identifiability constraint by reparameterising each retained posterior draw. Specifically, within each draw, we divide the scanner-specific scales *δ_s_* by their geometric mean and multiply *σ* by the same factor. The aligned scales have unit geometric mean. This transformation preserves each product σ^2^*δ_s_*^2^ draw by draw and therefore preserves both the likelihood and the posterior distribution of the scanner-specific residual variances. We use the aligned draws for harmonisation.

### Harmonised-outcome transformation

We obtain posterior draws of the model parameters and fitted functions by Markov chain Monte Carlo (MCMC), as described in Section 2.2.3. Let 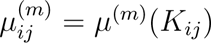 and 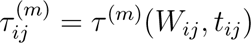 denote the scanner and biological effects at draw *m*. At each retained posterior draw *m*, we harmonise observation (*i, j*) using

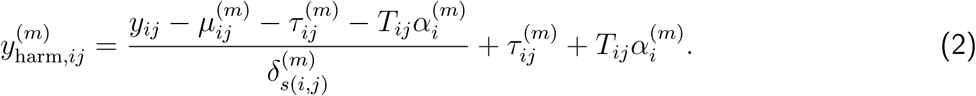

This removes the estimated scanner-related mean effect and adjusts the residual scale, while retaining the estimated biological and subject-specific components.

We return these harmonised draws to the original IDP scale and average across the *M* retained draws to obtain the point estimates,

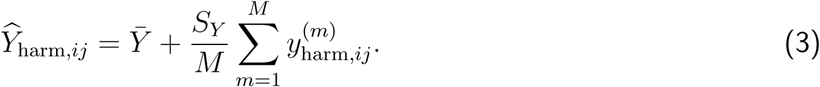

#### 2.2.3 Model fitting

We fit Long-BARTharm separately for each IDP using a Gibbs sampler that combines BART and mixed-effects updates (Spanbauer & Sparapani, 2021; Tan & Roy, 2019). At each iteration, we update the scanner-related ensemble *µ*, followed by the biological ensemble *τ* . We then draw the subject-specific random effects and their covariance matrix Σ*_α_*. Finally, we draw the scanner-specific variance multipliers *δ_s_*^2^ and the global residual variance *σ*^2^.

BART uses Bayesian backfitting to sample from the posterior distribution of the tree ensemble. At each Markov chain Monte Carlo (MCMC) iteration, the algorithm updates each tree’s structure with a Metropolis–Hastings step and its terminal-node values with a Gibbs step (Chipman et al., 2010). To update the scanner ensemble *µ*, we use the residuals 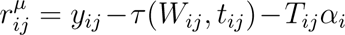 as the response. These are the observed IDPs after subtracting the current biological and subject-specific effects. To update the biological ensemble *τ* , we subtract the newly sampled scanner effect and the current subject-specific effect, giving 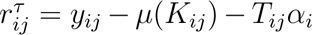. Both ensemble updates use observation weights 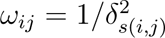.

Given the updated ensemble values *µ_ij_* = *µ*(*K_ij_*) and *τ_ij_* = *τ* (*W_ij_, t_ij_*), we calculate the residuals for the subject-specific effects, *r_ij_^α^*= *y_ij_ − µ_ij_ − τ_ij_*. We use these residuals and the observation weights *ω_ij_* to draw each subject’s random intercept and slope from their joint conditional distribution. We then draw the shared covariance matrix Σ*_α_* from its full conditional distribution.

Finally, we calculate the residuals from the full model, *r_ij_*= *y_ij_ − µ_ij_ − τ_ij_ − T_ij_α_i_*. For each scanner *s*, we use its residuals and the current *σ*^2^ to draw *δ_s_*^2^ from its full conditional distribution. We then update the observation weights and draw *σ*^2^ from its full conditional distribution using the residuals and updated weights.

We implement the sampler in R (R Core Team, 2023) using the SoftBart package (Linero, 2022). The package allows us to update the scanner and biological ensembles directly within our Gibbs sampler without modifying the underlying C++ code. Both ensembles use soft decision trees, in which the predictor value at each split determines the probability of following either branch. This allows the fitted response to vary smoothly with continuous predictors (Linero & Yang, 2018). Before fitting, we standardise each IDP using its training-data mean and standard deviation and quantile-transform the continuous predictors other than visit time. Algorithm 1 summarises the sampler.

**Algorithm 1**: Gibbs sampler for Long-BARTharm Each line uses the most recently sampled values. For *µ* and *τ* , the notation *∼*[ *· | ·* ] denotes one weighted backfitting sweep targeting the indicated conditional posterior.

|  |  |
|---|---|
| 1 | Initialise $\tau_{ij}^{(0)} \leftarrow 0$ and $\alpha_i^{(0)} \leftarrow \mathbf{0}$. |
| 2 | Initialise $\sigma^{(0)}$, $\{\delta_s^{(0)}\}_{s=1}^{S}$ and $\Sigma_\alpha^{(0)}$. Set $\omega_{ij}^{(0)} \leftarrow 1/(\delta_{s(i,j)}^{(0)})^2$. |
| 3 | **for** $\mathrm{iter} \leftarrow 1$ **to** $\mathrm{max_iter}$ **do** |
| 4 | $r_{ij}^{\mu} \leftarrow y_{ij} - \tau_{ij} - T_{ij}\alpha_i$ |
| 5 | $\mu \sim [\mu \mid r^{\mu}, K, \omega, \sigma^2]$ |
| 6 | $r_{ij}^{\tau} \leftarrow y_{ij} - \mu_{ij} - T_{ij}\alpha_i$ |
| 7 | $\tau \sim [\tau \mid r^{\tau}, W, t, \omega, \sigma^2]$ |
| 8 |  |
| 9 | $r_{ij}^{\alpha} \leftarrow y_{ij} - \mu_{ij} - \tau_{ij}$ |
| 10 | $\alpha_i \sim [\alpha_i \mid r_i^{\alpha}, T_i, \omega_i, \sigma^2, \Sigma_\alpha]$ for each subject $i$ |
| 11 | $\Sigma_\alpha \sim [\Sigma_\alpha \mid \{\alpha_i\}_{i=1}^{N}]$ |
| 12 |  |
| 13 | $r_{ij} \leftarrow y_{ij} - \mu_{ij} - \tau_{ij} - T_{ij}\alpha_i$ |
| 14 | $\delta_s^2 \sim [\delta_s^2 \mid \{r_{ij} : s(i,j) = s\}, \sigma^2]$ for each scanner $s$ |
| 15 | $\omega_{ij} \leftarrow 1/\delta_{s(i,j)}^2$ |
| 16 | $\sigma^2 \sim [\sigma^2 \mid r, \omega]$ |
| 17 | **Store** $\mu_{ij}^{\mathrm{iter}},\ \tau_{ij}^{\mathrm{iter}},\ \alpha_i^{\mathrm{iter}},\ T_{ij}\alpha_i^{\mathrm{iter}},\ \Sigma_\alpha^{\mathrm{iter}},\ \sigma^{\mathrm{iter}},\ \{\delta_s^{\mathrm{iter}}\}_{s=1}^{S}$ for retained draws. |
| 18 | **end for** |

We inspect trace plots of the variance components, particularly the global residual variance *σ*^2^, to assess convergence and guide the choice of burn-in and thinning intervals. For each posterior draw retained after burn-in and thinning, we align the scanner multipliers *δ_s_*and the global scale *σ* as described in Section 2.2.2. We then apply Equation (2) to each observation at each retained draw and average its harmonised values on the original IDP scale.

#### 2.2.4 Posterior predictive harmonisation

Posterior predictive harmonisation uses the posterior distribution obtained from the training data to harmonise new observations without repeating MCMC inference. For each retained posterior draw, we evaluate the scanner-related and biological BART ensembles at the covariates for the new scan.

Cross-sectional BARTharm can harmonise scans from new subjects. In the longitudinal setting, we harmonise new visits from subjects represented in the training data. For each retained iteration, we reuse the posterior draw of the subject’s random intercept and slope from the training fit to calculate the subject-specific effect at the time of the new visit.

We standardise the new observed IDP using the training-set mean and standard deviation and harmonise it draw by draw using Equation (2). In the cross-sectional setting, we omit the subject-specific term. We return the harmonised draws to the original IDP scale and average them to obtain a point estimate.

We expect posterior predictive harmonisation to be most reliable when the joint distribution of the covariates in the new data is similar to that in the training data. In addition, each new scan must come from a scanner *s* represented in the training data so that its aligned scale multipliers *δ*^(*m*)^ are available. The same requirement applies to the other categorical predictors, whose levels in the new scans must all have been observed during training.

#### 2.2.5 Uncertainty quantification and downstream inference

Posterior sampling allows us to quantify uncertainty in the estimated model components and, consequently, in the harmonised measurements. Again, let 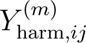 denote the harmonised IDP on its original scale for observation (*i, j*) in retained posterior draw *m*. We estimate its posterior variance and define the corresponding inverse-variance weight as

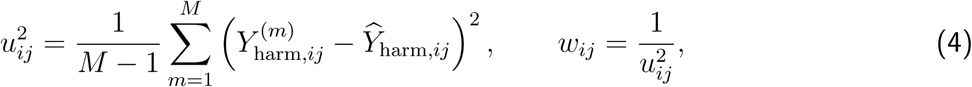

where *M* is the number of retained draws and *Y*_harm*,ij*_ is their mean. This calculation is the same in the cross-sectional and longitudinal settings. It measures harmonisation uncertainty conditional on the observed data.

We fit downstream regression models to the harmonised IDPs using weights *w_ij_*, where observations with lower harmonisation uncertainty receive greater weight. We compare weighted and unweighted fits using the same covariates to assess the effect of uncertainty weighting on downstream inference.

### 2.3 Simulation studies

We first assess how well Long-BARTharm removes scanner-related variation while preserving biological differences and subject-specific changes over time. We consider both a fully synthetic setting and a semi-synthetic setting, where we use covariates from a real dataset. In each setting, we vary whether the biological and scanner-related effects are linear or non-linear, giving four scenarios.

#### 2.3.1 Simulation design

For subject *i* at visit *j*, we generate the observed IDP as

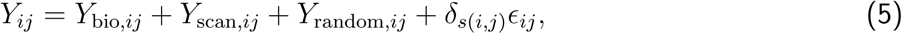

where *Y*_bio_*_,ij_* is the contribution of the biological covariates, *Y*_scan_*_,ij_* is the contribution of the scanner-related effects, and *Y*_random_*_,ij_* is the subject-specific trajectory generated by a random intercept and slope. The multiplier *δ_s_*_(_*_i,j_*_)_ scales the residual *ɛ_ij_* according to the scanner used at that visit.

The target of harmonisation is the clean outcome

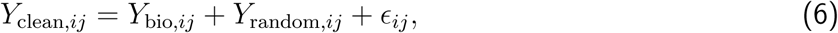

which retains the biological contribution, the subject-specific trajectory and the underlying residual.

In both settings, we generate the biological effect from the biological covariates and the scanner effect from the IQMs. The linear specifications use weighted sums of the predictors, while the non-linear specifications include quadratic terms, cubic terms and pairwise interactions. In each realisation, we independently draw a random intercept and slope for each subject to generate their trajectory. We scale the biological, scanner and subject-specific contributions to their respective target standard deviations so that their magnitudes are comparable across scenarios. We draw a residual for each scan using a global residual standard deviation *σ* that is fixed across realisations. To allow residual variability to differ across scanners, we draw one positive multiplier *δ_s_* per scanner and normalise the multipliers to have geometric mean one, giving residual standard deviation *σδ_s_* for scanner *s*.

We use a fully-synthetic simulations setting to assess how accurately the methods recover the known scanner-free outcome as we vary the form of the biological and scanner-related effects. Here we simulate 200 subjects with five equally spaced visits each, giving 1,000 scans per dataset. Each of five scanners is assigned 40 subjects, with each subject scanned on the same scanner at every visit. We generate five biological covariates per subject and five IQMs per scan. The biological covariates remain fixed across visits, while the IQMs have scanner-specific means and variances.

To evaluate the methods under realistic patterns of follow-up and scanner use, we perform a semi-synthetic simulation study using real data from the the Alzheimer’s Disease Neuroimaging Initiative (ADNI) dataset (Petersen et al., 2010). We simulate outcomes for 1,787 observed scans from 370 subjects across 48 scanners, with two to eight visits per subject. We generate the biological effect for each subject from 14 demographic and clinical covariates recorded at their first available visit. The scanner effect is generated separately for each scan from its 26 observed IQMs. Within each scenario, the biological and scanner effects remain fixed across realisations because the observed predictors and their coefficients are fixed. This design preserves the empirical longitudinal structure, biological covariate distributions, IQM distributions, and scanner assignments of the real dataset while maintaining a known decomposition into biological, scanner-related, subject-specific, and residual components.

#### 2.3.2 Performance evaluation

For each simulation scenario, we generate 1,000 realisations of the dataset. Each BARTharm MCMC chain runs for 10,000 iterations. After discarding the first 2,000 as burn-in, we retain every second draw, giving 4,000 posterior draws per dataset. We calculate the evaluation metrics separately for each dataset and report their means across realisations.

We compare Long-BARTharm with LongComBat in both simulation settings, using evaluation metrics similar to those used in cross-sectional BARTharm (Prevot et al., 2025). To assess recovery of the clean outcome, we compare *Y*_harm_ with *Y*_clean_ using two metrics. A lower root mean squared error (RMSE) indicates closer agreement in magnitude, while a higher Pearson correlation (BIO Corr) indicates better preservation of biological variability. We also calculate the absolute Pearson correlation between *Y*_harm_ and the scanner component *Y*_scan_ (IQMs Corr), with values closer to zero indicating a weaker remaining association.

For each method, we report *δ* RMSE to assess how accurately it estimates the scanner-specific residual scale multipliers *δ_s_*. Within each realisation, we average the squared differences between the estimated and true multipliers across scanners before taking the square root. Lower values indicate more accurate estimation of the multipliers.

### 2.4 Real-data analysis

#### 2.4.1 ADNI Dataset and Preprocessing

The real-world analyses were conducted using neuroimaging data from the Alzheimer’s Disease Neuroimaging Initiative (ADNI) (Petersen et al., 2010), a longitudinal, multi-site study designed to investigate the progression of Alzheimer’s disease through neuroimaging, clinical, cognitive, and biological assessments. Data were obtained from the ADNI database (http://adni.loni.usc.edu/), focusing on the ADNIGO and ADNI2 study phases. Repeated observations were used for longitudinal harmonisation, while the first available observation from each participant was selected for cross-sectional posterior predictive analyses. Ethical approval for ADNI was obtained from the Institutional Review Boards of participating institutions, and all participants provided written informed consent. Data were accessed and analysed in accordance with ADNI’s Data Use Agreement and Data Sharing and Publication Policy.

Cortical thickness and volume measurements were obtained from the UCSF Longitudinal FreeSurfer (version 5.1) processing collection available through ADNI (Reuter et al., 2012), which incorporates repeated scans to improve longitudinal measurement consistency. The resulting IDPs were used as harmonisation outcomes. Image quality metrics (IQMs) were extracted from structural MRI acquisitions using MRIQC, which computes quantitative, no-reference measures of image quality, including noise, tissue contrast, and intensity non-uniformity (Esteban et al., 2017). As noted in the original BARTharm manuscript (Prevot et al., 2025), IQMs associated with biological characteristics may lead to the removal of biologically relevant variation during harmonisation. Candidate IQMs were therefore evaluated using subject-matter knowledge and screened for associations with biological and clinical characteristics. Pairwise associations were examined using Spearman rank correlations, while multivariable linear regression models assessed their joint associations with each IQM, quantified by the coefficient of determination (*R*^2^). Analyses were restricted to each participant’s first available observation to avoid dependence between repeated measurements.

Scanner configurations were defined by unique combinations of scanner identifier, manufacturer, model, magnetic field strength, receive coil, and acquisition site (Beer et al., 2020). Configurations represented by fewer than two observations were excluded to support the estimation of scanner-specific effects. For the longitudinal dataset, we retained repeated measurements from 358 participants, yielding 1,707 observations across 33 unique scanner configurations, with an average of 4.77 observations per participant. For the cross-sectional dataset, we extracted the first available observation from each participant.

The longitudinal cohort comprised 100 (27.9%) cognitively normal (CN) participants, 216 (60.4%) with mild cognitive impairment (MCI), and 42 (11.7%) with Alzheimer’s disease (AD). The cohort included 177 females (49.4%), with a mean baseline age of 71.93 years (SD = 7.05, range: 55.02–88.70 years) and a mean baseline Mini-Mental State Examination (MMSE) score of 27.84. No participant changed diagnostic group during follow-up. A detailed summary of the cohort’s demographic and clinical characteristics is provided in Supplementary Table 4.

Fourteen biological and clinical covariates were included in the harmonisation model: age, sex, years of education, cognitive diagnosis, handedness, self-identified race, retirement status, five domain-specific Clinical Dementia Rating (CDR) scores (memory, orientation, judgement and problem solving, community affairs, and home and hobbies), global CDR score, and MMSE total score. All covariates were held at their baseline values throughout longitudinal harmonisation.

#### 2.4.2 Longitudinal analysis

Long-BARTharm was compared with LongComBat, implemented using restricted maximum likelihood (REML) estimation and empirical Bayes (EB) adjustment of scanner-specific effects, with participant-specific random intercepts (Beer et al., 2020). Both harmonisation methods incorporated the same 14 baseline biological and clinical covariates, with time included as a fixed effect. For Long-BARTharm, 200 trees were used for the IQM-related component and 50 for the biological component. Posterior inference was based on 55,000 Gibbs sampling iterations, with the first 5,000 discarded as burn-in and every tenth subsequent iteration retained, yielding 5,000 posterior samples per fit. Unharmonised IDPs served as a reference for assessing the effects of harmonisation.

To assess the removal of scanner-related information following harmonisation, a multiclass support vector machine (Cortes & Vapnik, 1995) with a radial basis function kernel was used to predict scanner identity from unharmonised, Long-BARTharm, and LongComBat IDPs. Performance was evaluated using five-fold cross-validation with participant-level splits. Chance-level performance was estimated using 1,000 scanner-label permutations, exchanging complete scanner trajectories between participants with equal numbers of observations to preserve longitudinal structure. We report overall and balanced classification accuracy (mean recall across scanner classes), mean permutation-based chance-level accuracies, 95% permutation intervals for balanced accuracy, and empirical permutation-based *p*-values.

To further investigate scanner-related variability, we examined scanner-specific residual variances before and after harmonisation. For each IDP, linear mixed-effects models were fitted using restricted maximum likelihood to the unharmonised data, Long-BARTharm, and LongComBat. Models included fixed effects for baseline age, sex, diagnostic group, time, diagnosis-by-time interactions, and scanner configuration, together with participant-specific random intercepts and slopes for time. Residual variances were calculated for each scanner configuration, and their homogeneity was assessed using the Fligner–Killeen (FK) test (Conover et al., 1981) to evaluate residual heterogeneity associated with multiplicative scanner effects.

Preservation of longitudinal biological effects was assessed using linear mixed-effects models fitted separately to each IDP before and after harmonisation using restricted maximum likelihood. Models included fixed effects for age, sex, MMSE total score, diagnostic group, time (years since each participant’s first available scan), and diagnosis-by-time interactions, with participant-specific random intercepts and slopes for time. With cognitively normal (CN) participants as the reference group, the time coefficient represented their estimated annual change, expressed in microns per year for cortical-thickness IDPs. Statistical significance was assessed using Kenward–Roger (KR) model comparisons (Kenward & Roger, 1997) between the full model and a reduced model excluding the main effect of time. Time coefficients and corresponding *p*-values were compared across unharmonised and harmonised IDPs, while the spatial distribution of Bonferroni-adjusted *p*-values across cortical regions was visualised using the fsbrain package (Schaefer & Ecker, 2020).

#### 2.4.3 Posterior predictive

Posterior predictive analysis was performed using both the cross-sectional and longitudinal models. The cross-sectional dataset was split into training and test sets using a fixed random seed, targeting a 10% test proportion. Test-set selection was restricted to scanner configurations with at least 15 observations, yielding 323 (90.2%) training and 35 (9.8%) test participants. The training set included 33 scanner configurations, six of which were represented in the test set.

To assess the preservation of age-related associations, age-effect estimates were compared across unharmonised data, ComBat, and BARTharm using full-data and posterior predictive harmonisation. ComBat was implemented using EB estimation. Comparisons included the combined harmonised training and test sets against the fully harmonised dataset, and the posterior predictively harmonised test set against the corresponding full-data harmonised values, with unharmonised and ComBat data similarly restricted to test participants. For each cortical-thickness IDP, ordinary least-squares regression models were fitted with age, sex, diagnosis, and MMSE score as covariates. Age-effect coefficients and corresponding *p*-values were obtained, with significance assessed at nominal and Bonferroni-corrected thresholds.

For the longitudinal posterior predictive analysis, participants with more than two longitudinal observations were selected, resulting in 351 eligible participants. The final observation from each eligible participant was assigned to the test set, yielding 351 held-out observations (20.6% of the total). The remaining 1,356 observations (79.4%) formed the training set.

To assess the preservation of longitudinal associations in cortical-thickness IDPs, time-effect coefficients and corresponding *p*-values were compared between the combined harmonised training and test sets and the fully harmonised dataset across unharmonised, LongComBat, and Long-BARTharm data, using the modelling and statistical testing procedures described in Section 2.4.2. Agreement between posterior predictive and full-data Long-BARTharm harmonisation was further evaluated by comparing test-set IDPs with their corresponding full-data harmonised values using scatter plots with identity lines.

#### 2.4.4 Uncertainty weighting

Posterior uncertainty in BARTharm harmonisation was examined across cortical-volume IDPs in both cross-sectional and longitudinal settings by comparing its distributions across regions of interest (ROIs). For visualisation, posterior variances were normalised by the squared standard deviation of the corresponding ROI. Posterior uncertainty was further examined at the individual-subject level for cortical-volume IDPs in both cross-sectional BARTharm and Long-BARTharm. Posterior variance was plotted for each subject to visualise the variability in uncertainty associated with individual harmonised estimates. To evaluate the influence of uncertainty weighting on the preservation of age and time-related associations, the regression models described in Sections 2.4.3 and 2.4.2, respectively, were fitted to BARTharm and Long-BARTharm-harmonised cortical-thickness IDPs using both unweighted and inverse-variance-weighted regression. Age and time-effect coefficients and corresponding *p*-values were compared with those obtained using ComBat in the cross-sectional setting and LongComBat in the longitudinal setting.

## 3 Results

### 3.1 Simulation studies

Table 1 summarises performance across the four fully synthetic scenarios. Long-BARTharm outperforms LongComBat on all four metrics in every scenario. Across 1,000 realisations per scenario, mean harmonisation RMSE ranges from 0.266 to 0.325 for Long-BARTharm and from 0.655 to 0.714 for LongComBat. These lower RMSE values show that Long-BARTharm recovers the known scanner-free outcomes more accurately than LongComBat in all four scenarios. The harmonised values correlate more strongly with the known scanner-free outcomes for Long-BARTharm (mean BIO Corr 0.992–0.995) than for LongComBat (0.888–0.904). Mean IQMs Corr values are lower for Long-BARTharm (0.031–0.040) than for LongComBat (0.325–0.379), indicating less remaining association with the simulated scanner component. This finding is consistent with scan-level IQMs capturing variation between scans acquired on the same scanner, which scanner identifiers alone cannot distinguish. Long-BARTharm also estimates the scanner-specific residual scale multipliers more accurately, with mean *δ* RMSE of 0.066–0.085 versus 0.303–0.312 for LongComBat.

**Table 1:** Mean performance across 1,000 fully synthetic datasets per scenario. BARTharm and ComBat denote Long-BARTharm and LongComBat, respectively. Bold values indicate the better result for each metric within a scenario.

| τ (BIO) | μ (IQMs) | RMSE |  | BIO Corr |  | IQMs Corr |  | δ RMSE |  |
|---|---|---|---|---|---|---|---|---|---|
|  |  | BARTharm | ComBat | BARTharm | ComBat | BARTharm | ComBat | BARTharm | ComBat |
| Linear | Linear | **0.266** | 0.709 | **0.995** | 0.890 | **0.032** | 0.379 | **0.066** | 0.304 |
| Linear | Non-linear | **0.325** | 0.655 | **0.992** | 0.904 | **0.040** | 0.325 | **0.085** | 0.312 |
| Non-linear | Linear | **0.299** | 0.714 | **0.994** | 0.888 | **0.031** | 0.379 | **0.066** | 0.303 |
| Non-linear | Non-linear | **0.320** | 0.661 | **0.992** | 0.902 | **0.039** | 0.325 | **0.085** | 0.312 |

Table 2 summarises performance across the four semi-synthetic scenarios based on ADNI. As in the fully synthetic simulations, Long-BARTharm achieves lower mean harmonisation RMSE, higher mean BIO Corr and lower mean IQMs Corr than LongComBat in all four scenarios. Mean harmonisation RMSE ranges from 0.355 to 0.386 for Long-BARTharm and from 0.774 to 0.810 for LongComBat, indicating more accurate recovery of the known scanner-free outcomes. The harmonised values also correlate more strongly with these outcomes for Long-BARTharm (mean BIO Corr 0.984–0.985) than for LongComBat (0.842–0.861). Mean IQMs Corr values are lower for Long-BARTharm (0.041–0.076) than for LongComBat (0.286–0.337), indicating less remaining association with the simulated scanner component.

**Table 2:** Mean performance across 1,000 semi-synthetic ADNI datasets per scenario. BARTharm and ComBat denote heteroskedastic Long-BARTharm and LongComBat, respectively. Bold values indicate the better result for each metric within a scenario.

| τ (BIO) | μ (IQMs) | RMSE |  | BIO Corr |  | IQMs Corr |  | δ RMSE |  |
|---|---|---|---|---|---|---|---|---|---|
|  |  | BARTharm | ComBat | BARTharm | ComBat | BARTharm | ComBat | BARTharm | ComBat |
| Linear | Linear | **0.366** | 0.804 | **0.985** | 0.848 | **0.051** | 0.337 | 0.528 | **0.417** |
| Linear | Non-linear | **0.386** | 0.774 | **0.984** | 0.861 | **0.041** | 0.307 | 0.669 | **0.633** |
| Non-linear | Linear | **0.355** | 0.810 | **0.985** | 0.842 | **0.076** | 0.316 | 0.528 | **0.417** |
| Non-linear | Non-linear | **0.373** | 0.781 | **0.984** | 0.854 | **0.063** | 0.286 | 0.665 | **0.632** |

Long-BARTharm estimates the scanner-specific residual scale multipliers more accurately in the fully synthetic simulations, whereas LongComBat achieves lower mean *δ* RMSE in all four semi-synthetic scenarios (0.417–0.633 for LongComBat versus 0.528–0.669 for Long-BARTharm). However, this advantage for LongComBat is concentrated among the 14 scanners with fewer than 10 scans each. For scanners with at least 20 scans, Long-BARTharm achieves lower mean *δ* RMSE in all four scenarios.

### 3.2 Real-data

#### 3.2.1 Longitudinal

We illustrate the harmonisation analysis using longitudinal neuroimaging data from the Alzheimer’s Disease Neuroimaging Initiative (ADNI). The analysis began with an assessment of IQMs to identify those potentially associated with biological and clinical characteristics, followed by the evaluation of harmonisation methods. Multivariable regression analyses revealed variation in the proportion of IQM variability explained by baseline biological and clinical characteristics (Table 3). The highest R² values were observed for the residual partial volume effect (RPVE) metrics (0.267-0.387), followed by the intracranial volume fraction (ICVS) metrics (0.205-0.242). In contrast, the remaining IQMs exhibited lower R² values (0.032-0.166). Based on these findings and their potential sensitivity to brain anatomy and tissue composition, the RPVE and ICVS metrics were excluded to reduce the risk of removing biologically relevant variation during harmonisation. The remaining 20 IQMs were retained for harmonisation.

**Table 3:** Coefficients of determination (*R*^2^) from multivariable linear regression models assessing associations between candidate IQMs and baseline biological and clinical characteristics. IQMs are ordered by decreasing *R*^2^. Asterisks indicate the six RPVE and ICVS metrics excluded from subsequent harmonisation analyses. IQM: image quality metric; RPVE: residual partial volume effect; ICVS: intracranial volume fraction

| **IQM** | *R*<sup>2</sup> | **IQM** | *R*<sup>2</sup> |
|---|---|---|---|
| rpve_wm<sup>*</sup> | 0.3869 | fwhm_x | 0.0885 |
| rpve_gm<sup>*</sup> | 0.3639 | snrd_wm | 0.0788 |
| rpve_csf<sup>*</sup> | 0.2673 | inu_range | 0.0785 |
| icvs_csf<sup>*</sup> | 0.2420 | fwhm_avg | 0.0785 |
| icvs_gm<sup>*</sup> | 0.2342 | fwhm_z | 0.0772 |
| icvs_wm<sup>*</sup> | 0.2054 | snrd_total | 0.0753 |
| cnr | 0.1662 | snrd_csf | 0.0720 |
| snr_wm | 0.1510 | snrd_gm | 0.0719 |
| cjv | 0.1262 | fwhm_y | 0.0701 |
| efc | 0.1116 | qi_2 | 0.0631 |
| snr_gm | 0.1015 | fber | 0.0566 |
| snr_csf | 0.0961 | inu_med | 0.0514 |
| snr_total | 0.0895 | qi_1 | 0.0322 |

**Table 4:**
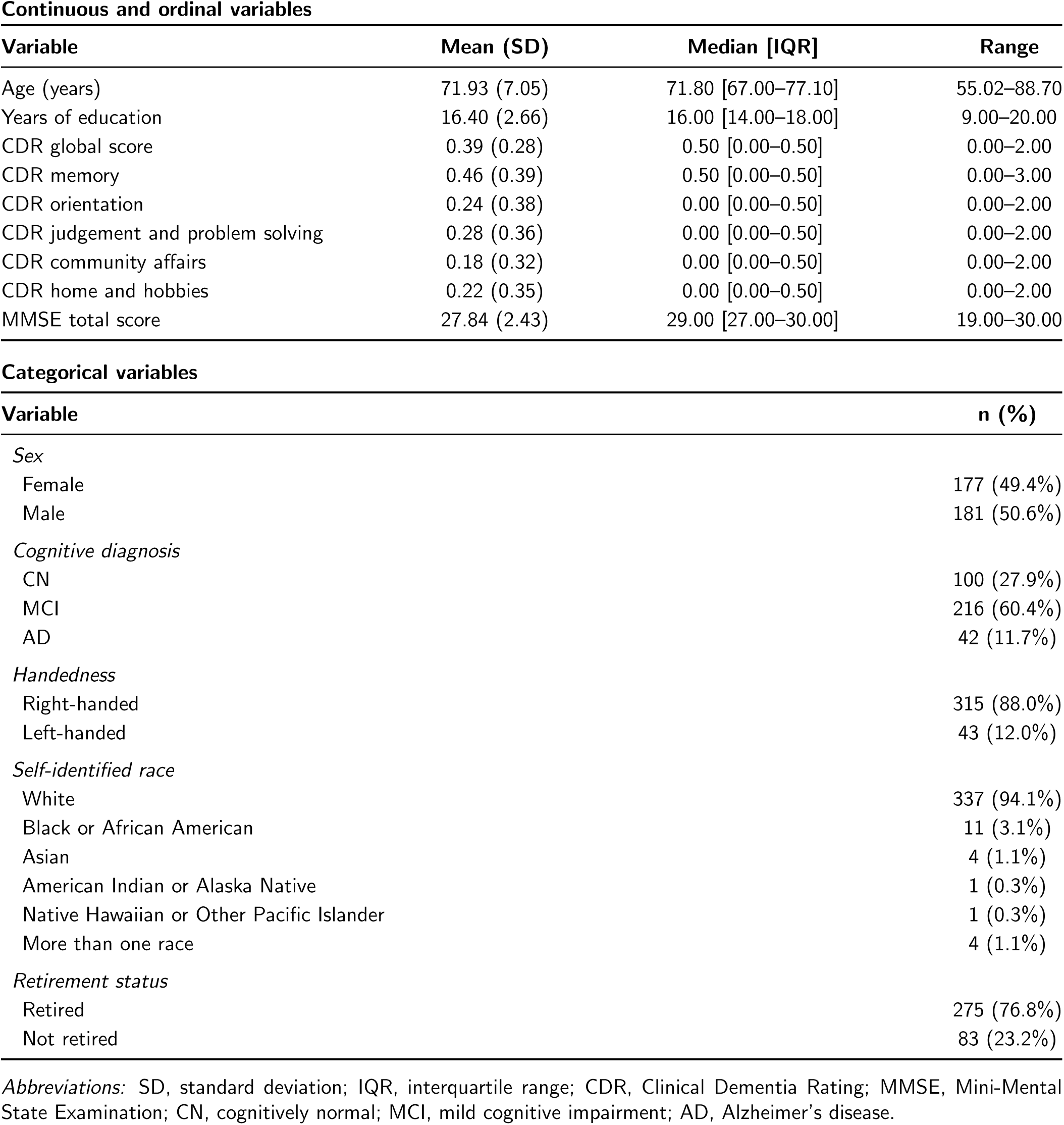
Baseline demographic and clinical characteristics of the ADNI cohort (*N* = 358). Continuous and ordinal variables are summarised as mean (standard deviation), median [interquartile range], and range. Categorical variables are reported as counts and percentages. All characteristics correspond to baseline measurements.

| **Continuous and ordinal variables** |  |  |  |
|---|---|---|---|
| **Variable** | **Mean (SD)** | **Median [IQR]** | **Range** |
| Age (years) | 71.93 (7.05) | 71.80 [67.00–77.10] | 55.02–88.70 |
| Years of education | 16.40 (2.66) | 16.00 [14.00–18.00] | 9.00–20.00 |
| CDR global score | 0.39 (0.28) | 0.50 [0.00–0.50] | 0.00–2.00 |
| CDR memory | 0.46 (0.39) | 0.50 [0.00–0.50] | 0.00–3.00 |
| CDR orientation | 0.24 (0.38) | 0.00 [0.00–0.50] | 0.00–2.00 |
| CDR judgement and problem solving | 0.28 (0.36) | 0.00 [0.00–0.50] | 0.00–2.00 |
| CDR community affairs | 0.18 (0.32) | 0.00 [0.00–0.50] | 0.00–2.00 |
| CDR home and hobbies | 0.22 (0.35) | 0.00 [0.00–0.50] | 0.00–2.00 |
| MMSE total score | 27.84 (2.43) | 29.00 [27.00–30.00] | 19.00–30.00 |
| **Categorical variables** |  |  |  |
| **Variable** |  |  | **n (%)** |
| *Sex* |  |  |  |
| Female |  |  | 177 (49.4%) |
| Male |  |  | 181 (50.6%) |
| *Cognitive diagnosis* |  |  |  |
| CN |  |  | 100 (27.9%) |
| MCI |  |  | 216 (60.4%) |
| AD |  |  | 42 (11.7%) |
| *Handedness* |  |  |  |
| Right-handed |  |  | 315 (88.0%) |
| Left-handed |  |  | 43 (12.0%) |
| *Self-identified race* |  |  |  |
| White |  |  | 337 (94.1%) |
| Black or African American |  |  | 11 (3.1%) |
| Asian |  |  | 4 (1.1%) |
| American Indian or Alaska Native |  |  | 1 (0.3%) |
| Native Hawaiian or Other Pacific Islander |  |  | 1 (0.3%) |
| More than one race |  |  | 4 (1.1%) |
| *Retirement status* |  |  |  |
| Retired |  |  | 275 (76.8%) |
| Not retired |  |  | 83 (23.2%) |

*Abbreviations:* SD, standard deviation; IQR, interquartile range; CDR, Clinical Dementia Rating; MMSE, Mini-Mental State Examination; CN, cognitively normal; MCI, mild cognitive impairment; AD, Alzheimer’s disease.

**Table 5:**
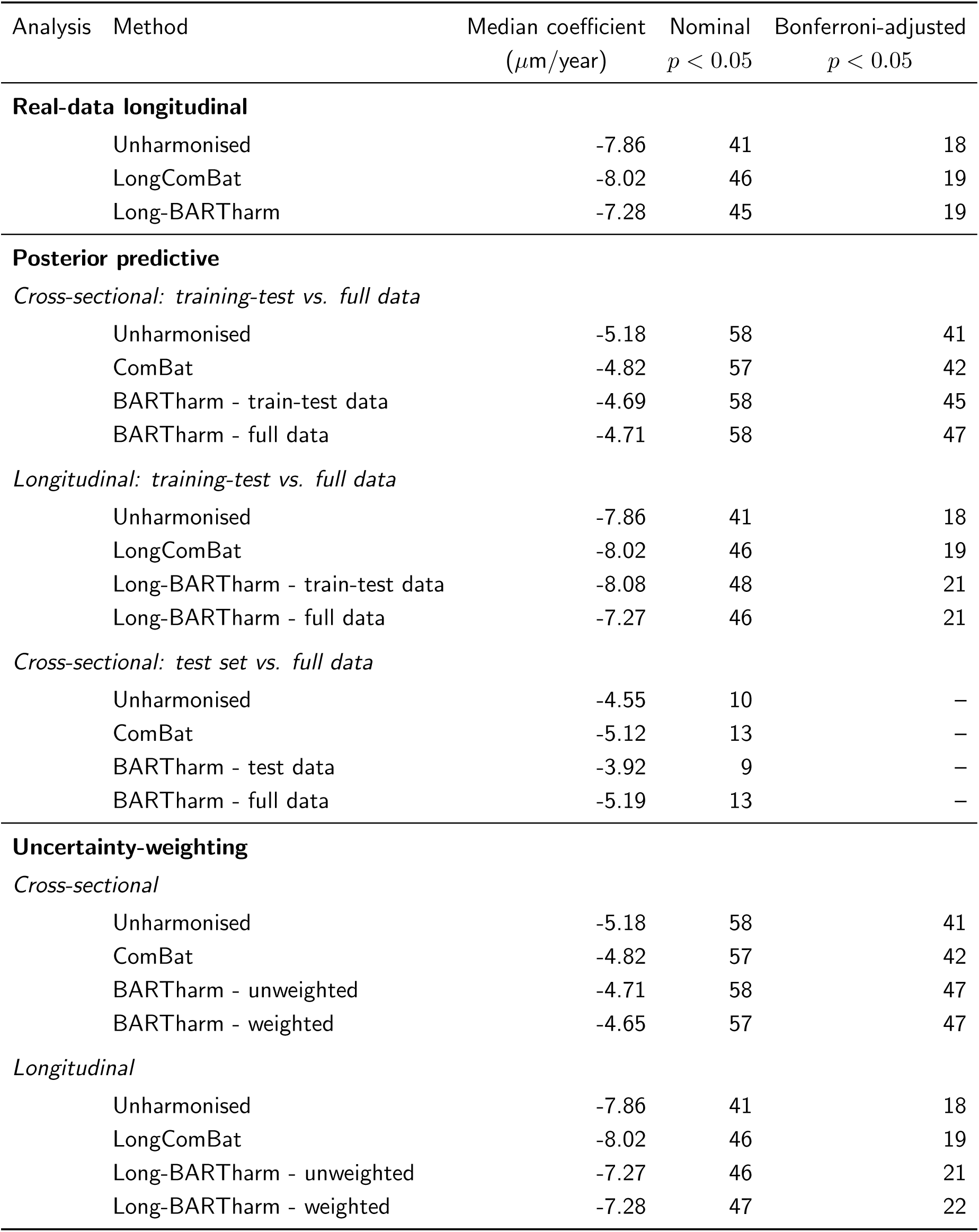
Summary of median cross-sectional age-effect and longitudinal time-effect coefficients across harmonisation approaches, together with the number of significant cortical-thickness IDPs at nominal and Bonferroni-adjusted significance thresholds.

| Analysis | Method | Median coefficient | Nominal | Bonferroni-adjusted |
|---|---|---|---|---|
|  |  | (*μ*m/year) | *p* < 0.05 | *p* < 0.05 |
| **Real-data longitudinal** |  |  |  |  |
|  | Unharmonised | -7.86 | 41 | 18 |
|  | LongComBat | -8.02 | 46 | 19 |
|  | Long-BARTharm | -7.28 | 45 | 19 |
| **Posterior predictive** |  |  |  |  |
| *Cross-sectional: training-test vs. full data* |  |  |  |  |
|  | Unharmonised | -5.18 | 58 | 41 |
|  | ComBat | -4.82 | 57 | 42 |
|  | BARTharm - train-test data | -4.69 | 58 | 45 |
|  | BARTharm - full data | -4.71 | 58 | 47 |
| *Longitudinal: training-test vs. full data* |  |  |  |  |
|  | Unharmonised | -7.86 | 41 | 18 |
|  | LongComBat | -8.02 | 46 | 19 |
|  | Long-BARTharm - train-test data | -8.08 | 48 | 21 |
|  | Long-BARTharm - full data | -7.27 | 46 | 21 |
| *Cross-sectional: test set vs. full data* |  |  |  |  |
|  | Unharmonised | -4.55 | 10 | – |
|  | ComBat | -5.12 | 13 | – |
|  | BARTharm - test data | -3.92 | 9 | – |
|  | BARTharm - full data | -5.19 | 13 | – |
| **Uncertainty-weighting** |  |  |  |  |
| *Cross-sectional* |  |  |  |  |
|  | Unharmonised | -5.18 | 58 | 41 |
|  | ComBat | -4.82 | 57 | 42 |
|  | BARTharm - unweighted | -4.71 | 58 | 47 |
|  | BARTharm - weighted | -4.65 | 57 | 47 |
| *Longitudinal* |  |  |  |  |
|  | Unharmonised | -7.86 | 41 | 18 |
|  | LongComBat | -8.02 | 46 | 19 |
|  | Long-BARTharm - unweighted | -7.27 | 46 | 21 |
|  | Long-BARTharm - weighted | -7.28 | 47 | 22 |

**Table 6:** Estimated longitudinal time-effect coefficients for cortical-thickness features across three harmonisation methods, separately for the left and right hemispheres. Values are expressed in *µ*m/year.

|  | **Left hemisphere** |  |  | **Right hemisphere** |  |  |
|---|---|---|---|---|---|---|
| Features | Unharm. | Long-BART | LongComBat | Unharm. | Long-BART | LongComBat |
| Banks Superior Temporal Sulcus | -10.62 | -7.94 | -9.84 | -8.97 | -8.96 | -9.14 |
| Caudal Anterior Cingulate | -4.17 | -3.41 | -3.80 | -7.84 | -8.17 | -7.92 |
| Caudal Middle Frontal | -6.85 | -6.46 | -7.20 | -4.49 | -4.75 | -4.33 |
| Cuneus | -2.83 | -2.64 | -2.80 | -4.07 | -4.15 | -4.04 |
| Entorhinal | -22.32 | -22.43 | -23.88 | -25.29 | -24.11 | -23.16 |
| Frontal Pole | -5.50 | -4.74 | -4.90 | -0.22 | -0.77 | -0.25 |
| Fusiform | -17.45 | -16.47 | -17.48 | -15.41 | -13.55 | -15.11 |
| Inferior Parietal | -9.00 | -8.82 | -8.43 | -7.40 | -6.74 | -7.31 |
| Inferior Temporal | -18.98 | -17.59 | -18.43 | -15.72 | -15.27 | -15.46 |
| Isthmus Cingulate | -14.33 | -13.04 | -14.37 | -12.89 | -11.82 | -12.97 |
| Lateral Occipital | -9.76 | -7.98 | -9.37 | -9.30 | -8.36 | -9.23 |
| Lateral Orbitofrontal | -8.91 | -6.66 | -8.58 | -6.54 | -4.71 | -6.53 |
| Lingual | -8.81 | -7.77 | -8.88 | -7.68 | -6.71 | -7.71 |
| Medial Orbitofrontal | -10.85 | -11.53 | -11.06 | -4.55 | -4.06 | -4.81 |
| Middle Temporal | -14.47 | -13.24 | -13.81 | -13.44 | -14.08 | -13.64 |
| Paracentral | -5.62 | -5.46 | -6.08 | -4.88 | -5.64 | -6.20 |
| Parahippocampal | -18.06 | -16.67 | -18.40 | -16.79 | -16.12 | -17.75 |
| Pars Opercularis | -7.23 | -6.51 | -6.86 | -5.52 | -5.61 | -5.08 |
| Pars Orbitalis | -4.05 | -1.36 | -3.26 | -3.08 | -1.43 | -2.86 |
| Pars Triangularis | -6.79 | -5.98 | -6.46 | -7.59 | -8.09 | -8.13 |
| Pericalcarine | -2.03 | -2.41 | -2.36 | -0.65 | -1.56 | -1.17 |
| Postcentral | -4.76 | -4.23 | -5.07 | -4.58 | -4.69 | -4.72 |
| Posterior Cingulate | -10.24 | -9.04 | -10.29 | -10.44 | -10.30 | -10.41 |
| Precentral | -9.86 | -9.07 | -11.08 | -5.84 | -7.20 | -6.45 |
| Precuneus | -9.05 | -9.29 | -9.30 | -10.60 | -9.81 | -10.90 |
| Rostral Anterior Cingulate | -2.71 | -2.65 | -2.90 | 2.58 | 1.87 | 2.72 |
| Rostral Middle Frontal | -3.32 | -2.76 | -3.13 | 1.30 | 0.99 | 1.75 |
| Superior Frontal | -7.88 | -7.29 | -7.14 | -4.89 | -4.62 | -4.65 |
| Superior Parietal | -6.30 | -5.76 | -6.14 | -6.52 | -6.25 | -6.44 |
| Superior Temporal | -15.78 | -14.09 | -15.09 | -13.59 | -14.42 | -14.26 |
| Supramarginal | -10.17 | -9.40 | -9.93 | -8.08 | -9.65 | -8.38 |
| Temporal Pole | -17.58 | -15.95 | -16.65 | -28.34 | -27.31 | -28.37 |
| Transverse Temporal | -6.69 | -5.19 | -6.90 | -8.82 | -7.27 | -8.57 |

**Table 7:** Kenward–Roger *p*-values for longitudinal time-effect coefficients of cortical-thickness features across three harmonisation methods, separately for the left and right hemispheres.

|  | **Left hemisphere** |  |  | **Right hemisphere** |  |  |
|---|---|---|---|---|---|---|
| Features | Unharm. | Long-BART | LongComBat | Unharm. | Long-BART | LongComBat |
| Banks Superior Temporal Sulcus | 0.006 | 0.024 | 0.009 | 0.011 | 0.007 | 0.007 |
| Caudal Anterior Cingulate | 0.324 | 0.392 | 0.372 | 0.098 | 0.061 | 0.092 |
| Caudal Middle Frontal | 0.069 | 0.044 | 0.049 | 0.222 | 0.115 | 0.215 |
| Cuneus | 0.173 | 0.158 | 0.176 | 0.048 | 0.024 | 0.045 |
| Entorhinal | 8.57e-4 | 3.60e-4 | 2.65e-4 | 5.10e-4 | 3.79e-4 | 9.85e-4 |
| Frontal Pole | 0.338 | 0.358 | 0.385 | 0.964 | 0.862 | 0.959 |
| Fusiform | 9.61e-6 | 1.93e-6 | 3.67e-6 | 2.39e-4 | 1.60e-4 | 1.49e-4 |
| Inferior Parietal | 0.008 | 0.001 | 0.008 | 0.018 | 0.013 | 0.014 |
| Inferior Temporal | 1.03e-5 | 8.95e-6 | 8.45e-6 | 1.19e-4 | 2.43e-5 | 5.37e-5 |
| Isthmus Cingulate | 4.32e-4 | 3.90e-4 | 2.26e-4 | 8.50e-4 | 9.52e-4 | 3.08e-4 |
| Lateral Occipital | 7.02e-4 | 0.003 | 0.001 | 0.002 | 0.002 | 0.001 |
| Lateral Orbitofrontal | 0.005 | 0.025 | 0.005 | 0.060 | 0.119 | 0.048 |
| Lingual | 2.23e-5 | 2.03e-5 | 1.24e-5 | 3.77e-4 | 3.48e-4 | 2.53e-4 |
| Medial Orbitofrontal | 7.40e-4 | 1.16e-4 | 4.15e-4 | 0.181 | 0.188 | 0.143 |
| Middle Temporal | 2.43e-4 | 1.29e-4 | 2.73e-4 | 4.49e-4 | 2.50e-5 | 2.04e-4 |
| Paracentral | 0.102 | 0.060 | 0.073 | 0.116 | 0.040 | 0.049 |
| Parahippocampal | 3.62e-5 | 2.58e-5 | 1.55e-5 | 3.96e-4 | 1.12e-4 | 7.08e-5 |
| Pars Opercularis | 0.024 | 0.020 | 0.028 | 0.094 | 0.052 | 0.110 |
| Pars Orbitalis | 0.324 | 0.703 | 0.402 | 0.409 | 0.671 | 0.426 |
| Pars Triangularis | 0.045 | 0.038 | 0.050 | 0.016 | 0.004 | 0.008 |
| Pericalcarine | 0.347 | 0.179 | 0.263 | 0.769 | 0.412 | 0.593 |
| Postcentral | 0.050 | 0.039 | 0.036 | 0.057 | 0.018 | 0.043 |
| Posterior Cingulate | 0.002 | 0.003 | 0.002 | 0.005 | 0.003 | 0.003 |
| Precentral | 0.007 | 9.54e-4 | 0.003 | 0.098 | 0.016 | 0.066 |
| Precuneus | 0.002 | 1.86e-4 | 7.53e-4 | 8.21e-5 | 1.96e-5 | 2.62e-5 |
| Rostral Anterior Cingulate | 0.498 | 0.473 | 0.457 | 0.545 | 0.626 | 0.514 |
| Rostral Middle Frontal | 0.287 | 0.274 | 0.297 | 0.658 | 0.675 | 0.535 |
| Superior Frontal | 0.023 | 0.009 | 0.028 | 0.122 | 0.061 | 0.124 |
| Superior Parietal | 0.032 | 0.019 | 0.027 | 0.020 | 0.006 | 0.014 |
| Superior Temporal | 1.94e-6 | 1.98e-7 | 1.82e-6 | 1.94e-5 | 3.22e-7 | 6.64e-6 |
| Supramarginal | 0.003 | 9.25e-4 | 0.002 | 0.016 | 7.71e-4 | 0.008 |
| Temporal Pole | 0.002 | 0.002 | 0.003 | 8.79e-5 | 2.79e-5 | 4.52e-5 |
| Transverse Temporal | 0.073 | 0.099 | 0.062 | 0.024 | 0.051 | 0.030 |

**Table 8:** Fligner–Killeen chi-squared statistics (*χ*^2^) and associated *p*-values for cortical thickness features across three harmonisation methods, shown side by side for the left and right hemispheres.

|  | Left hemisphere |  |  |  |  |  | Right hemisphere |  |  |  |  |  |
|---|---|---|---|---|---|---|---|---|---|---|---|---|
|  | Unharmonised |  | Long-BARTharm |  | LongComBat |  | Unharmonised |  | Long-BARTharm |  | LongComBat |  |
| Feature | χ<sup>2</sup> | *p*-value | χ<sup>2</sup> | *p*-value | χ<sup>2</sup> | *p*-value | χ<sup>2</sup> | *p*-value | χ<sup>2</sup> | *p*-value | χ<sup>2</sup> | *p*-value |
| Banks Superior Temporal Sulcus | 73.61 | 4.01 × 10<sup>−5</sup> | 11.21 | 1.000 | 19.88 | 0.953 | 67.19 | 2.69 × 10<sup>−4</sup> | 6.78 | 1.000 | 11.16 | 1.000 |
| Caudal Anterior Cingulate | 92.83 | 7.76 × 10<sup>−8</sup> | 15.29 | 0.995 | 25.22 | 0.797 | 141.46 | 1.03 × 10<sup>−15</sup> | 9.07 | 1.000 | 23.19 | 0.872 |
| Caudal Middle Frontal | 55.53 | 0.006 | 12.05 | 0.999 | 15.65 | 0.993 | 73.20 | 4.54 × 10<sup>−5</sup> | 11.61 | 1.000 | 11.98 | 0.999 |
| Cuneus | 36.29 | 0.275 | 7.31 | 1.000 | 10.91 | 1.000 | 47.91 | 0.035 | 6.44 | 1.000 | 9.40 | 1.000 |
| Entorhinal | 45.07 | 0.063 | 14.88 | 0.996 | 23.22 | 0.871 | 79.90 | 5.65 × 10<sup>−6</sup> | 15.41 | 0.994 | 18.20 | 0.976 |
| Frontal Pole | 87.83 | 4.21 × 10<sup>−7</sup> | 8.04 | 1.000 | 21.88 | 0.911 | 47.33 | 0.040 | 9.01 | 1.000 | 13.97 | 0.998 |
| Fusiform | 38.65 | 0.194 | 9.65 | 1.000 | 21.92 | 0.910 | 68.57 | 1.80 × 10<sup>−4</sup> | 16.86 | 0.987 | 27.05 | 0.715 |
| Inferior Parietal | 71.11 | 8.53 × 10<sup>−5</sup> | 11.49 | 1.000 | 12.76 | 0.999 | 64.97 | 5.03 × 10<sup>−4</sup> | 13.13 | 0.999 | 12.72 | 0.999 |
| Inferior Temporal | 48.45 | 0.031 | 11.03 | 1.000 | 18.56 | 0.972 | 94.73 | 4.02 × 10<sup>−8</sup> | 22.64 | 0.889 | 27.50 | 0.694 |
| Isthmus Cingulate | 68.47 | 1.86 × 10<sup>−4</sup> | 12.01 | 0.999 | 20.04 | 0.950 | 56.83 | 0.004 | 13.37 | 0.998 | 11.29 | 1.000 |
| Lateral Occipital | 50.09 | 0.022 | 13.15 | 0.999 | 19.71 | 0.956 | 65.26 | 4.64 × 10<sup>−4</sup> | 12.38 | 0.999 | 18.33 | 0.975 |
| Lateral Orbitofrontal | 50.15 | 0.022 | 9.21 | 1.000 | 18.36 | 0.974 | 64.30 | 6.08 × 10<sup>−4</sup> | 10.39 | 1.000 | 13.12 | 0.999 |
| Lingual | 34.71 | 0.340 | 9.95 | 1.000 | 13.22 | 0.999 | 54.14 | 0.009 | 9.61 | 1.000 | 13.87 | 0.998 |
| Medial Orbitofrontal | 67.88 | 2.20 × 10<sup>−4</sup> | 13.00 | 0.999 | 18.32 | 0.975 | 49.59 | 0.024 | 9.22 | 1.000 | 12.12 | 0.999 |
| Middle Temporal | 49.94 | 0.023 | 14.33 | 0.997 | 20.88 | 0.934 | 85.85 | 8.16 × 10<sup>−7</sup> | 8.49 | 1.000 | 20.81 | 0.936 |
| Paracentral | 83.36 | 1.85 × 10<sup>−6</sup> | 10.24 | 1.000 | 19.80 | 0.955 | 70.53 | 1.01 × 10<sup>−4</sup> | 12.82 | 0.999 | 15.94 | 0.992 |
| Parahippocampal | 60.96 | 0.002 | 11.17 | 1.000 | 23.18 | 0.872 | 48.28 | 0.032 | 12.72 | 0.999 | 19.55 | 0.959 |
| Pars Opercularis | 62.27 | 0.001 | 12.53 | 0.999 | 22.01 | 0.907 | 66.35 | 3.41 × 10<sup>−4</sup> | 12.58 | 0.999 | 9.56 | 1.000 |
| Pars Orbitalis | 94.64 | 4.16 × 10<sup>−8</sup> | 6.22 | 1.000 | 12.93 | 0.999 | 94.38 | 4.55 × 10<sup>−8</sup> | 7.01 | 1.000 | 15.07 | 0.995 |
| Pars Triangularis | 59.46 | 0.002 | 15.48 | 0.994 | 23.62 | 0.858 | 72.16 | 6.23 × 10<sup>−5</sup> | 5.57 | 1.000 | 10.04 | 1.000 |
| Pericalcarine | 81.19 | 3.73 × 10<sup>−6</sup> | 9.08 | 1.000 | 18.50 | 0.973 | 63.16 | 8.32 × 10<sup>−4</sup> | 8.64 | 1.000 | 13.04 | 0.999 |
| Postcentral | 47.81 | 0.036 | 18.46 | 0.973 | 13.45 | 0.998 | 67.68 | 2.33 × 10<sup>−4</sup> | 14.54 | 0.997 | 14.87 | 0.996 |
| Posterior Cingulate | 75.46 | 2.28 × 10<sup>−5</sup> | 24.29 | 0.834 | 32.79 | 0.428 | 72.87 | 5.02 × 10<sup>−5</sup> | 18.47 | 0.973 | 20.76 | 0.937 |
| Precentral | 71.47 | 7.66 × 10<sup>−5</sup> | 14.27 | 0.997 | 15.27 | 0.995 | 64.91 | 5.13 × 10<sup>−4</sup> | 15.11 | 0.995 | 18.64 | 0.971 |
| Precuneus | 67.67 | 2.34 × 10<sup>−4</sup> | 13.38 | 0.998 | 16.29 | 0.990 | 66.77 | 3.03 × 10<sup>−4</sup> | 15.67 | 0.993 | 14.04 | 0.998 |
| Rostral Anterior Cingulate | 52.27 | 0.013 | 15.40 | 0.994 | 17.78 | 0.980 | 82.16 | 2.73 × 10<sup>−6</sup> | 16.84 | 0.987 | 20.84 | 0.935 |
| Rostral Middle Frontal | 63.69 | 7.19 × 10<sup>−4</sup> | 13.76 | 0.998 | 15.41 | 0.994 | 140.09 | 1.76 × 10<sup>−15</sup> | 8.41 | 1.000 | 20.10 | 0.950 |
| Superior Frontal | 109.33 | 2.21 × 10<sup>−10</sup> | 25.68 | 0.778 | 29.56 | 0.591 | 155.87 | 3.19 × 10<sup>−18</sup> | 19.81 | 0.955 | 19.35 | 0.962 |
| Superior Parietal | 64.85 | 5.21 × 10<sup>−4</sup> | 10.14 | 1.000 | 14.99 | 0.995 | 48.69 | 0.030 | 4.68 | 1.000 | 5.89 | 1.000 |
| Superior Temporal | 99.54 | 7.47 × 10<sup>−9</sup> | 19.32 | 0.962 | 24.78 | 0.815 | 135.19 | 1.21 × 10<sup>−14</sup> | 21.28 | 0.925 | 25.03 | 0.805 |
| Supramarginal | 65.76 | 4.03 × 10<sup>−4</sup> | 12.41 | 0.999 | 17.61 | 0.981 | 58.49 | 0.003 | 8.23 | 1.000 | 11.52 | 1.000 |
| Temporal Pole | 76.57 | 1.61 × 10<sup>−5</sup> | 11.44 | 1.000 | 14.49 | 0.997 | 49.48 | 0.025 | 16.43 | 0.990 | 16.97 | 0.986 |
| Transverse Temporal | 44.13 | 0.075 | 14.55 | 0.997 | 13.91 | 0.998 | 76.28 | 1.76 × 10<sup>−5</sup> | 10.77 | 1.000 | 14.46 | 0.997 |

**Table 9:**
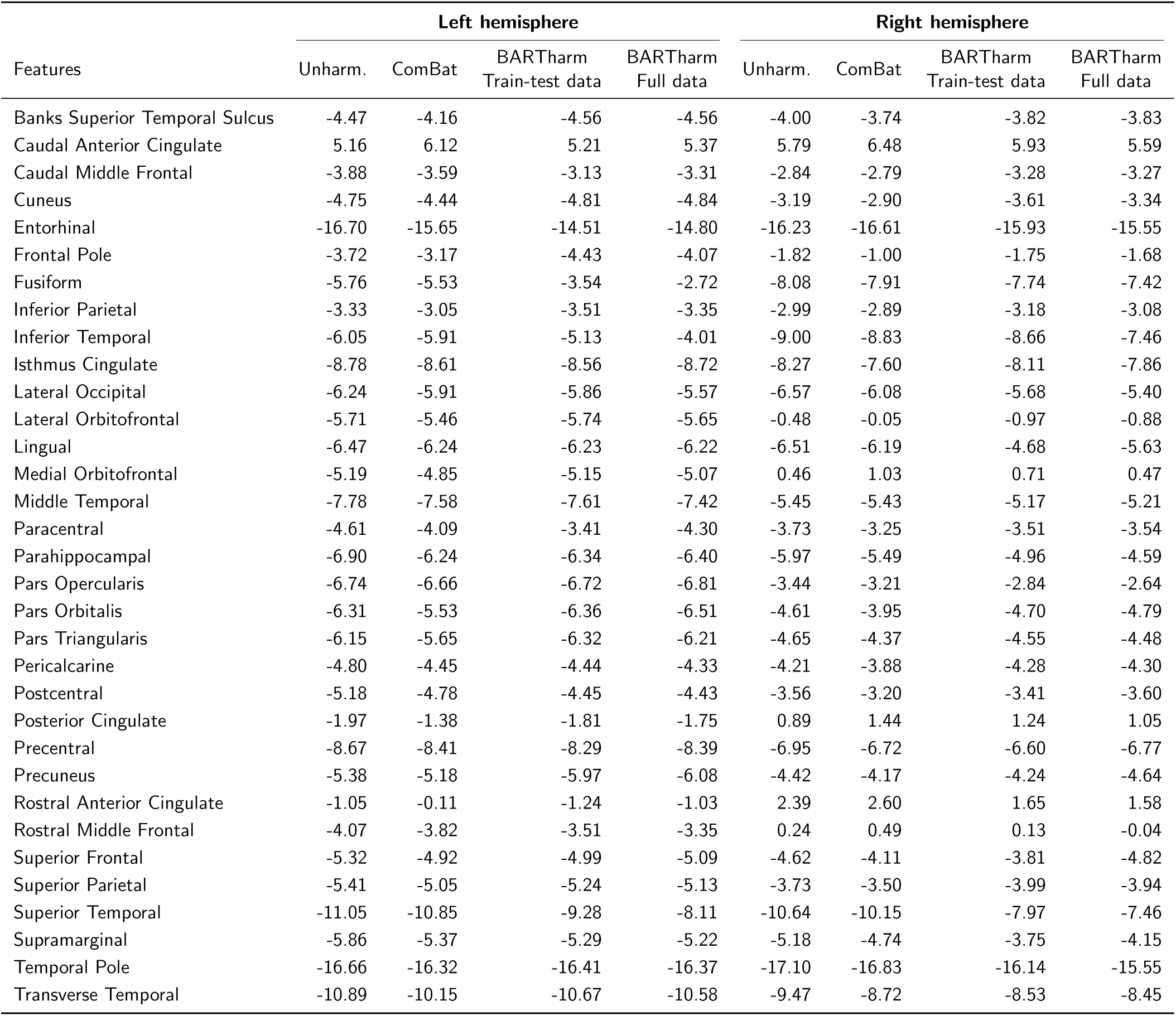
Estimated cross-sectional age-effect coefficients for cortical-thickness features across unharmonised data, ComBat, and BARTharm applied to the train-test and full datasets. Values are expressed in *µ*m/year.

**Table 10:** Nominal *p*-values for cross-sectional age-effect coefficients of cortical-thickness features across unharmonised data, ComBat, and BARTharm applied to the train-test and full datasets, separately for the left and right hemispheres.

|  | Left hemisphere |  |  |  | Right hemisphere |  |  |  |
|---|---|---|---|---|---|---|---|---|
| Features | Unharm. | ComBat | BARTharm<br>Train-test data | BARTharm<br>Full data | Unharm. | ComBat | BARTharm<br>Train-test data | BARTharm<br>Full data |
| Banks Superior Temporal Sulcus | 0.002 | 0.002 | 5.37 × 10<sup>−4</sup> | 5.73 × 10<sup>−4</sup> | 0.002 | 0.001 | 0.002 | 0.002 |
| Caudal Anterior Cingulate | 0.035 | 0.007 | 0.031 | 0.026 | 0.015 | 0.004 | 0.017 | 0.019 |
| Caudal Middle Frontal | 9.40 × 10<sup>−4</sup> | 8.70 × 10<sup>−4</sup> | 0.003 | 0.002 | 0.019 | 0.011 | 0.004 | 0.004 |
| Cuneus | 1.67 × 10<sup>−5</sup> | 3.44 × 10<sup>−6</sup> | 5.01 × 10<sup>−7</sup> | 3.75 × 10<sup>−7</sup> | 0.005 | 0.002 | 9.09 × 10<sup>−7</sup> | 7.00 × 10<sup>−5</sup> |
| Entorhinal | 7.42 × 10<sup>−7</sup> | 3.94 × 10<sup>−7</sup> | 2.01 × 10<sup>−6</sup> | 1.55 × 10<sup>−6</sup> | 9.65 × 10<sup>−6</sup> | 8.47 × 10<sup>−7</sup> | 3.02 × 10<sup>−6</sup> | 5.09 × 10<sup>−6</sup> |
| Frontal Pole | 0.047 | 0.068 | 0.019 | 0.027 | 0.339 | 0.576 | 0.362 | 0.380 |
| Fusiform | 1.12 × 10<sup>−4</sup> | 3.26 × 10<sup>−5</sup> | 2.75 × 10<sup>−4</sup> | 0.002 | 5.30 × 10<sup>−8</sup> | 6.76 × 10<sup>−9</sup> | 1.30 × 10<sup>−8</sup> | 3.21 × 10<sup>−8</sup> |
| Inferior Parietal | 0.004 | 0.003 | 3.79 × 10<sup>−4</sup> | 3.33 × 10<sup>−4</sup> | 0.013 | 0.006 | 0.001 | 5.96 × 10<sup>−4</sup> |
| Inferior Temporal | 5.51 × 10<sup>−5</sup> | 1.48 × 10<sup>−5</sup> | 2.17 × 10<sup>−6</sup> | 4.02 × 10<sup>−6</sup> | 8.47 × 10<sup>−10</sup> | 2.03 × 10<sup>−11</sup> | 1.41 × 10<sup>−11</sup> | 9.34 × 10<sup>−12</sup> |
| Isthmus Cingulate | 2.60 × 10<sup>−6</sup> | 4.37 × 10<sup>−7</sup> | 1.41 × 10<sup>−6</sup> | 5.09 × 10<sup>−7</sup> | 5.17 × 10<sup>−6</sup> | 5.15 × 10<sup>−6</sup> | 2.91 × 10<sup>−6</sup> | 6.61 × 10<sup>−6</sup> |
| Lateral Occipital | 6.91 × 10<sup>−8</sup> | 7.37 × 10<sup>−9</sup> | 3.51 × 10<sup>−9</sup> | 2.69 × 10<sup>−11</sup> | 9.71 × 10<sup>−8</sup> | 1.71 × 10<sup>−8</sup> | 1.12 × 10<sup>−8</sup> | 2.57 × 10<sup>−10</sup> |
| Lateral Orbitofrontal | 8.33 × 10<sup>−6</sup> | 2.97 × 10<sup>−6</sup> | 2.25 × 10<sup>−6</sup> | 2.49 × 10<sup>−6</sup> | 0.705 | 0.969 | 0.440 | 0.480 |
| Lingual | 9.95 × 10<sup>−10</sup> | 1.02 × 10<sup>−10</sup> | 3.65 × 10<sup>−10</sup> | 2.30 × 10<sup>−10</sup> | 9.17 × 10<sup>−9</sup> | 1.26 × 10<sup>−9</sup> | 1.04 × 10<sup>−10</sup> | 5.18 × 10<sup>−10</sup> |
| Medial Orbitofrontal | 1.13 × 10<sup>−5</sup> | 3.86 × 10<sup>−6</sup> | 3.19 × 10<sup>−6</sup> | 4.43 × 10<sup>−6</sup> | 0.748 | 0.422 | 0.619 | 0.741 |
| Middle Temporal | 4.14 × 10<sup>−8</sup> | 4.47 × 10<sup>−9</sup> | 2.87 × 10<sup>−9</sup> | 2.44 × 10<sup>−9</sup> | 3.48 × 10<sup>−5</sup> | 4.35 × 10<sup>−6</sup> | 5.89 × 10<sup>−5</sup> | 1.05 × 10<sup>−5</sup> |
| Paracentral | 3.37 × 10<sup>−4</sup> | 3.80 × 10<sup>−4</sup> | 9.13 × 10<sup>−4</sup> | 3.34 × 10<sup>−4</sup> | 0.005 | 0.004 | 0.002 | 5.70 × 10<sup>−4</sup> |
| Parahippocampal | 0.010 | 0.011 | 0.013 | 0.012 | 0.009 | 0.009 | 0.019 | 0.029 |
| Pars Opercularis | 6.68 × 10<sup>−10</sup> | 2.35 × 10<sup>−11</sup> | 1.78 × 10<sup>−11</sup> | 1.36 × 10<sup>−11</sup> | 0.003 | 0.002 | 0.008 | 0.013 |
| Pars Orbitalis | 2.58 × 10<sup>−4</sup> | 3.50 × 10<sup>−4</sup> | 7.40 × 10<sup>−5</sup> | 4.90 × 10<sup>−5</sup> | 0.007 | 0.010 | 0.006 | 0.005 |
| Pars Triangularis | 1.89 × 10<sup>−7</sup> | 1.98 × 10<sup>−7</sup> | 3.50 × 10<sup>−8</sup> | 2.37 × 10<sup>−8</sup> | 5.36 × 10<sup>−5</sup> | 3.15 × 10<sup>−5</sup> | 3.76 × 10<sup>−5</sup> | 4.05 × 10<sup>−5</sup> |
| Pericalcarine | 3.60 × 10<sup>−7</sup> | 1.65 × 10<sup>−8</sup> | 2.75 × 10<sup>−9</sup> | 1.87 × 10<sup>−8</sup> | 4.77 × 10<sup>−5</sup> | 5.59 × 10<sup>−6</sup> | 7.18 × 10<sup>−7</sup> | 3.21 × 10<sup>−7</sup> |
| Postcentral | 3.70 × 10<sup>−6</sup> | 4.79 × 10<sup>−7</sup> | 2.65 × 10<sup>−8</sup> | 4.25 × 10<sup>−7</sup> | 7.92 × 10<sup>−4</sup> | 7.48 × 10<sup>−4</sup> | 4.63 × 10<sup>−4</sup> | 3.26 × 10<sup>−4</sup> |
| Posterior Cingulate | 0.141 | 0.267 | 0.169 | 0.182 | 0.546 | 0.292 | 0.387 | 0.455 |
| Precentral | 6.84 × 10<sup>−12</sup> | 1.33 × 10<sup>−13</sup> | 1.40 × 10<sup>−12</sup> | 8.45 × 10<sup>−13</sup> | 6.10 × 10<sup>−8</sup> | 6.83 × 10<sup>−9</sup> | 6.50 × 10<sup>−9</sup> | 8.34 × 10<sup>−10</sup> |
| Precuneus | 9.05 × 10<sup>−6</sup> | 2.51 × 10<sup>−6</sup> | 1.54 × 10<sup>−7</sup> | 8.75 × 10<sup>−8</sup> | 1.31 × 10<sup>−4</sup> | 4.82 × 10<sup>−5</sup> | 2.15 × 10<sup>−6</sup> | 3.98 × 10<sup>−6</sup> |
| Rostral Anterior Cingulate | 0.609 | 0.955 | 0.552 | 0.620 | 0.242 | 0.163 | 0.406 | 0.416 |
| Rostral Middle Frontal | 1.50 × 10<sup>−5</sup> | 5.94 × 10<sup>−6</sup> | 6.15 × 10<sup>−7</sup> | 7.48 × 10<sup>−7</sup> | 0.791 | 0.565 | 0.882 | 0.959 |
| Superior Frontal | 1.90 × 10<sup>−6</sup> | 1.42 × 10<sup>−6</sup> | 1.27 × 10<sup>−6</sup> | 8.04 × 10<sup>−7</sup> | 4.07 × 10<sup>−5</sup> | 6.05 × 10<sup>−5</sup> | 4.14 × 10<sup>−5</sup> | 2.17 × 10<sup>−11</sup> |
| Superior Parietal | 1.48 × 10<sup>−5</sup> | 3.78 × 10<sup>−6</sup> | 7.05 × 10<sup>−7</sup> | 6.24 × 10<sup>−7</sup> | 0.002 | 8.11 × 10<sup>−4</sup> | 1.11 × 10<sup>−4</sup> | 1.08 × 10<sup>−4</sup> |
| Superior Temporal | 1.44 × 10<sup>−14</sup> | 3.50 × 10<sup>−16</sup> | 4.52 × 10<sup>−17</sup> | 4.87 × 10<sup>−15</sup> | 4.31 × 10<sup>−15</sup> | 4.45 × 10<sup>−17</sup> | 3.18 × 10<sup>−18</sup> | 1.97 × 10<sup>−20</sup> |
| Supramarginal | 1.82 × 10<sup>−7</sup> | 1.46 × 10<sup>−7</sup> | 3.63 × 10<sup>−7</sup> | 5.98 × 10<sup>−7</sup> | 4.47 × 10<sup>−6</sup> | 2.29 × 10<sup>−6</sup> | 8.29 × 10<sup>−8</sup> | 3.38 × 10<sup>−6</sup> |
| Temporal Pole | 9.11 × 10<sup>−9</sup> | 8.44 × 10<sup>−10</sup> | 3.98 × 10<sup>−10</sup> | 1.65 × 10<sup>−10</sup> | 1.18 × 10<sup>−8</sup> | 4.17 × 10<sup>−10</sup> | 2.70 × 10<sup>−9</sup> | 4.54 × 10<sup>−9</sup> |
| Transverse Temporal | 1.18 × 10<sup>−9</sup> | 1.61 × 10<sup>−10</sup> | 9.22 × 10<sup>−11</sup> | 1.29 × 10<sup>−10</sup> | 5.08 × 10<sup>−7</sup> | 2.80 × 10<sup>−7</sup> | 7.26 × 10<sup>−7</sup> | 1.50 × 10<sup>−6</sup> |

**Table 11:**
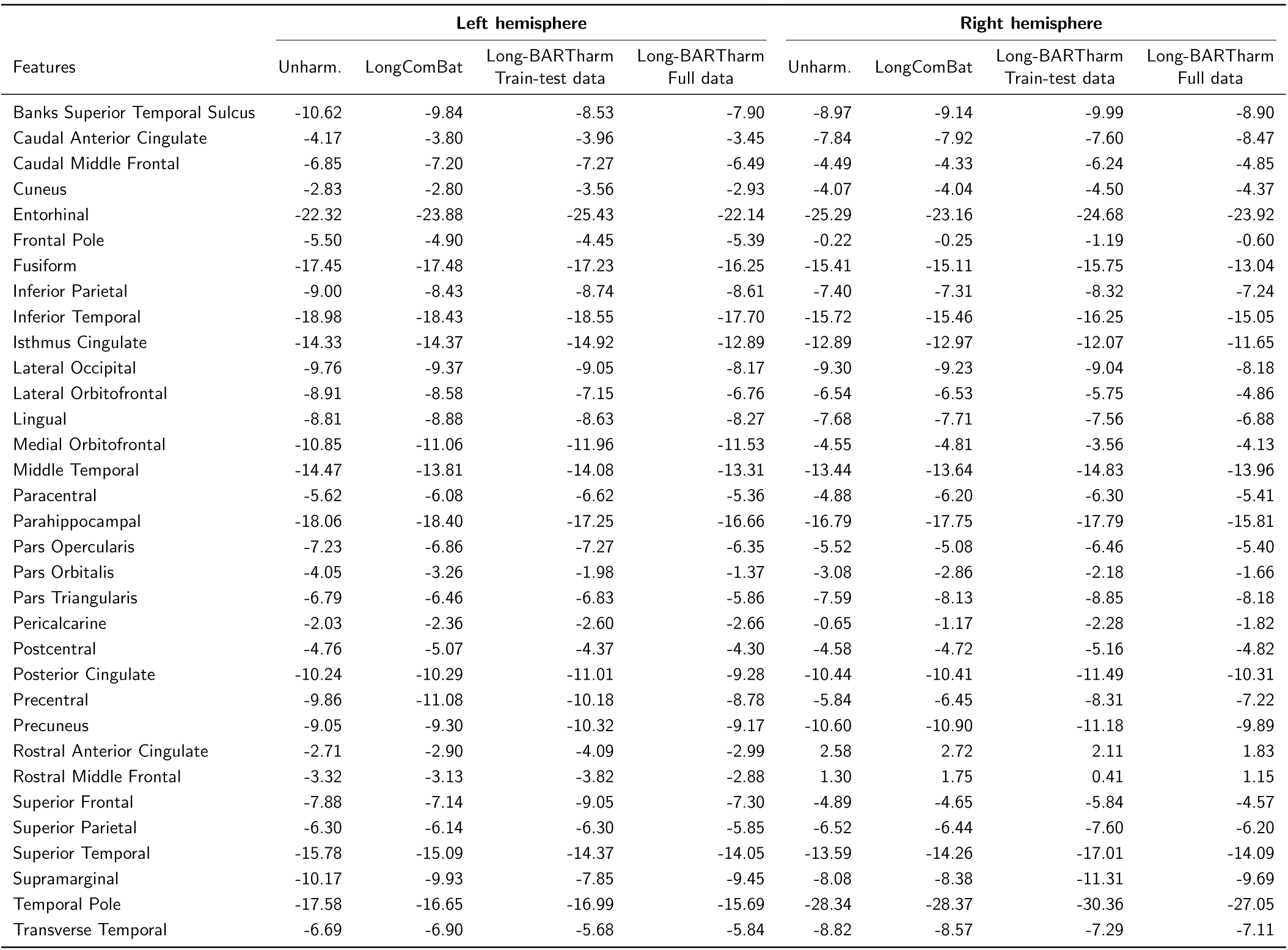
Estimated longitudinal time-effect coefficients for cortical-thickness features across unharmonised data, LongComBat, and Long-BARTharm applied to the train-test and full datasets, separately for the left and right hemispheres. Values are expressed in *µ*m/year.

|  | **Left hemisphere** |  |  |  | **Right hemisphere** |  |  |  |
|---|---|---|---|---|---|---|---|---|
| Features | Unharm. | LongComBat | Long-BARTharm<br>Train-test data | Long-BARTharm<br>Full data | Unharm. | LongComBat | Long-BARTharm<br>Train-test data | Long-BARTharm<br>Full data |
| Banks Superior Temporal Sulcus | -10.62 | -9.84 | -8.53 | -7.90 | -8.97 | -9.14 | -9.99 | -8.90 |
| Caudal Anterior Cingulate | -4.17 | -3.80 | -3.96 | -3.45 | -7.84 | -7.92 | -7.60 | -8.47 |
| Caudal Middle Frontal | -6.85 | -7.20 | -7.27 | -6.49 | -4.49 | -4.33 | -6.24 | -4.85 |
| Cuneus | -2.83 | -2.80 | -3.56 | -2.93 | -4.07 | -4.04 | -4.50 | -4.37 |
| Entorhinal | -22.32 | -23.88 | -25.43 | -22.14 | -25.29 | -23.16 | -24.68 | -23.92 |
| Frontal Pole | -5.50 | -4.90 | -4.45 | -5.39 | -0.22 | -0.25 | -1.19 | -0.60 |
| Fusiform | -17.45 | -17.48 | -17.23 | -16.25 | -15.41 | -15.11 | -15.75 | -13.04 |
| Inferior Parietal | -9.00 | -8.43 | -8.74 | -8.61 | -7.40 | -7.31 | -8.32 | -7.24 |
| Inferior Temporal | -18.98 | -18.43 | -18.55 | -17.70 | -15.72 | -15.46 | -16.25 | -15.05 |
| Isthmus Cingulate | -14.33 | -14.37 | -14.92 | -12.89 | -12.89 | -12.97 | -12.07 | -11.65 |
| Lateral Occipital | -9.76 | -9.37 | -9.05 | -8.17 | -9.30 | -9.23 | -9.04 | -8.18 |
| Lateral Orbitofrontal | -8.91 | -8.58 | -7.15 | -6.76 | -6.54 | -6.53 | -5.75 | -4.86 |
| Lingual | -8.81 | -8.88 | -8.63 | -8.27 | -7.68 | -7.71 | -7.56 | -6.88 |
| Medial Orbitofrontal | -10.85 | -11.06 | -11.96 | -11.53 | -4.55 | -4.81 | -3.56 | -4.13 |
| Middle Temporal | -14.47 | -13.81 | -14.08 | -13.31 | -13.44 | -13.64 | -14.83 | -13.96 |
| Paracentral | -5.62 | -6.08 | -6.62 | -5.36 | -4.88 | -6.20 | -6.30 | -5.41 |
| Parahippocampal | -18.06 | -18.40 | -17.25 | -16.66 | -16.79 | -17.75 | -17.79 | -15.81 |
| Pars Opercularis | -7.23 | -6.86 | -7.27 | -6.35 | -5.52 | -5.08 | -6.46 | -5.40 |
| Pars Orbitalis | -4.05 | -3.26 | -1.98 | -1.37 | -3.08 | -2.86 | -2.18 | -1.66 |
| Pars Triangularis | -6.79 | -6.46 | -6.83 | -5.86 | -7.59 | -8.13 | -8.85 | -8.18 |
| Pericalcarine | -2.03 | -2.36 | -2.60 | -2.66 | -0.65 | -1.17 | -2.28 | -1.82 |
| Postcentral | -4.76 | -5.07 | -4.37 | -4.30 | -4.58 | -4.72 | -5.16 | -4.82 |
| Posterior Cingulate | -10.24 | -10.29 | -11.01 | -9.28 | -10.44 | -10.41 | -11.49 | -10.31 |
| Precentral | -9.86 | -11.08 | -10.18 | -8.78 | -5.84 | -6.45 | -8.31 | -7.22 |
| Precuneus | -9.05 | -9.30 | -10.32 | -9.17 | -10.60 | -10.90 | -11.18 | -9.89 |
| Rostral Anterior Cingulate | -2.71 | -2.90 | -4.09 | -2.99 | 2.58 | 2.72 | 2.11 | 1.83 |
| Rostral Middle Frontal | -3.32 | -3.13 | -3.82 | -2.88 | 1.30 | 1.75 | 0.41 | 1.15 |
| Superior Frontal | -7.88 | -7.14 | -9.05 | -7.30 | -4.89 | -4.65 | -5.84 | -4.57 |
| Superior Parietal | -6.30 | -6.14 | -6.30 | -5.85 | -6.52 | -6.44 | -7.60 | -6.20 |
| Superior Temporal | -15.78 | -15.09 | -14.37 | -14.05 | -13.59 | -14.26 | -17.01 | -14.09 |
| Supramarginal | -10.17 | -9.93 | -7.85 | -9.45 | -8.08 | -8.38 | -11.31 | -9.69 |
| Temporal Pole | -17.58 | -16.65 | -16.99 | -15.69 | -28.34 | -28.37 | -30.36 | -27.05 |
| Transverse Temporal | -6.69 | -6.90 | -5.68 | -5.84 | -8.82 | -8.57 | -7.29 | -7.11 |

Scanner classification using the unharmonised IDPs yielded an overall accuracy of 9.2% and a balanced accuracy of 5.3%, indicating that scanner identity could be predicted from the original imaging features. Following harmonisation, overall accuracy decreased to 2.8% for LongComBat and 6.5% for Long-BARTharm, with corresponding balanced accuracies of 1.3% and 3.4%, respectively. Permutation of scanner labels yielded mean chance-level accuracies of approximately 5.5% overall and 3.0% balanced accuracy, with 95% permutation intervals for balanced accuracy ranging from approximately 1.8% to 4.5% across methods. Classification performance was significantly above chance for the unharmonised IDPs (permutation *p* = 0.002 for both metrics), whereas neither harmonisation method showed significantly above-chance performance (LongComBat: *p* = 0.999 for overall accuracy and *p* = 1.000 for balanced accuracy; Long-BARTharm: *p* = 0.195 and *p* = 0.282, respectively). These findings indicate that scanner identity was more readily predicted from the unharmonised IDPs, while both harmonisation methods reduced the scanner-related information available to the classifier.

Scanner-specific residual variances for cortical thickness in the left inferior temporal ROI are shown in Figure 1. The unharmonised data exhibited significant heterogeneity in residual variance across scanners (FK *χ*^2^ = 48.45*, p* = 0.031). Following harmonisation, this heterogeneity was reduced for both Long-BARTharm (FK *χ*^2^ = 11.03*, p* = 0.999) and LongComBat (FK *χ*^2^ = 18.56*, p* = 0.971). Across the 66 cortical-thickness IDPs, Long-BARTharm yielded smaller FK statistics than LongComBat in 59 regions, indicating lower residual variance heterogeneity in the majority of cortical regions. Both harmonisation methods yielded smaller FK statistics than the unharmonised data across all 66 regions. Complete results are provided in Supplementary Table 8.

**Figure 1:**
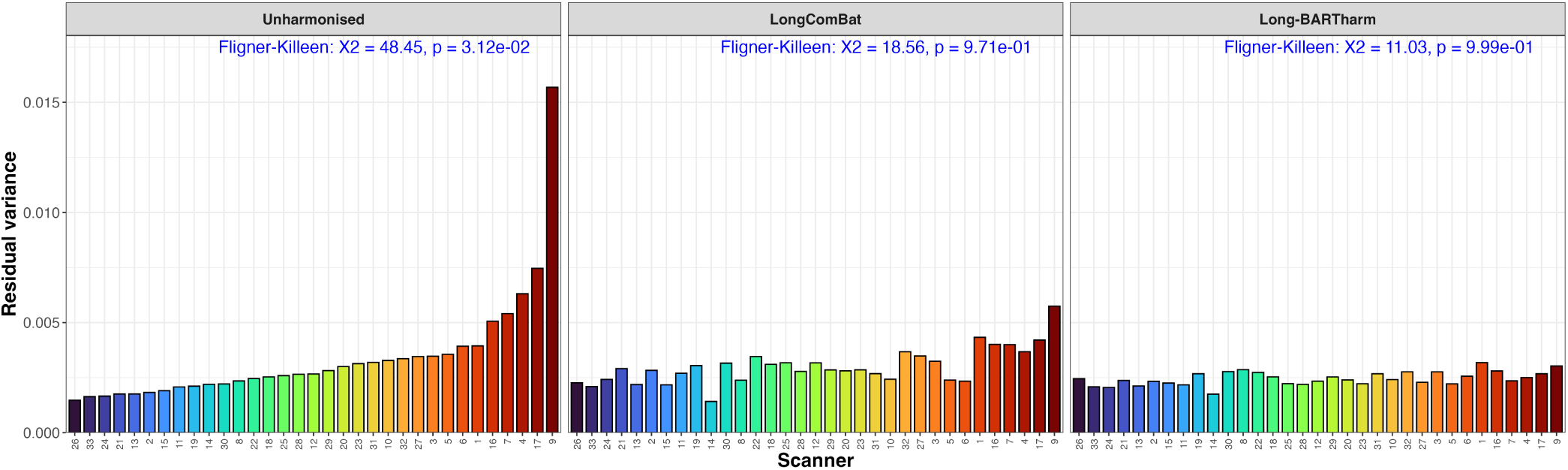
Scanner-specific residual variances for the left inferior temporal ROI in unharmonised data, Long-BARTharm, and LongComBat. Bars show within-scanner residual variance from linear mixed-effects. The Fligner–Killeen *χ*^2^ statistic and *p*-value test homogeneity of residual variances across scanners. Bars are sorted by increasing residual variance in the unharmonised data. ROI: region of interest.

Long-BARTharm and LongComBat identified comparable numbers of significant longitudinal time effects across the 66 cortical-thickness IDPs, with both methods showing a modest increase relative to the unharmonised data at nominal and Bonferroni-adjusted significance thresholds (Supplementary Table 5). Among IDPs with significant effects, Long-BARTharm yielded smaller nominal *p*-values in 29 IDPs, compared with 18 for LongComBat, and smaller Bonferroni-adjusted *p*-values in 11 IDPs, compared with 9 for LongComBat (Figure 2A). Regional *p*-value patterns are illustrated in Figure 2B, with complete results provided in Supplementary Table 7. Estimated longitudinal time coefficients were predominantly negative, consistent with cortical thinning over time in cognitively normal participants, and showed broadly comparable magnitudes across methods (Supplementary Figure 7; Supplementary Table 6).

**Figure 2:**
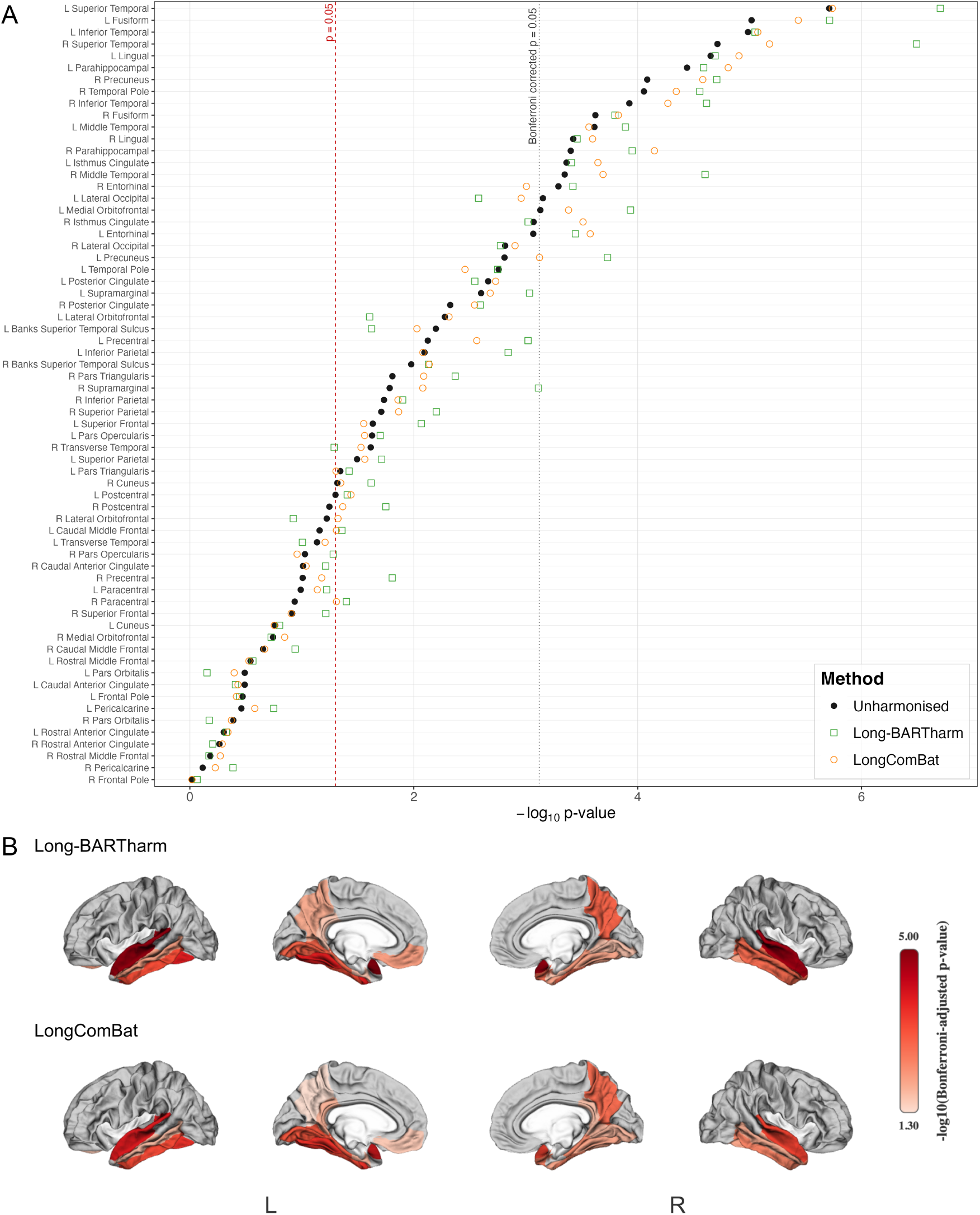
A. Nominal Kenward–Roger *p*-values for longitudinal time effects in cognitively normal participants, obtained from linear mixed-effects models fitted to cortical-thickness IDPs in unharmonised data and following Long-BARTharm and LongComBat harmonisation. Dashed and dotted lines indicate the nominal *p* = 0.05 and Bonferroni-corrected significance thresholds, respectively. B. Bonferroni-adjusted *p*-values after harmonisation for Long-BARTharm and LongComBat, mapped onto the cortical surface. Grey indicates non-significant regions; light pink and dark red indicate higher and lower adjusted *p*-values, respectively.

#### 3.2.2 Posterior predictive

In the cross-sectional analysis comparing train-test and full-data harmonisation, both BARTharm approaches and ComBat identified comparable numbers of significant cross-sectional age associations across the 66 cortical-thickness IDPs at the nominal significance threshold (Supplementary Table 5). In pairwise comparisons restricted to IDPs with significant effects, BARTharm yielded smaller nominal *p*-values than ComBat in 37 of 57 IDPs under combined training and test-set harmonisation and 33 of 57 IDPs under full-data harmonisation. This pattern persisted after Bonferroni correction, with BARTharm yielding smaller adjusted *p*-values in 29 of 41 and 25 of 41 significant IDPs, respectively. Regional *p*-value patterns are illustrated in Figure 3A, with complete results provided in Supplementary Table 9. Age-effect coefficients were broadly comparable across harmonisation approaches, with estimates visualised in Supplementary Figure 8A and reported in Supplementary Table 9, respectively.

**Figure 3:**
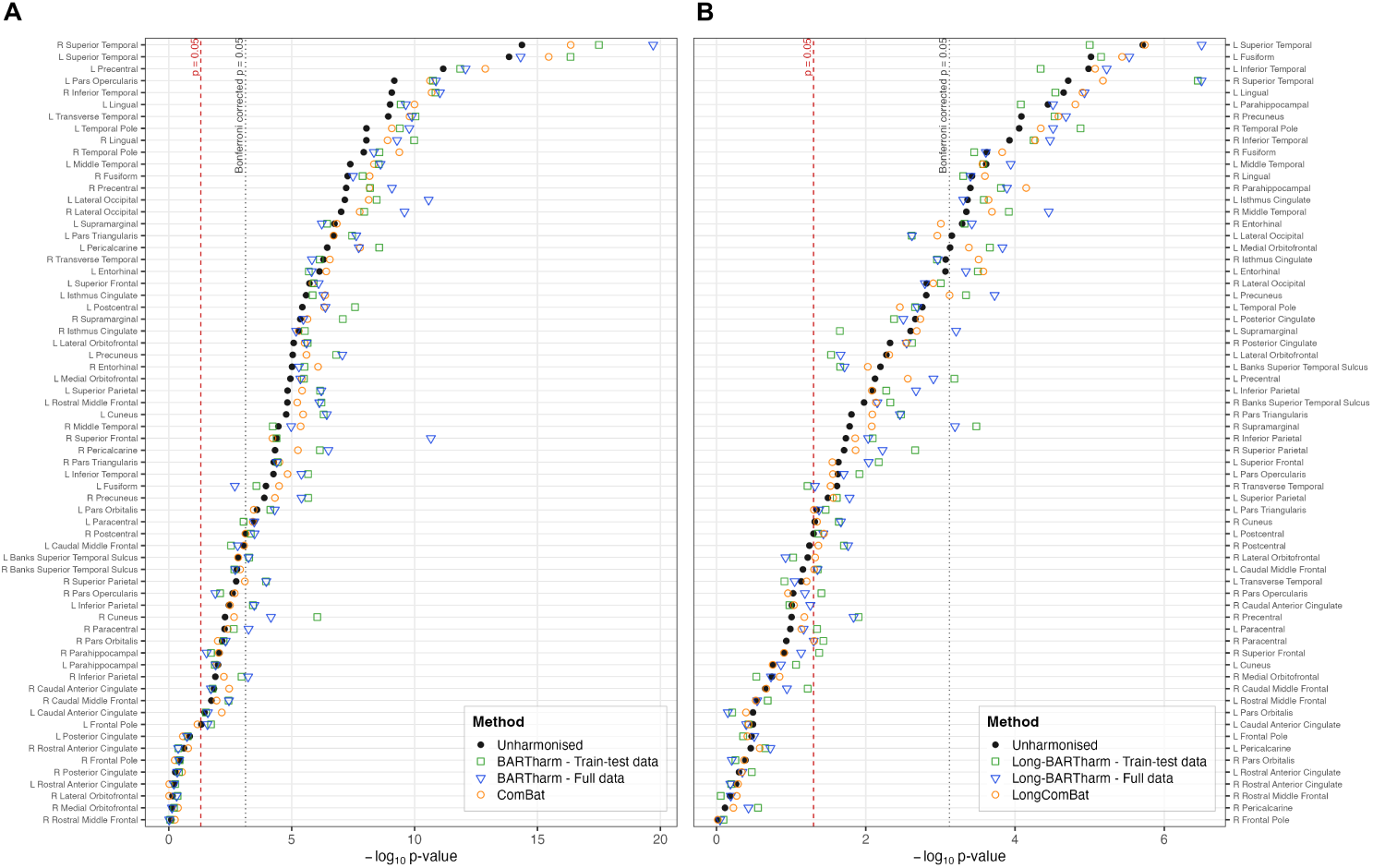
Nominal *p*-values for cortical-thickness IDPs in the combined training/test and full-data harmonisation analyses. A. Cross-sectional age-effect *p*-values obtained from linear regression models adjusted for sex, diagnosis, and MMSE score, comparing unharmonised data, ComBat, and BARTharm. B. Longitudinal time-effect *p*-values obtained from linear mixed-effects models using Kenward–Roger tests, comparing unharmonised data, LongComBat, and Long-BARTharm. Dashed and dotted lines indicate the nominal *p* = 0.05 and Bonferroni-corrected significance thresholds, respectively.

When comparing the test-set harmonisation and full-data harmonisation, full-data harmonisation with BARTharm and ComBat identified similar numbers of significant associations, whereas test-set harmonisation identified fewer, comparable to the unharmonised data (Supplementary Table 5). However, BARTharm yielded smaller significant nominal *p*-values in 6 of 9 IDPs following test-set harmonisation and 6 of 12 following full-data harmonisation than ComBat (Figure 4). Estimated age-effect coefficients were broadly comparable across harmonisation approaches. Regional estimates and *p*-values are provided in Supplementary Tables 13 and 14, respectively.

**Figure 4:**
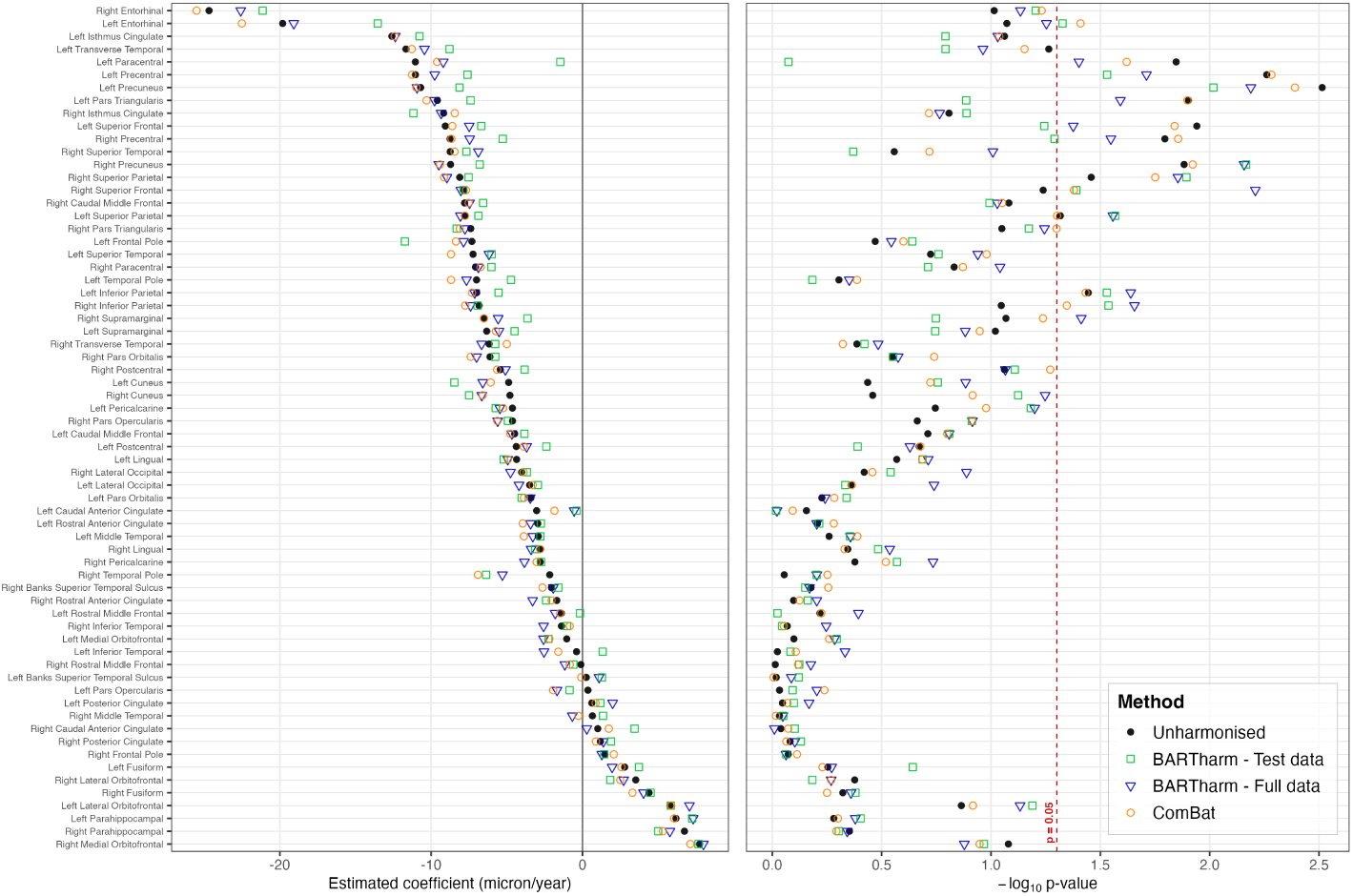
Cross-sectional age-effect coefficients and corresponding nominal *p*-values for cortical-thickness IDPs in the held-out test set (*n* = 35). Estimates were obtained from linear regression models. Methods include unharmonised data, cross-sectional ComBat, and cross-sectional BARTharm harmonisation using posterior predictive and full-data approaches, with all analyses restricted to the same test participants. Dashed line indicates the nominal *p* = 0.05.

In the longitudinal analysis comparing combined training and test-set and full-data harmonisation, all harmonisation approaches identified similar numbers of significant time effects, exceeding the unharmonised data at both nominal and Bonferroni-adjusted thresholds, with Long-BARTharm retaining slightly more associations than LongComBat after correction (Supplementary Table 5). In pairwise comparisons restricted to IDPs with significant effects, Long-BARTharm yielded smaller nominal *p*-values in 26 of 44 IDPs with combined training/test harmonisation and 30 of 45 with full-data harmonisation. After Bonferroni correction, Long-BARTharm yielded smaller adjusted *p*-values in 7 of 18 and 12 of 18 significant IDPs, respectively (Figure 3B). Estimated longitudinal time-effect coefficients were also predominantly negative across cortical-thickness IDPs, consistent with cortical thinning over time, and showed broadly comparable magnitudes across harmonisation approaches (Supplementary Figure 8B). Regional estimates and *p*-values are provided in Supplementary Tables 11 and 12, respectively.

**Table 12:** Kenward–Roger *p*-values for longitudinal time-effect coefficients of cortical-thickness features across unharmonised data, LongComBat, and Long-BARTharm applied to the train-test and full datasets, separately for the left and right hemispheres.

|  | **Left hemisphere** |  |  |  | **Right hemisphere** |  |  |  |
|---|---|---|---|---|---|---|---|---|
| Features | Unharm. | LongComBat | Long-BARTharm<br>Train-test data | Long-BARTharm<br>Full data | Unharm. | LongComBat | Long-BARTharm<br>Train-test data | Long-BARTharm<br>Full data |
| Banks Superior Temporal Sulcus | 0.006 | 0.009 | 0.022 | 0.019 | 0.011 | 0.007 | 0.005 | 0.007 |
| Caudal Anterior Cingulate | 0.324 | 0.372 | 0.373 | 0.399 | 0.098 | 0.092 | 0.105 | 0.055 |
| Caudal Middle Frontal | 0.069 | 0.049 | 0.044 | 0.044 | 0.222 | 0.215 | 0.060 | 0.114 |
| Cuneus | 0.173 | 0.176 | 0.086 | 0.137 | 0.048 | 0.045 | 0.023 | 0.022 |
| Entorhinal | 8.57 × 10<sup>−4</sup> | 2.65 × 10<sup>−4</sup> | 3.14 × 10<sup>−4</sup> | 4.56 × 10<sup>−4</sup> | 5.10 × 10<sup>−4</sup> | 9.85 × 10<sup>−4</sup> | 4.75 × 10<sup>−4</sup> | 3.79 × 10<sup>−4</sup> |
| Frontal Pole | 0.338 | 0.385 | 0.439 | 0.312 | 0.964 | 0.959 | 0.803 | 0.893 |
| Fusiform | 9.61 × 10<sup>−6</sup> | 3.68 × 10<sup>−6</sup> | 7.02 × 10<sup>−6</sup> | 2.95 × 10<sup>−6</sup> | 2.39 × 10<sup>−4</sup> | 1.49 × 10<sup>−4</sup> | 3.52 × 10<sup>−4</sup> | 2.46 × 10<sup>−4</sup> |
| Inferior Parietal | 0.008 | 0.008 | 0.005 | 0.002 | 0.018 | 0.014 | 0.008 | 0.009 |
| Inferior Temporal | 1.03 × 10<sup>−5</sup> | 8.45 × 10<sup>−6</sup> | 4.54 × 10<sup>−5</sup> | 5.91 × 10<sup>−6</sup> | 1.19 × 10<sup>−4</sup> | 5.37 × 10<sup>−5</sup> | 5.62 × 10<sup>−5</sup> | 3.39 × 10<sup>−5</sup> |
| Isthmus Cingulate | 4.32 × 10<sup>−4</sup> | 2.26 × 10<sup>−4</sup> | 2.62 × 10<sup>−4</sup> | 4.94 × 10<sup>−4</sup> | 8.50 × 10<sup>−4</sup> | 3.08 × 10<sup>−4</sup> | 0.001 | 0.001 |
| Lateral Occipital | 7.02 × 10<sup>−4</sup> | 0.001 | 0.002 | 0.002 | 0.002 | 0.001 | 9.87 × 10<sup>−4</sup> | 0.002 |
| Lateral Orbitofrontal | 0.005 | 0.005 | 0.029 | 0.022 | 0.060 | 0.048 | 0.094 | 0.119 |
| Lingual | 2.23 × 10<sup>−5</sup> | 1.24 × 10<sup>−5</sup> | 2.88 × 10<sup>−5</sup> | 1.17 × 10<sup>−5</sup> | 3.77 × 10<sup>−4</sup> | 2.53 × 10<sup>−4</sup> | 4.94 × 10<sup>−4</sup> | 3.95 × 10<sup>−4</sup> |
| Medial Orbitofrontal | 7.40 × 10<sup>−4</sup> | 4.15 × 10<sup>−4</sup> | 2.17 × 10<sup>−4</sup> | 1.47 × 10<sup>−4</sup> | 0.181 | 0.143 | 0.292 | 0.186 |
| Middle Temporal | 2.43 × 10<sup>−4</sup> | 2.73 × 10<sup>−4</sup> | 2.65 × 10<sup>−4</sup> | 1.14 × 10<sup>−4</sup> | 4.49 × 10<sup>−4</sup> | 2.04 × 10<sup>−4</sup> | 1.21 × 10<sup>−4</sup> | 3.53 × 10<sup>−5</sup> |
| Paracentral | 0.102 | 0.073 | 0.045 | 0.068 | 0.116 | 0.049 | 0.037 | 0.050 |
| Parahippocampal | 3.62 × 10<sup>−5</sup> | 1.55 × 10<sup>−5</sup> | 8.34 × 10<sup>−5</sup> | 3.10 × 10<sup>−5</sup> | 3.96 × 10<sup>−4</sup> | 7.08 × 10<sup>−5</sup> | 1.54 × 10<sup>−4</sup> | 1.28 × 10<sup>−4</sup> |
| Pars Opercularis | 0.024 | 0.028 | 0.012 | 0.020 | 0.094 | 0.110 | 0.039 | 0.065 |
| Pars Orbitalis | 0.324 | 0.402 | 0.616 | 0.704 | 0.409 | 0.426 | 0.556 | 0.623 |
| Pars Triangularis | 0.045 | 0.050 | 0.034 | 0.042 | 0.016 | 0.008 | 0.003 | 0.003 |
| Pericalcarine | 0.347 | 0.263 | 0.222 | 0.190 | 0.769 | 0.593 | 0.278 | 0.372 |
| Postcentral | 0.050 | 0.036 | 0.043 | 0.037 | 0.057 | 0.043 | 0.020 | 0.017 |
| Posterior Cingulate | 0.002 | 0.002 | 0.004 | 0.003 | 0.005 | 0.003 | 0.002 | 0.003 |
| Precentral | 0.007 | 0.003 | 6.49 × 10<sup>−4</sup> | 0.001 | 0.098 | 0.066 | 0.012 | 0.015 |
| Precuneus | 0.002 | 7.53 × 10<sup>−4</sup> | 4.53 × 10<sup>−4</sup> | 1.88 × 10<sup>−4</sup> | 8.21 × 10<sup>−5</sup> | 2.62 × 10<sup>−5</sup> | 2.95 × 10<sup>−5</sup> | 2.08 × 10<sup>−5</sup> |
| Rostral Anterior Cingulate | 0.498 | 0.457 | 0.336 | 0.450 | 0.545 | 0.514 | 0.636 | 0.650 |
| Rostral Middle Frontal | 0.287 | 0.297 | 0.206 | 0.285 | 0.658 | 0.535 | 0.877 | 0.642 |
| Superior Frontal | 0.023 | 0.028 | 0.007 | 0.009 | 0.122 | 0.124 | 0.042 | 0.073 |
| Superior Parietal | 0.032 | 0.027 | 0.024 | 0.016 | 0.020 | 0.014 | 0.002 | 0.006 |
| Superior Temporal | 1.94 × 10<sup>−6</sup> | 1.82 × 10<sup>−6</sup> | 9.97 × 10<sup>−6</sup> | 3.16 × 10<sup>−7</sup> | 1.94 × 10<sup>−5</sup> | 6.64 × 10<sup>−6</sup> | 3.55 × 10<sup>−7</sup> | 3.16 × 10<sup>−7</sup> |
| Supramarginal | 0.003 | 0.002 | 0.022 | 6.15 × 10<sup>−4</sup> | 0.016 | 0.008 | 3.30 × 10<sup>−4</sup> | 6.37 × 10<sup>−4</sup> |
| Temporal Pole | 0.002 | 0.003 | 0.002 | 0.002 | 8.79 × 10<sup>−5</sup> | 4.52 × 10<sup>−5</sup> | 1.32 × 10<sup>−5</sup> | 3.09 × 10<sup>−5</sup> |
| Transverse Temporal | 0.073 | 0.062 | 0.123 | 0.090 | 0.024 | 0.030 | 0.060 | 0.048 |

**Table 13:** Estimated cross-sectional age-effect coefficients for cortical-thickness features across unharmonised data, ComBat, and BARTharm applied to the test and full datasets, separately for the left and right hemispheres. Values are expressed in *µ*m/year.

|  | Left hemisphere |  |  |  | Right hemisphere |  |  |  |
|---|---|---|---|---|---|---|---|---|
| Features | Unharm. | ComBat | BARTharm<br>Test data | BARTharm<br>Full data | Unharm. | ComBat | BARTharm<br>Test data | BARTharm<br>Full data |
| Banks Superior Temporal Sulcus | 0.25 | -0.08 | 1.29 | 1.10 | -2.05 | -2.64 | -1.59 | -1.93 |
| Caudal Anterior Cingulate | -3.03 | -1.85 | -0.39 | -0.55 | 1.01 | 1.74 | 3.45 | 0.29 |
| Caudal Middle Frontal | -4.48 | -4.77 | -3.84 | -4.67 | -7.79 | -7.52 | -6.57 | -7.46 |
| Cuneus | -4.88 | -6.07 | -8.47 | -6.59 | -4.80 | -6.60 | -7.50 | -6.65 |
| Entorhinal | -19.82 | -22.51 | -13.53 | -19.08 | -24.68 | -25.51 | -21.14 | -22.59 |
| Frontal Pole | -7.31 | -8.37 | -11.74 | -7.86 | 1.48 | 2.07 | 1.43 | 1.28 |
| Fusiform | 2.79 | 2.57 | 3.73 | 1.96 | 4.40 | 3.31 | 4.50 | 4.04 |
| Inferior Parietal | -6.99 | -7.30 | -5.55 | -7.13 | -6.84 | -7.75 | -6.93 | -7.39 |
| Inferior Temporal | -0.39 | -1.60 | 1.34 | -2.54 | -1.39 | -0.84 | -1.02 | -2.57 |
| Isthmus Cingulate | -12.60 | -12.41 | -10.77 | -12.37 | -9.18 | -8.44 | -11.17 | -9.34 |
| Lateral Occipital | -3.49 | -3.29 | -2.95 | -4.19 | -4.02 | -3.89 | -3.68 | -4.76 |
| Lateral Orbitofrontal | 5.79 | 5.86 | 5.83 | 7.07 | 3.52 | 2.49 | 1.84 | 2.72 |
| Lingual | -4.35 | -4.93 | -5.20 | -4.94 | -2.80 | -2.80 | -3.12 | -3.39 |
| Medial Orbitofrontal | -1.04 | -2.19 | -2.26 | -2.57 | 7.74 | 7.13 | 7.67 | 8.00 |
| Middle Temporal | -2.92 | -3.87 | -2.78 | -3.28 | 0.66 | -0.26 | 1.36 | -0.67 |
| Paracentral | -11.04 | -9.61 | -1.47 | -9.20 | -7.07 | -6.74 | -6.02 | -6.88 |
| Parahippocampal | 6.18 | 6.06 | 7.25 | 7.32 | 6.74 | 5.32 | 5.00 | 5.78 |
| Pars Opercularis | 0.36 | -1.94 | -0.85 | -1.68 | -4.62 | -5.63 | -4.93 | -5.61 |
| Pars Orbitalis | -3.41 | -3.87 | -4.01 | -3.45 | -6.12 | -7.39 | -5.76 | -7.00 |
| Pars Triangularis | -9.59 | -10.30 | -7.39 | -9.78 | -7.39 | -8.11 | -8.32 | -7.76 |
| Pericalcarine | -4.63 | -5.24 | -5.73 | -5.46 | -2.79 | -3.03 | -2.73 | -3.84 |
| Postcentral | -4.37 | -3.88 | -2.39 | -3.69 | -5.43 | -5.64 | -3.82 | -5.09 |
| Posterior Cingulate | 0.62 | 0.87 | 1.18 | 2.00 | 1.19 | 0.89 | 1.89 | 1.38 |
| Precentral | -11.04 | -11.26 | -7.60 | -9.77 | -8.75 | -8.66 | -5.27 | -7.45 |
| Precuneus | -10.69 | -10.98 | -8.12 | -10.94 | -8.72 | -9.42 | -6.80 | -9.49 |
| Rostral Anterior Cingulate | -2.96 | -3.95 | -2.75 | -3.43 | -1.70 | -2.04 | -2.41 | -3.30 |
| Rostral Middle Frontal | -1.40 | -1.48 | -0.17 | -1.80 | -0.11 | -0.84 | -0.60 | -1.16 |
| Superior Frontal | -9.07 | -8.60 | -6.69 | -7.48 | -7.83 | -7.70 | -7.87 | -8.04 |
| Superior Parietal | -7.75 | -7.77 | -6.88 | -8.07 | -8.11 | -9.15 | -7.52 | -8.96 |
| Superior Temporal | -7.23 | -8.69 | -6.00 | -6.18 | -8.75 | -8.45 | -7.67 | -6.88 |
| Supramarginal | -6.33 | -5.72 | -4.49 | -5.51 | -6.49 | -6.51 | -3.63 | -5.57 |
| Temporal Pole | -7.00 | -8.69 | -4.73 | -7.67 | -2.16 | -6.91 | -6.37 | -5.29 |
| Transverse Temporal | -11.67 | -11.28 | -8.79 | -10.44 | -6.17 | -5.00 | -5.77 | -6.67 |

**Table 14:** Nominal *p*-values for cross-sectional age-effect coefficients of cortical-thickness features across unharmonised data, ComBat, and BARTharm applied to the test and full datasets, separately for the left and right hemispheres.

|  | Left hemisphere |  |  |  | Right hemisphere |  |  |  |
|---|---|---|---|---|---|---|---|---|
| Features | Unharm. | ComBat | BARTharm<br>Test data | BARTharm<br>Full data | Unharm. | ComBat | BARTharm<br>Test data | BARTharm<br>Full data |
| Banks Superior Temporal Sulcus | 0.959 | 0.987 | 0.756 | 0.818 | 0.665 | 0.555 | 0.707 | 0.675 |
| Caudal Anterior Cingulate | 0.698 | 0.807 | 0.961 | 0.952 | 0.913 | 0.846 | 0.789 | 0.978 |
| Caudal Middle Frontal | 0.194 | 0.159 | 0.155 | 0.155 | 0.083 | 0.089 | 0.102 | 0.094 |
| Cuneus | 0.366 | 0.189 | 0.175 | 0.131 | 0.347 | 0.121 | 0.075 | 0.057 |
| Entorhinal | 0.085 | 0.039 | 0.047 | 0.056 | 0.097 | 0.059 | 0.062 | 0.073 |
| Frontal Pole | 0.338 | 0.251 | 0.229 | 0.286 | 0.846 | 0.771 | 0.865 | 0.865 |
| Fusiform | 0.557 | 0.589 | 0.228 | 0.535 | 0.476 | 0.562 | 0.417 | 0.436 |
| Inferior Parietal | 0.036 | 0.037 | 0.030 | 0.023 | 0.090 | 0.045 | 0.029 | 0.022 |
| Inferior Temporal | 0.947 | 0.780 | 0.826 | 0.465 | 0.854 | 0.886 | 0.902 | 0.567 |
| Isthmus Cingulate | 0.087 | 0.092 | 0.162 | 0.093 | 0.156 | 0.192 | 0.129 | 0.172 |
| Lateral Occipital | 0.436 | 0.433 | 0.464 | 0.182 | 0.380 | 0.349 | 0.288 | 0.129 |
| Lateral Orbitofrontal | 0.137 | 0.121 | 0.065 | 0.074 | 0.420 | 0.538 | 0.656 | 0.538 |
| Lingual | 0.270 | 0.206 | 0.205 | 0.194 | 0.452 | 0.467 | 0.328 | 0.290 |
| Medial Orbitofrontal | 0.797 | 0.550 | 0.507 | 0.518 | 0.083 | 0.113 | 0.108 | 0.132 |
| Middle Temporal | 0.550 | 0.408 | 0.441 | 0.439 | 0.927 | 0.963 | 0.891 | 0.895 |
| Paracentral | 0.014 | 0.024 | 0.843 | 0.040 | 0.147 | 0.134 | 0.194 | 0.091 |
| Parahippocampal | 0.523 | 0.501 | 0.395 | 0.417 | 0.444 | 0.509 | 0.497 | 0.454 |
| Pars Opercularis | 0.927 | 0.577 | 0.807 | 0.626 | 0.217 | 0.122 | 0.122 | 0.122 |
| Pars Orbitalis | 0.593 | 0.521 | 0.457 | 0.573 | 0.282 | 0.182 | 0.280 | 0.266 |
| Pars Triangularis | 0.013 | 0.013 | 0.130 | 0.026 | 0.089 | 0.050 | 0.067 | 0.057 |
| Pericalcarine | 0.180 | 0.105 | 0.066 | 0.063 | 0.419 | 0.302 | 0.269 | 0.184 |
| Postcentral | 0.213 | 0.210 | 0.407 | 0.235 | 0.087 | 0.054 | 0.078 | 0.086 |
| Posterior Cingulate | 0.898 | 0.852 | 0.797 | 0.679 | 0.829 | 0.863 | 0.740 | 0.788 |
| Precentral | 0.005 | 0.005 | 0.029 | 0.019 | 0.016 | 0.014 | 0.051 | 0.028 |
| Precuneus | 0.003 | 0.004 | 0.010 | 0.006 | 0.013 | 0.012 | 0.007 | 0.007 |
| Rostral Anterior Cingulate | 0.621 | 0.523 | 0.610 | 0.626 | 0.800 | 0.750 | 0.688 | 0.626 |
| Rostral Middle Frontal | 0.605 | 0.597 | 0.947 | 0.404 | 0.969 | 0.761 | 0.753 | 0.666 |
| Superior Frontal | 0.011 | 0.014 | 0.057 | 0.042 | 0.058 | 0.042 | 0.041 | 0.006 |
| Superior Parietal | 0.048 | 0.050 | 0.027 | 0.028 | 0.035 | 0.018 | 0.013 | 0.014 |
| Superior Temporal | 0.189 | 0.105 | 0.174 | 0.115 | 0.277 | 0.191 | 0.426 | 0.098 |
| Supramarginal | 0.095 | 0.113 | 0.180 | 0.131 | 0.085 | 0.058 | 0.179 | 0.039 |
| Temporal Pole | 0.496 | 0.410 | 0.655 | 0.445 | 0.882 | 0.559 | 0.626 | 0.626 |
| Transverse Temporal | 0.054 | 0.070 | 0.161 | 0.109 | 0.410 | 0.476 | 0.379 | 0.328 |

**Table 15:**
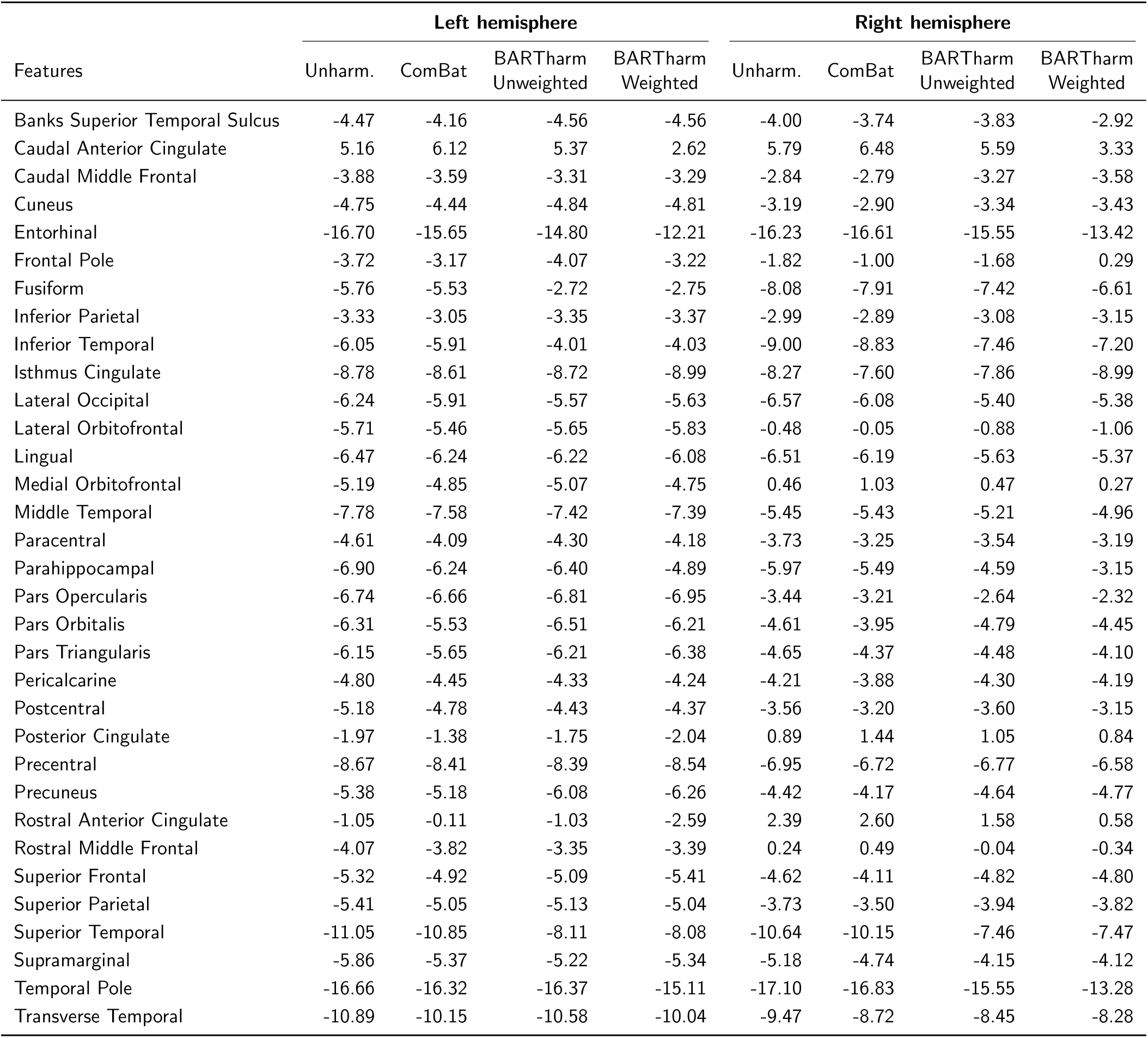
Estimated cross-sectional age-effect coefficients for cortical-thickness features across unharmonised data, ComBat, and BARTharm with and without uncertainty weighting, separately for the left and right hemispheres. Values are expressed in *µ*m/year.

Scatter plots comparing posterior predictive Long-BARTharm harmonisation of the test set with the corresponding full-data harmonised values showed close agreement across cortical volume IDPs (Supplementary Figure 10). Values were concentrated around the identity line, indicating that posterior predictive harmonisation closely reproduced the results obtained when all subject-timepoints were included in the harmonisation procedure. This consistency suggests that Long-BARTharm can effectively harmonise previously unseen subject-timepoints while preserving the harmonised values obtained from the full dataset.

#### 3.2.3 Uncertainty weighting

In the cross-sectional analysis, all approaches identified similar numbers of significant age associations across the 66 cortical-thickness IDPs at the nominal threshold, although both BARTharm approaches retained more associations after Bonferroni correction (Supplementary Table 5). Despite identifying comparable numbers of significant IDPs, weighted BARTharm yielded smaller nominal *p*-values than unweighted BARTharm in 50 of the 55 IDPs significant under both approaches. This pattern was also observed after Bonferroni correction, with smaller adjusted *p*-values in 43 of 46 significant IDPs (Figure 6A). Estimated age-effect coefficients were similar across harmonisation approaches (Supplementary Figure 9A). Regional estimates and *p*-values are provided in Supplementary Tables 15 and 16.

**Figure 5:**
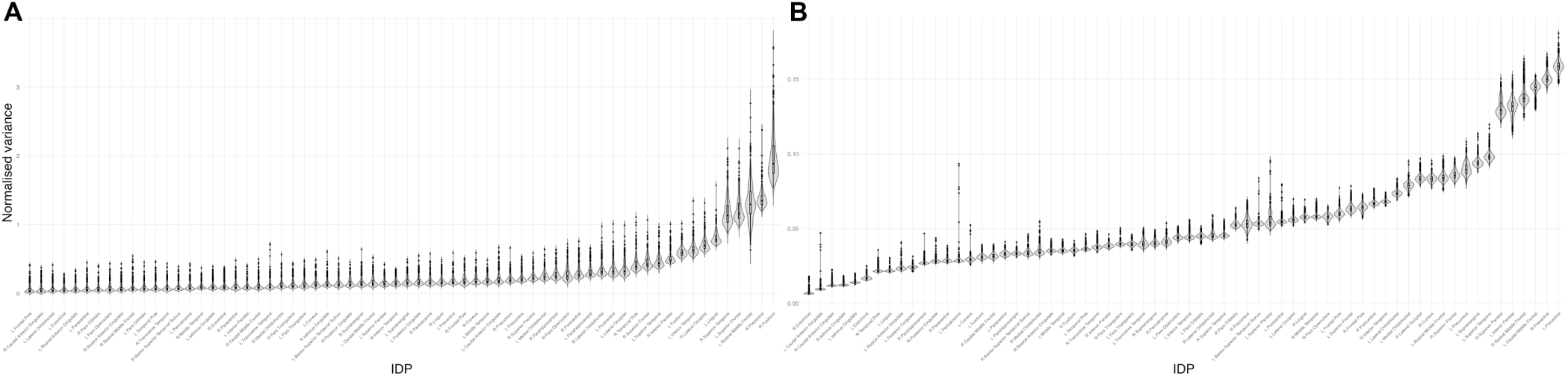
Distribution of normalised posterior uncertainty across cortical-volume regions of interest (ROIs) for A. cross-sectional BARTharm and B. Long-BARTharm. Violin plots show the distribution of normalised posterior variance across ROIs, with wider sections indicating greater density of observations. ROIs are ordered by increasing median normalised posterior variance within each analysis.

**Figure 6:**
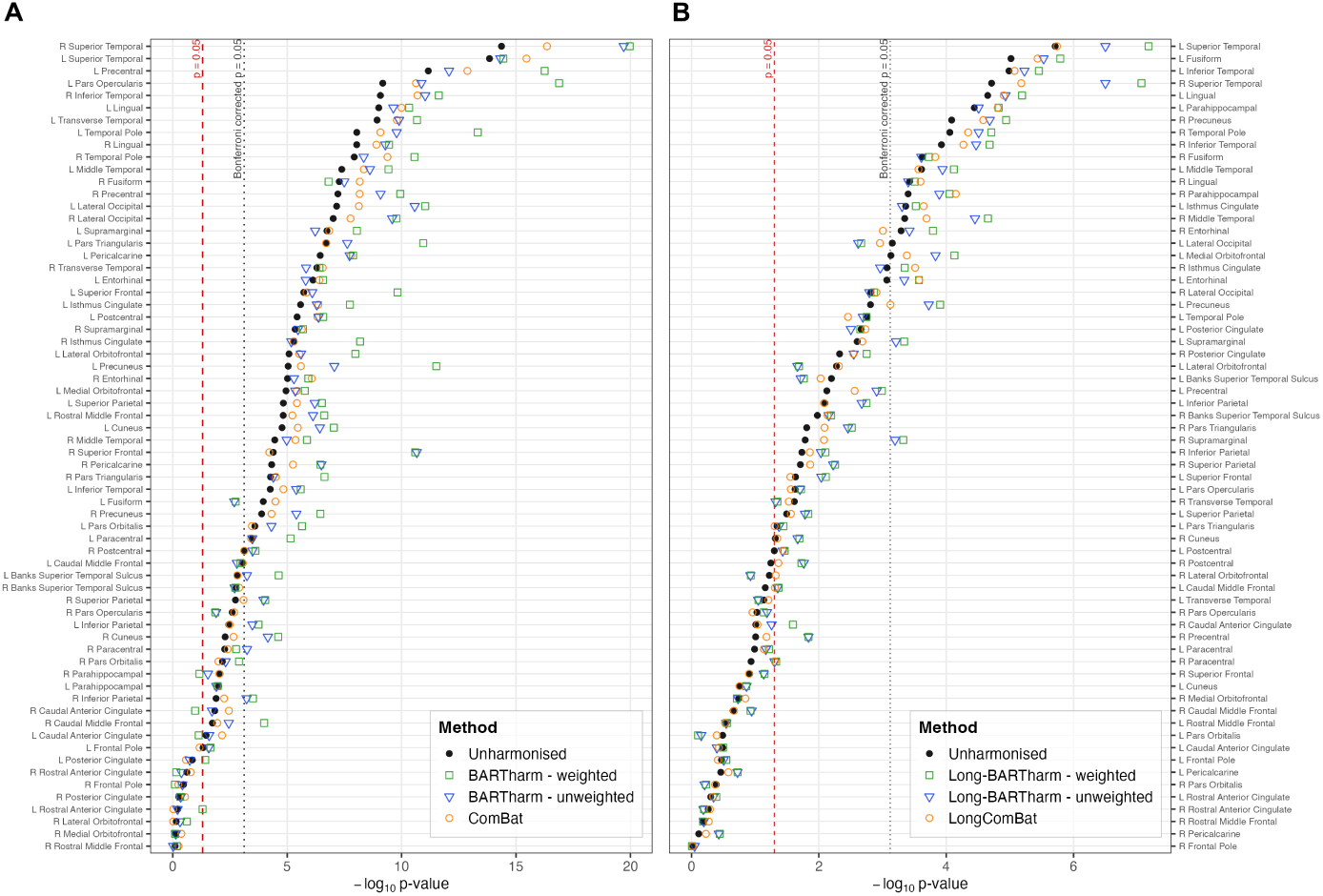
Effect of uncertainty weighting on the statistical significance of age- and time-related associations across cortical-thickness IDPs. A. Cross-sectional age-effect *p*-values from linear regression models. B. Longitudinal time-effect *p*-values from Kenward–Roger tests of linear mixed-effects models. Both panels compare unharmonised data, ComBat or LongComBat, and BARTharm or Long-BARTharm, with and without inverse-variance weighting. Dashed and dotted lines indicate the nominal *p* = 0.05 and Bonferroni-adjusted significance thresholds, respectively.

**Figure 7:**
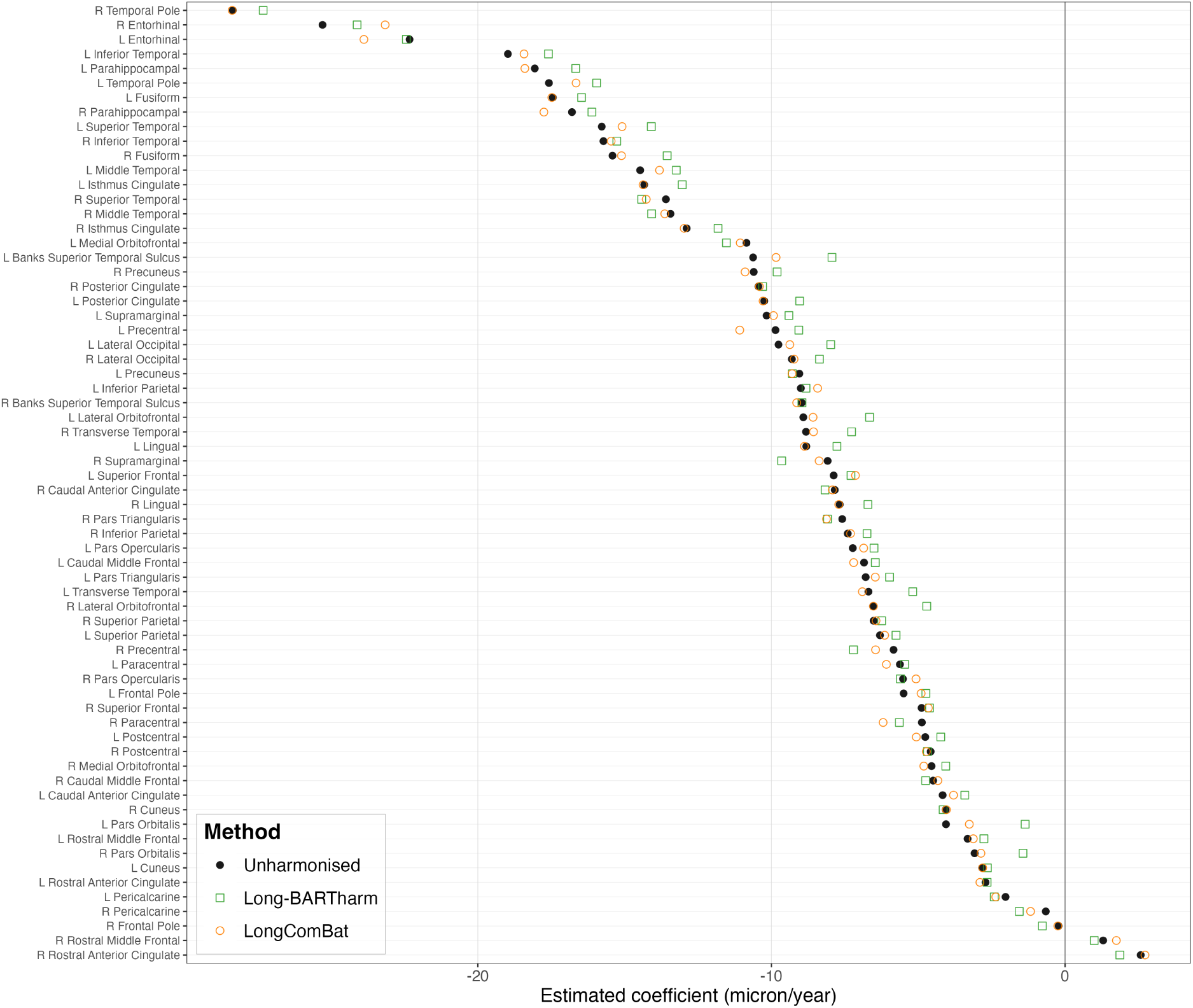
Estimated longitudinal time coefficients for cortical-thickness IDPs in unharmonised data and following Long-BARTharm and LongComBat harmonisation. Coefficients represent the estimated annual change in cortical thickness (µm/year) in cognitively normal participants, obtained from linear mixed-effects models. The horizontal reference line indicates zero longitudinal change.

**Figure 8:**
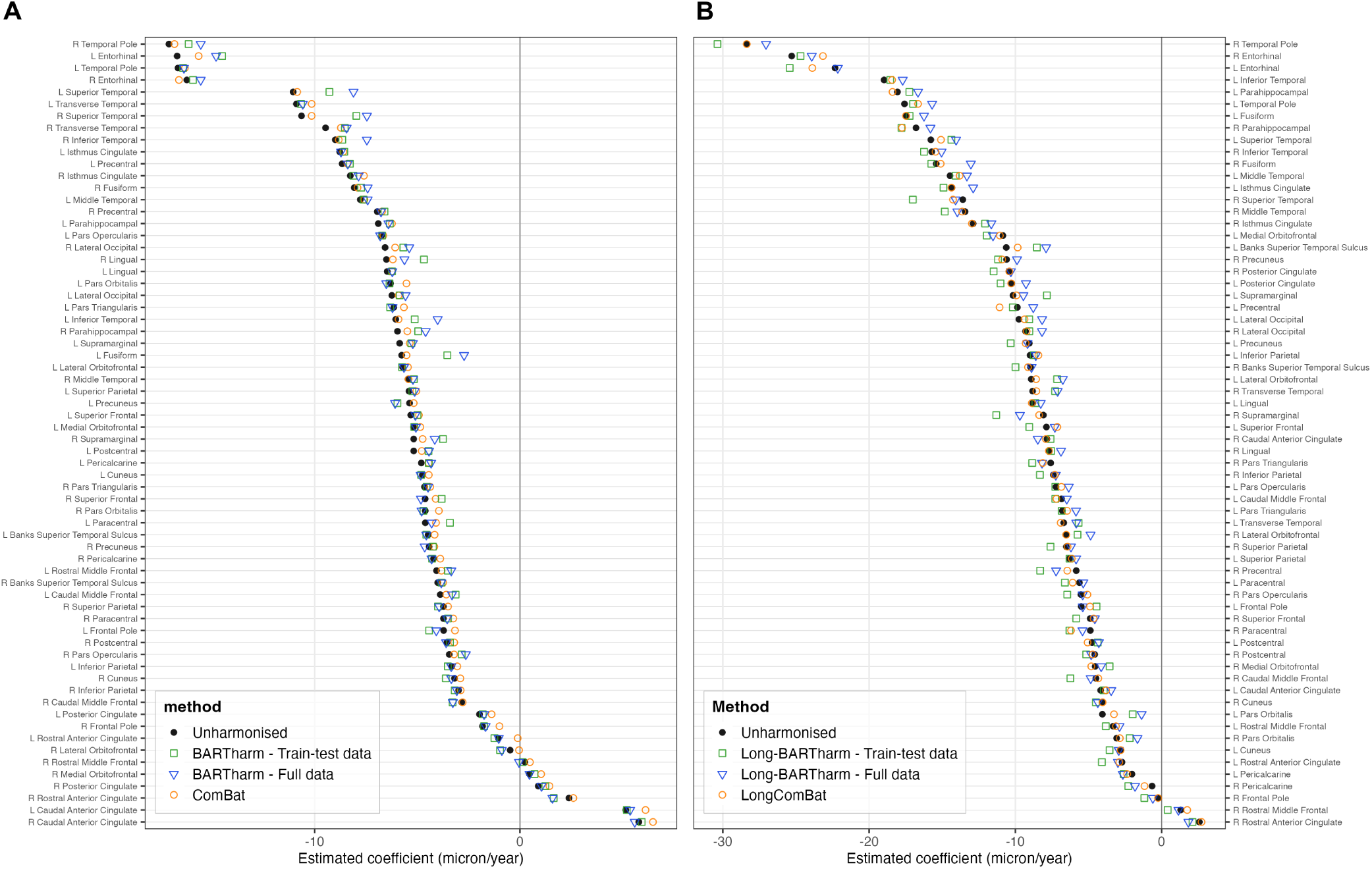
Estimated age- and time-effect coefficients for cortical-thickness IDPs in the combined training/test and full-data harmonisation analyses. A. Cross-sectional age-effect coefficients obtained from linear regression models adjusted for sex, diagnosis, and MMSE score, comparing unharmonised data, ComBat, and BARTharm. B. Longitudinal time-effect coefficients obtained from linear mixed-effects models, representing estimated annual changes in cortical thickness (*µ*m/year) in cognitively normal participants, comparing unharmonised data, LongComBat, and Long-BARTharm. Horizontal reference lines indicate zero effect.

**Figure 9:**
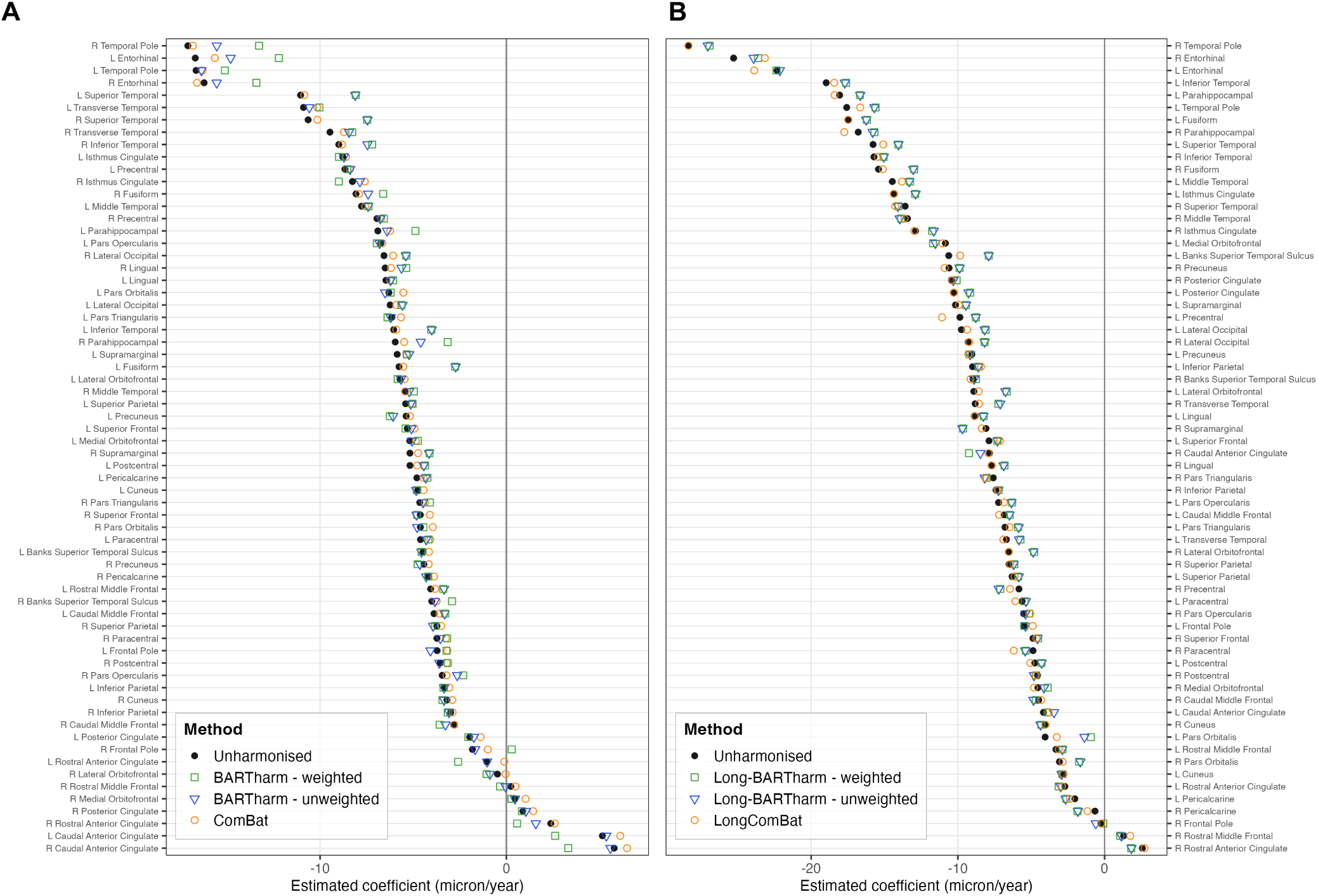
Estimated age- and time-related associations across cortical-thickness IDPs following uncertainty weighting. A. Cross-sectional age-effect coefficients from linear regression models. B. Longitudinal time-effect coefficients from linear mixed-effects models, representing estimated annual changes in cognitively normal participants. Coefficients in both panels are expressed in *µ*m/year. Both panels compare unharmonised data, ComBat or LongComBat, and BARTharm or Long-BARTharm, with and without inverse-variance weighting.

**Figure 10:**
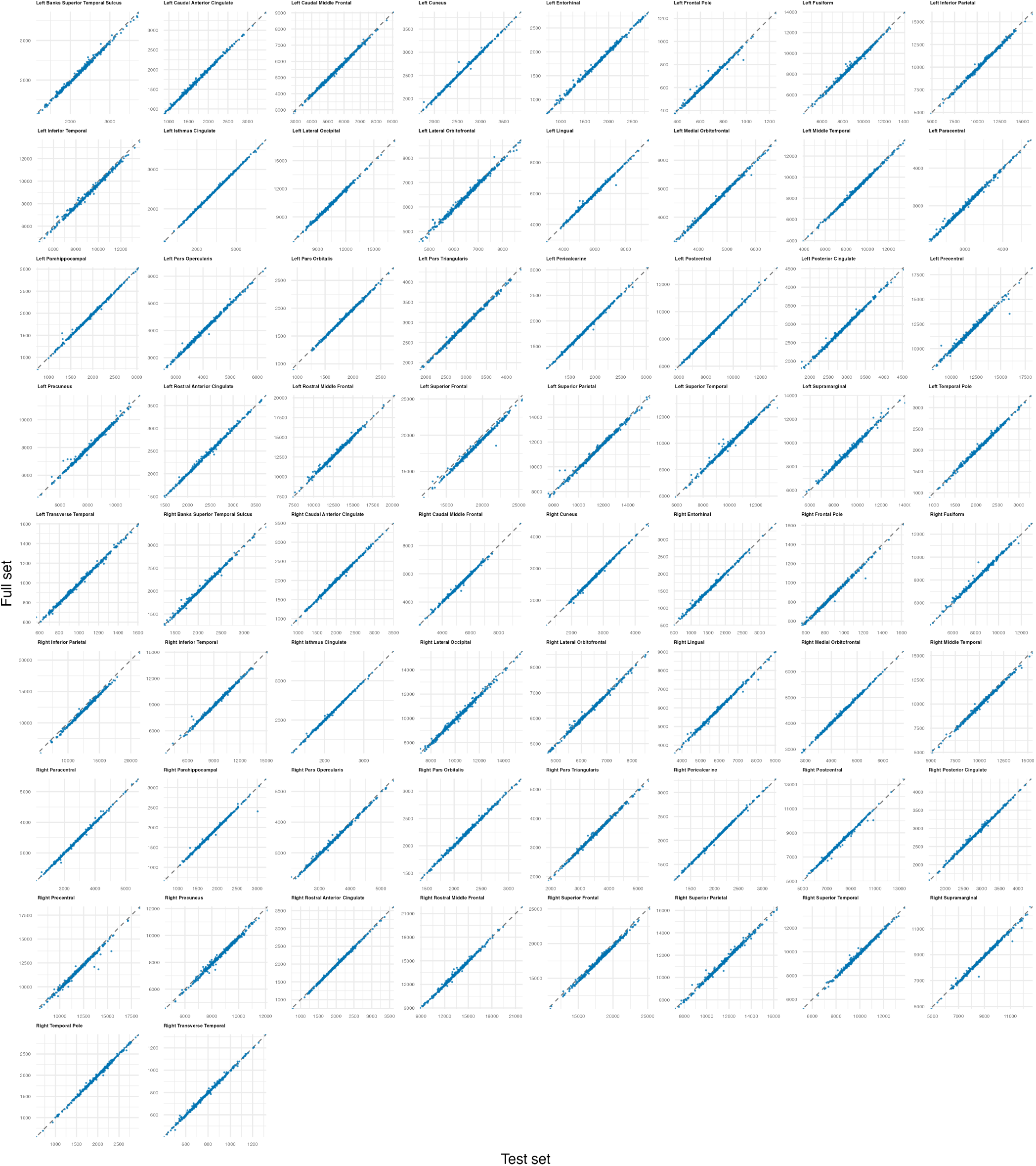
Comparison of posterior predictive and full-data Long-BARTharm harmonisation for cortical-volume IDPs. Scatter plots show the harmonised values obtained for the held-out test set using posterior predictive harmonisation against the corresponding values obtained from full-data harmonisation. The full-data results were restricted to the same participants and time-points included in the held-out test set. The diagonal identity line indicates perfect agreement between the two approaches.

**Figure 11:**
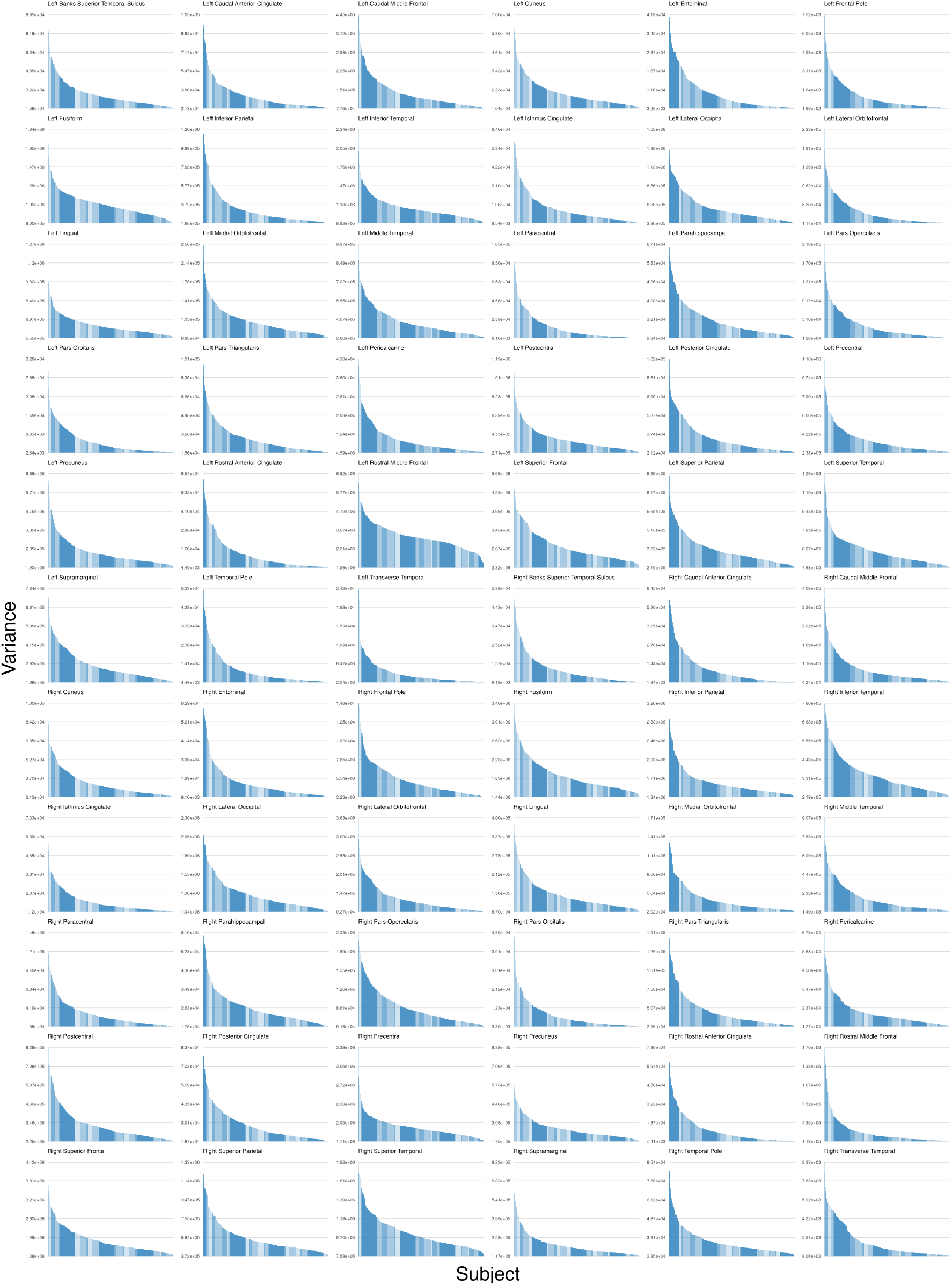
Subject-level posterior uncertainty for cortical-volume IDPs following cross-sectional BARTharm harmonisation. Bars represent the posterior variance associated with individual subjects for each IDP, ordered from highest to lowest posterior variance within each ROI.

**Table 16:**
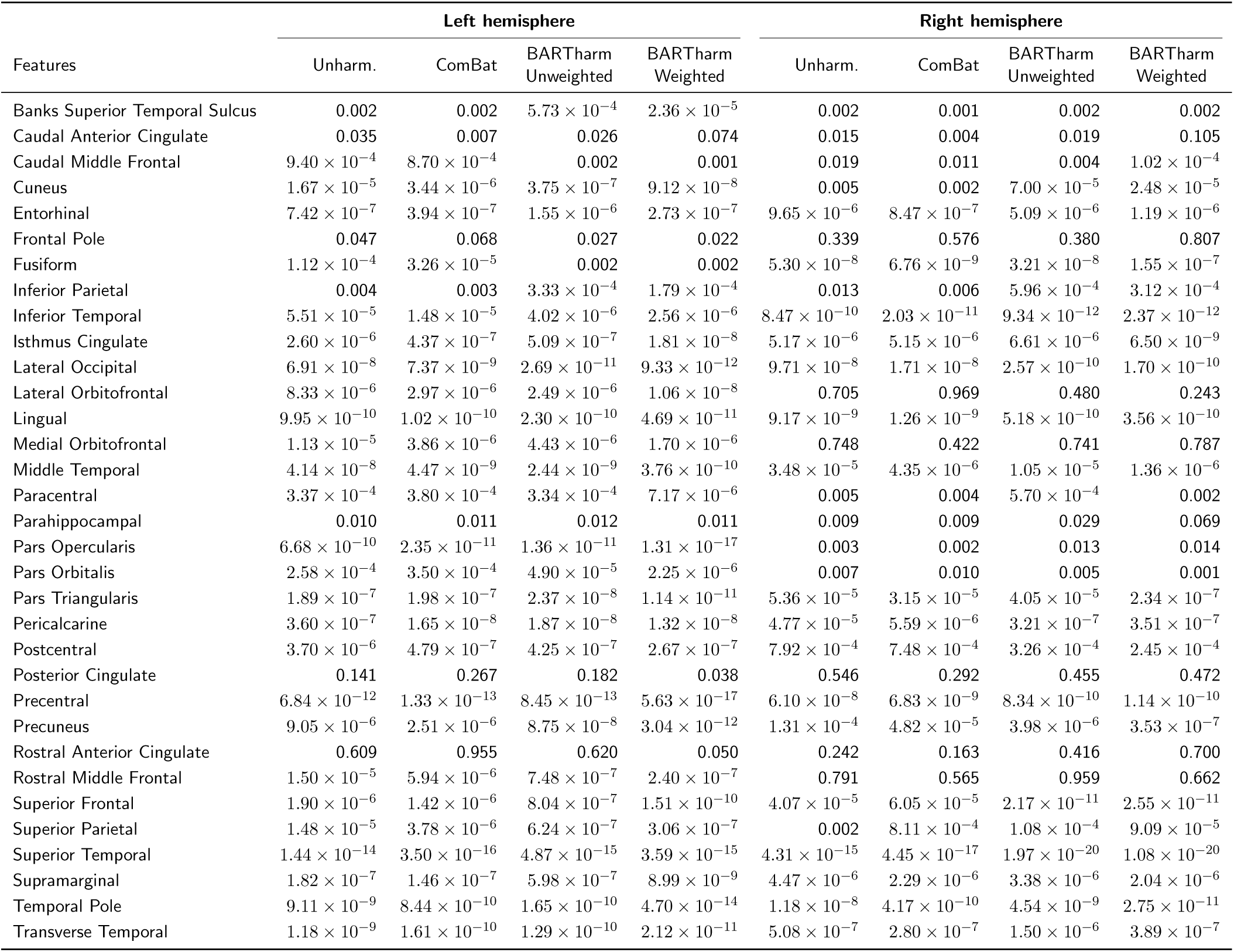
Nominal *p*-values for cross-sectional age-effect coefficients of cortical-thickness features across unharmonised data, ComBat, and unweighted and uncertainty-weighted BARTharm, separately for the left and right hemispheres.

|  | **Left hemisphere** |  |  |  | **Right hemisphere** |  |  |  |
|---|---|---|---|---|---|---|---|---|
| Features | Unharm. | ComBat | BARTharm<br>Unweighted | BARTharm<br>Weighted | Unharm. | ComBat | BARTharm<br>Unweighted | BARTharm<br>Weighted |
| Banks Superior Temporal Sulcus | 0.002 | 0.002 | 5.73 × 10<sup>−4</sup> | 2.36 × 10<sup>−5</sup> | 0.002 | 0.001 | 0.002 | 0.002 |
| Caudal Anterior Cingulate | 0.035 | 0.007 | 0.026 | 0.074 | 0.015 | 0.004 | 0.019 | 0.105 |
| Caudal Middle Frontal | 9.40 × 10<sup>−4</sup> | 8.70 × 10<sup>−4</sup> | 0.002 | 0.001 | 0.019 | 0.011 | 0.004 | 1.02 × 10<sup>−4</sup> |
| Cuneus | 1.67 × 10<sup>−5</sup> | 3.44 × 10<sup>−6</sup> | 3.75 × 10<sup>−7</sup> | 9.12 × 10<sup>−8</sup> | 0.005 | 0.002 | 7.00 × 10<sup>−5</sup> | 2.48 × 10<sup>−5</sup> |
| Entorhinal | 7.42 × 10<sup>−7</sup> | 3.94 × 10<sup>−7</sup> | 1.55 × 10<sup>−6</sup> | 2.73 × 10<sup>−7</sup> | 9.65 × 10<sup>−6</sup> | 8.47 × 10<sup>−7</sup> | 5.09 × 10<sup>−6</sup> | 1.19 × 10<sup>−6</sup> |
| Frontal Pole | 0.047 | 0.068 | 0.027 | 0.022 | 0.339 | 0.576 | 0.380 | 0.807 |
| Fusiform | 1.12 × 10<sup>−4</sup> | 3.26 × 10<sup>−5</sup> | 0.002 | 0.002 | 5.30 × 10<sup>−8</sup> | 6.76 × 10<sup>−9</sup> | 3.21 × 10<sup>−8</sup> | 1.55 × 10<sup>−7</sup> |
| Inferior Parietal | 0.004 | 0.003 | 3.33 × 10<sup>−4</sup> | 1.79 × 10<sup>−4</sup> | 0.013 | 0.006 | 5.96 × 10<sup>−4</sup> | 3.12 × 10<sup>−4</sup> |
| Inferior Temporal | 5.51 × 10<sup>−5</sup> | 1.48 × 10<sup>−5</sup> | 4.02 × 10<sup>−6</sup> | 2.56 × 10<sup>−6</sup> | 8.47 × 10<sup>−10</sup> | 2.03 × 10<sup>−11</sup> | 9.34 × 10<sup>−12</sup> | 2.37 × 10<sup>−12</sup> |
| Isthmus Cingulate | 2.60 × 10<sup>−6</sup> | 4.37 × 10<sup>−7</sup> | 5.09 × 10<sup>−7</sup> | 1.81 × 10<sup>−8</sup> | 5.17 × 10<sup>−6</sup> | 5.15 × 10<sup>−6</sup> | 6.61 × 10<sup>−6</sup> | 6.50 × 10<sup>−9</sup> |
| Lateral Occipital | 6.91 × 10<sup>−8</sup> | 7.37 × 10<sup>−9</sup> | 2.69 × 10<sup>−11</sup> | 9.33 × 10<sup>−12</sup> | 9.71 × 10<sup>−8</sup> | 1.71 × 10<sup>−8</sup> | 2.57 × 10<sup>−10</sup> | 1.70 × 10<sup>−10</sup> |
| Lateral Orbitofrontal | 8.33 × 10<sup>−6</sup> | 2.97 × 10<sup>−6</sup> | 2.49 × 10<sup>−6</sup> | 1.06 × 10<sup>−8</sup> | 0.705 | 0.969 | 0.480 | 0.243 |
| Lingual | 9.95 × 10<sup>−10</sup> | 1.02 × 10<sup>−10</sup> | 2.30 × 10<sup>−10</sup> | 4.69 × 10<sup>−11</sup> | 9.17 × 10<sup>−9</sup> | 1.26 × 10<sup>−9</sup> | 5.18 × 10<sup>−10</sup> | 3.56 × 10<sup>−10</sup> |
| Medial Orbitofrontal | 1.13 × 10<sup>−5</sup> | 3.86 × 10<sup>−6</sup> | 4.43 × 10<sup>−6</sup> | 1.70 × 10<sup>−6</sup> | 0.748 | 0.422 | 0.741 | 0.787 |
| Middle Temporal | 4.14 × 10<sup>−8</sup> | 4.47 × 10<sup>−9</sup> | 2.44 × 10<sup>−9</sup> | 3.76 × 10<sup>−10</sup> | 3.48 × 10<sup>−5</sup> | 4.35 × 10<sup>−6</sup> | 1.05 × 10<sup>−5</sup> | 1.36 × 10<sup>−6</sup> |
| Paracentral | 3.37 × 10<sup>−4</sup> | 3.80 × 10<sup>−4</sup> | 3.34 × 10<sup>−4</sup> | 7.17 × 10<sup>−6</sup> | 0.005 | 0.004 | 5.70 × 10<sup>−4</sup> | 0.002 |
| Parahippocampal | 0.010 | 0.011 | 0.012 | 0.011 | 0.009 | 0.009 | 0.029 | 0.069 |
| Pars Opercularis | 6.68 × 10<sup>−10</sup> | 2.35 × 10<sup>−11</sup> | 1.36 × 10<sup>−11</sup> | 1.31 × 10<sup>−17</sup> | 0.003 | 0.002 | 0.013 | 0.014 |
| Pars Orbitalis | 2.58 × 10<sup>−4</sup> | 3.50 × 10<sup>−4</sup> | 4.90 × 10<sup>−5</sup> | 2.25 × 10<sup>−6</sup> | 0.007 | 0.010 | 0.005 | 0.001 |
| Pars Triangularis | 1.89 × 10<sup>−7</sup> | 1.98 × 10<sup>−7</sup> | 2.37 × 10<sup>−8</sup> | 1.14 × 10<sup>−11</sup> | 5.36 × 10<sup>−5</sup> | 3.15 × 10<sup>−5</sup> | 4.05 × 10<sup>−5</sup> | 2.34 × 10<sup>−7</sup> |
| Pericalcarine | 3.60 × 10<sup>−7</sup> | 1.65 × 10<sup>−8</sup> | 1.87 × 10<sup>−8</sup> | 1.32 × 10<sup>−8</sup> | 4.77 × 10<sup>−5</sup> | 5.59 × 10<sup>−6</sup> | 3.21 × 10<sup>−7</sup> | 3.51 × 10<sup>−7</sup> |
| Postcentral | 3.70 × 10<sup>−6</sup> | 4.79 × 10<sup>−7</sup> | 4.25 × 10<sup>−7</sup> | 2.67 × 10<sup>−7</sup> | 7.92 × 10<sup>−4</sup> | 7.48 × 10<sup>−4</sup> | 3.26 × 10<sup>−4</sup> | 2.45 × 10<sup>−4</sup> |
| Posterior Cingulate | 0.141 | 0.267 | 0.182 | 0.038 | 0.546 | 0.292 | 0.455 | 0.472 |
| Precentral | 6.84 × 10<sup>−12</sup> | 1.33 × 10<sup>−13</sup> | 8.45 × 10<sup>−13</sup> | 5.63 × 10<sup>−17</sup> | 6.10 × 10<sup>−8</sup> | 6.83 × 10<sup>−9</sup> | 8.34 × 10<sup>−10</sup> | 1.14 × 10<sup>−10</sup> |
| Precuneus | 9.05 × 10<sup>−6</sup> | 2.51 × 10<sup>−6</sup> | 8.75 × 10<sup>−8</sup> | 3.04 × 10<sup>−12</sup> | 1.31 × 10<sup>−4</sup> | 4.82 × 10<sup>−5</sup> | 3.98 × 10<sup>−6</sup> | 3.53 × 10<sup>−7</sup> |
| Rostral Anterior Cingulate | 0.609 | 0.955 | 0.620 | 0.050 | 0.242 | 0.163 | 0.416 | 0.700 |
| Rostral Middle Frontal | 1.50 × 10<sup>−5</sup> | 5.94 × 10<sup>−6</sup> | 7.48 × 10<sup>−7</sup> | 2.40 × 10<sup>−7</sup> | 0.791 | 0.565 | 0.959 | 0.662 |
| Superior Frontal | 1.90 × 10<sup>−6</sup> | 1.42 × 10<sup>−6</sup> | 8.04 × 10<sup>−7</sup> | 1.51 × 10<sup>−10</sup> | 4.07 × 10<sup>−5</sup> | 6.05 × 10<sup>−5</sup> | 2.17 × 10<sup>−11</sup> | 2.55 × 10<sup>−11</sup> |
| Superior Parietal | 1.48 × 10<sup>−5</sup> | 3.78 × 10<sup>−6</sup> | 6.24 × 10<sup>−7</sup> | 3.06 × 10<sup>−7</sup> | 0.002 | 8.11 × 10<sup>−4</sup> | 1.08 × 10<sup>−4</sup> | 9.09 × 10<sup>−5</sup> |
| Superior Temporal | 1.44 × 10<sup>−14</sup> | 3.50 × 10<sup>−16</sup> | 4.87 × 10<sup>−15</sup> | 3.59 × 10<sup>−15</sup> | 4.31 × 10<sup>−15</sup> | 4.45 × 10<sup>−17</sup> | 1.97 × 10<sup>−20</sup> | 1.08 × 10<sup>−20</sup> |
| Supramarginal | 1.82 × 10<sup>−7</sup> | 1.46 × 10<sup>−7</sup> | 5.98 × 10<sup>−7</sup> | 8.99 × 10<sup>−9</sup> | 4.47 × 10<sup>−6</sup> | 2.29 × 10<sup>−6</sup> | 3.38 × 10<sup>−6</sup> | 2.04 × 10<sup>−6</sup> |
| Temporal Pole | 9.11 × 10<sup>−9</sup> | 8.44 × 10<sup>−10</sup> | 1.65 × 10<sup>−10</sup> | 4.70 × 10<sup>−14</sup> | 1.18 × 10<sup>−8</sup> | 4.17 × 10<sup>−10</sup> | 4.54 × 10<sup>−9</sup> | 2.75 × 10<sup>−11</sup> |
| Transverse Temporal | 1.18 × 10<sup>−9</sup> | 1.61 × 10<sup>−10</sup> | 1.29 × 10<sup>−10</sup> | 2.12 × 10<sup>−11</sup> | 5.08 × 10<sup>−7</sup> | 2.80 × 10<sup>−7</sup> | 1.50 × 10<sup>−6</sup> | 3.89 × 10<sup>−7</sup> |

**Table 17:** Estimated longitudinal time-effect coefficients for cortical-thickness features across unharmonised data, LongComBat, and unweighted and uncertainty-weighted Long-BARTharm, separately for the left and right hemispheres. Values are expressed in *µ*m/year.

|  | **Left hemisphere** |  |  |  | **Right hemisphere** |  |  |  |
|---|---|---|---|---|---|---|---|---|
| Features | Unharm. | LongComBat | Long-BARTharm<br>Unweighted | Long-BARTharm<br>Weighted | Unharm. | LongComBat | Long-BARTharm<br>Unweighted | Long-BARTharm<br>Weighted |
| Banks Superior Temporal Sulcus | -10.62 | -9.84 | -7.90 | -7.90 | -8.97 | -9.14 | -8.90 | -8.76 |
| Caudal Anterior Cingulate | -4.17 | -3.80 | -3.45 | -3.90 | -7.84 | -7.92 | -8.47 | -9.24 |
| Caudal Middle Frontal | -6.85 | -7.20 | -6.49 | -6.46 | -4.49 | -4.33 | -4.85 | -4.69 |
| Cuneus | -2.83 | -2.80 | -2.93 | -2.88 | -4.07 | -4.04 | -4.37 | -4.32 |
| Entorhinal | -22.32 | -23.88 | -22.14 | -22.36 | -25.29 | -23.16 | -23.92 | -23.62 |
| Frontal Pole | -5.50 | -4.90 | -5.39 | -5.48 | -0.22 | -0.25 | -0.60 | -0.09 |
| Fusiform | -17.45 | -17.48 | -16.25 | -16.21 | -15.41 | -15.11 | -13.04 | -12.98 |
| Inferior Parietal | -9.00 | -8.43 | -8.61 | -8.59 | -7.40 | -7.31 | -7.24 | -7.22 |
| Inferior Temporal | -18.98 | -18.43 | -17.70 | -17.62 | -15.72 | -15.46 | -15.05 | -15.06 |
| Isthmus Cingulate | -14.33 | -14.37 | -12.89 | -12.91 | -12.89 | -12.97 | -11.65 | -11.75 |
| Lateral Occipital | -9.76 | -9.37 | -8.17 | -8.11 | -9.30 | -9.23 | -8.18 | -8.17 |
| Lateral Orbitofrontal | -8.91 | -8.58 | -6.76 | -6.67 | -6.54 | -6.53 | -4.86 | -4.80 |
| Lingual | -8.81 | -8.88 | -8.27 | -8.24 | -7.68 | -7.71 | -6.88 | -6.82 |
| Medial Orbitofrontal | -10.85 | -11.06 | -11.53 | -11.69 | -4.55 | -4.81 | -4.13 | -3.88 |
| Middle Temporal | -14.47 | -13.81 | -13.31 | -13.26 | -13.44 | -13.64 | -13.96 | -13.86 |
| Paracentral | -5.62 | -6.08 | -5.36 | -5.45 | -4.88 | -6.20 | -5.41 | -5.36 |
| Parahippocampal | -18.06 | -18.40 | -16.66 | -16.68 | -16.79 | -17.75 | -15.81 | -15.68 |
| Pars Opercularis | -7.23 | -6.86 | -6.35 | -6.30 | -5.52 | -5.08 | -5.40 | -5.15 |
| Pars Orbitalis | -4.05 | -3.26 | -1.37 | -0.93 | -3.08 | -2.86 | -1.66 | -1.74 |
| Pars Triangularis | -6.79 | -6.46 | -5.86 | -5.90 | -7.59 | -8.13 | -8.18 | -8.04 |
| Pericalcarine | -2.03 | -2.36 | -2.66 | -2.61 | -0.65 | -1.17 | -1.82 | -1.84 |
| Postcentral | -4.76 | -5.07 | -4.30 | -4.30 | -4.58 | -4.72 | -4.82 | -4.67 |
| Posterior Cingulate | -10.24 | -10.29 | -9.28 | -9.16 | -10.44 | -10.41 | -10.31 | -10.08 |
| Precentral | -9.86 | -11.08 | -8.78 | -8.77 | -5.84 | -6.45 | -7.22 | -7.11 |
| Precuneus | -9.05 | -9.30 | -9.17 | -9.19 | -10.60 | -10.90 | -9.89 | -9.88 |
| Rostral Anterior Cingulate | -2.71 | -2.90 | -2.99 | -3.13 | 2.58 | 2.72 | 1.83 | 1.81 |
| Rostral Middle Frontal | -3.32 | -3.13 | -2.88 | -2.88 | 1.30 | 1.75 | 1.15 | 1.06 |
| Superior Frontal | -7.88 | -7.14 | -7.30 | -7.31 | -4.89 | -4.65 | -4.57 | -4.53 |
| Superior Parietal | -6.30 | -6.14 | -5.85 | -5.89 | -6.52 | -6.44 | -6.20 | -6.15 |
| Superior Temporal | -15.78 | -15.09 | -14.05 | -14.06 | -13.59 | -14.26 | -14.09 | -14.06 |
| Supramarginal | -10.17 | -9.93 | -9.45 | -9.45 | -8.08 | -8.38 | -9.69 | -9.65 |
| Temporal Pole | -17.58 | -16.65 | -15.69 | -15.61 | -28.34 | -28.37 | -27.05 | -26.92 |
| Transverse Temporal | -6.69 | -6.90 | -5.84 | -5.73 | -8.82 | -8.57 | -7.11 | -7.25 |

At the nominal threshold, weighted and unweighted Long-BARTharm identified similar numbers of significant longitudinal time effects to LongComBat, with all harmonisation approaches exceeding the unharmonised data. Following Bonferroni correction, weighted Long-BARTharm retained the most significant IDPs, followed by unweighted Long-BARTharm and LongComBat (Supplementary Table 5). Both Long-BARTharm approaches generally yielded smaller *p*-values than LongComBat, with weighted

Long-BARTharm outperforming its unweighted counterpart in 45 of 46 shared significant IDPs at the nominal threshold and all 21 after Bonferroni correction (Figure 6B). Estimated time-effect coefficients were similar across harmonisation approaches (Supplementary Figure 9B). Regional effect estimates and *p*-values are provided in Supplementary Tables 17 and 18.

**Table 18:** Kenward–Roger *p*-values for longitudinal time-effect coefficients of cortical-thickness features across unharmonised data, LongComBat, and unweighted and uncertainty-weighted Long-BARTharm, separately for the left and right hemispheres.

|  | Left hemisphere |  |  |  | Right hemisphere |  |  |  |
|---|---|---|---|---|---|---|---|---|
| Features | Unharm. | LongComBat | Long-BARTharm<br>Unweighted | Long-BARTharm<br>Weighted | Unharm. | LongComBat | Long-BARTharm<br>Unweighted | Long-BARTharm<br>Weighted |
| Banks Superior Temporal Sulcus | 0.006 | 0.009 | 0.019 | 0.017 | 0.011 | 0.007 | 0.007 | 0.006 |
| Caudal Anterior Cingulate | 0.324 | 0.372 | 0.399 | 0.318 | 0.098 | 0.092 | 0.055 | 0.026 |
| Caudal Middle Frontal | 0.069 | 0.049 | 0.044 | 0.043 | 0.222 | 0.215 | 0.114 | 0.118 |
| Cuneus | 0.173 | 0.176 | 0.137 | 0.138 | 0.048 | 0.045 | 0.022 | 0.020 |
| Entorhinal | 8.57 × 10<sup>−4</sup> | 2.65 × 10<sup>−4</sup> | 4.56 × 10<sup>−4</sup> | 2.68 × 10<sup>−4</sup> | 5.1 × 10<sup>−4</sup> | 9.85 × 10<sup>−4</sup> | 3.79 × 10<sup>−4</sup> | 1.61 × 10<sup>−4</sup> |
| Frontal Pole | 0.338 | 0.385 | 0.312 | 0.284 | 0.964 | 0.959 | 0.893 | 0.982 |
| Fusiform | 9.61 × 10<sup>−6</sup> | 3.68 × 10<sup>−6</sup> | 2.95 × 10<sup>−6</sup> | 1.62 × 10<sup>−6</sup> | 2.39 × 10<sup>−4</sup> | 1.49 × 10<sup>−4</sup> | 2.46 × 10<sup>−4</sup> | 1.88 × 10<sup>−4</sup> |
| Inferior Parietal | 0.008 | 0.008 | 0.002 | 0.002 | 0.018 | 0.014 | 0.009 | 0.008 |
| Inferior Temporal | 1.03 × 10<sup>−5</sup> | 8.45 × 10<sup>−6</sup> | 5.91 × 10<sup>−6</sup> | 3.5 × 10<sup>−6</sup> | 1.19 × 10<sup>−4</sup> | 5.37 × 10<sup>−5</sup> | 3.39 × 10<sup>−5</sup> | 2.08 × 10<sup>−5</sup> |
| Isthmus Cingulate | 4.32 × 10<sup>−4</sup> | 2.26 × 10<sup>−4</sup> | 4.94 × 10<sup>−4</sup> | 3 × 10<sup>−4</sup> | 8.5 × 10<sup>−4</sup> | 3.08 × 10<sup>−4</sup> | 0.001 | 4.49 × 10<sup>−4</sup> |
| Lateral Occipital | 7.02 × 10<sup>−4</sup> | 0.001 | 0.002 | 0.002 | 0.002 | 0.001 | 0.002 | 0.001 |
| Lateral Orbitofrontal | 0.005 | 0.005 | 0.022 | 0.021 | 0.060 | 0.048 | 0.119 | 0.117 |
| Lingual | 2.23 × 10<sup>−5</sup> | 1.24 × 10<sup>−5</sup> | 1.17 × 10<sup>−5</sup> | 6.45 × 10<sup>−6</sup> | 3.77 × 10<sup>−4</sup> | 2.53 × 10<sup>−4</sup> | 3.95 × 10<sup>−4</sup> | 3.12 × 10<sup>−4</sup> |
| Medial Orbitofrontal | 7.4 × 10<sup>−4</sup> | 4.15 × 10<sup>−4</sup> | 1.47 × 10<sup>−4</sup> | 7.42 × 10<sup>−5</sup> | 0.181 | 0.143 | 0.186 | 0.194 |
| Middle Temporal | 2.43 × 10<sup>−4</sup> | 2.73 × 10<sup>−4</sup> | 1.14 × 10<sup>−4</sup> | 7.56 × 10<sup>−5</sup> | 4.49 × 10<sup>−4</sup> | 2.04 × 10<sup>−4</sup> | 3.53 × 10<sup>−5</sup> | 2.21 × 10<sup>−5</sup> |
| Paracentral | 0.102 | 0.073 | 0.068 | 0.061 | 0.116 | 0.049 | 0.050 | 0.047 |
| Parahippocampal | 3.62 × 10<sup>−5</sup> | 1.55 × 10<sup>−5</sup> | 3.1 × 10<sup>−5</sup> | 1.5 × 10<sup>−5</sup> | 3.96 × 10<sup>−4</sup> | 7.08 × 10<sup>−5</sup> | 1.28 × 10<sup>−4</sup> | 8.89 × 10<sup>−5</sup> |
| Pars Opercularis | 0.024 | 0.028 | 0.020 | 0.019 | 0.094 | 0.110 | 0.065 | 0.073 |
| Pars Orbitalis | 0.324 | 0.402 | 0.704 | 0.788 | 0.409 | 0.426 | 0.623 | 0.593 |
| Pars Triangularis | 0.045 | 0.050 | 0.042 | 0.036 | 0.016 | 0.008 | 0.003 | 0.003 |
| Pericalcarine | 0.347 | 0.263 | 0.190 | 0.188 | 0.769 | 0.593 | 0.372 | 0.360 |
| Postcentral | 0.050 | 0.036 | 0.037 | 0.035 | 0.057 | 0.043 | 0.017 | 0.019 |
| Posterior Cingulate | 0.002 | 0.002 | 0.003 | 0.002 | 0.005 | 0.003 | 0.003 | 0.002 |
| Precentral | 0.007 | 0.003 | 0.001 | 0.001 | 0.098 | 0.066 | 0.015 | 0.015 |
| Precuneus | 0.002 | 7.53 × 10<sup>−4</sup> | 1.88 × 10<sup>−4</sup> | 1.24 × 10<sup>−4</sup> | 8.21 × 10<sup>−5</sup> | 2.62 × 10<sup>−5</sup> | 2.08 × 10<sup>−5</sup> | 1.14 × 10<sup>−5</sup> |
| Rostral Anterior Cingulate | 0.498 | 0.457 | 0.450 | 0.408 | 0.545 | 0.514 | 0.650 | 0.636 |
| Rostral Middle Frontal | 0.287 | 0.297 | 0.285 | 0.279 | 0.658 | 0.535 | 0.642 | 0.660 |
| Superior Frontal | 0.023 | 0.028 | 0.009 | 0.008 | 0.122 | 0.124 | 0.073 | 0.071 |
| Superior Parietal | 0.032 | 0.027 | 0.016 | 0.015 | 0.020 | 0.014 | 0.006 | 0.006 |
| Superior Temporal | 1.94 × 10<sup>−6</sup> | 1.82 × 10<sup>−6</sup> | 3.16 × 10<sup>−7</sup> | 6.64 × 10<sup>−8</sup> | 1.94 × 10<sup>−5</sup> | 6.64 × 10<sup>−6</sup> | 3.16 × 10<sup>−7</sup> | 8.59 × 10<sup>−8</sup> |
| Supramarginal | 0.003 | 0.002 | 6.15 × 10<sup>−4</sup> | 4.58 × 10<sup>−4</sup> | 0.016 | 0.008 | 6.37 × 10<sup>−4</sup> | 4.73 × 10<sup>−4</sup> |
| Temporal Pole | 0.002 | 0.003 | 0.002 | 0.002 | 8.79 × 10<sup>−5</sup> | 4.52 × 10<sup>−5</sup> | 3.09 × 10<sup>−5</sup> | 1.96 × 10<sup>−5</sup> |
| Transverse Temporal | 0.073 | 0.062 | 0.090 | 0.089 | 0.024 | 0.030 | 0.048 | 0.045 |

The distributions of normalised posterior variance across the 66 cortical-volume IDPs for cross-sectional BARTharm and Long-BARTharm are illustrated in Figure 5. In both analyses, posterior uncertainty varied across regions, with most IDPs exhibiting relatively concentrated distributions and a smaller number showing greater dispersion. These distributions demonstrate regional variability in the uncertainty associated with the harmonised estimates.

Posterior uncertainty for cortical-volume IDPs is illustrated for cross-sectional BARTharm and Long-BARTharm in Supplementary Figures 11 and 12, respectively. Each observation represents a subject in the cross-sectional analysis and a subject–timepoint in the longitudinal analysis. Within individual IDPs, posterior variance varied across observations, illustrating differences in uncertainty associated with the harmonised estimates.

**Figure 12:**
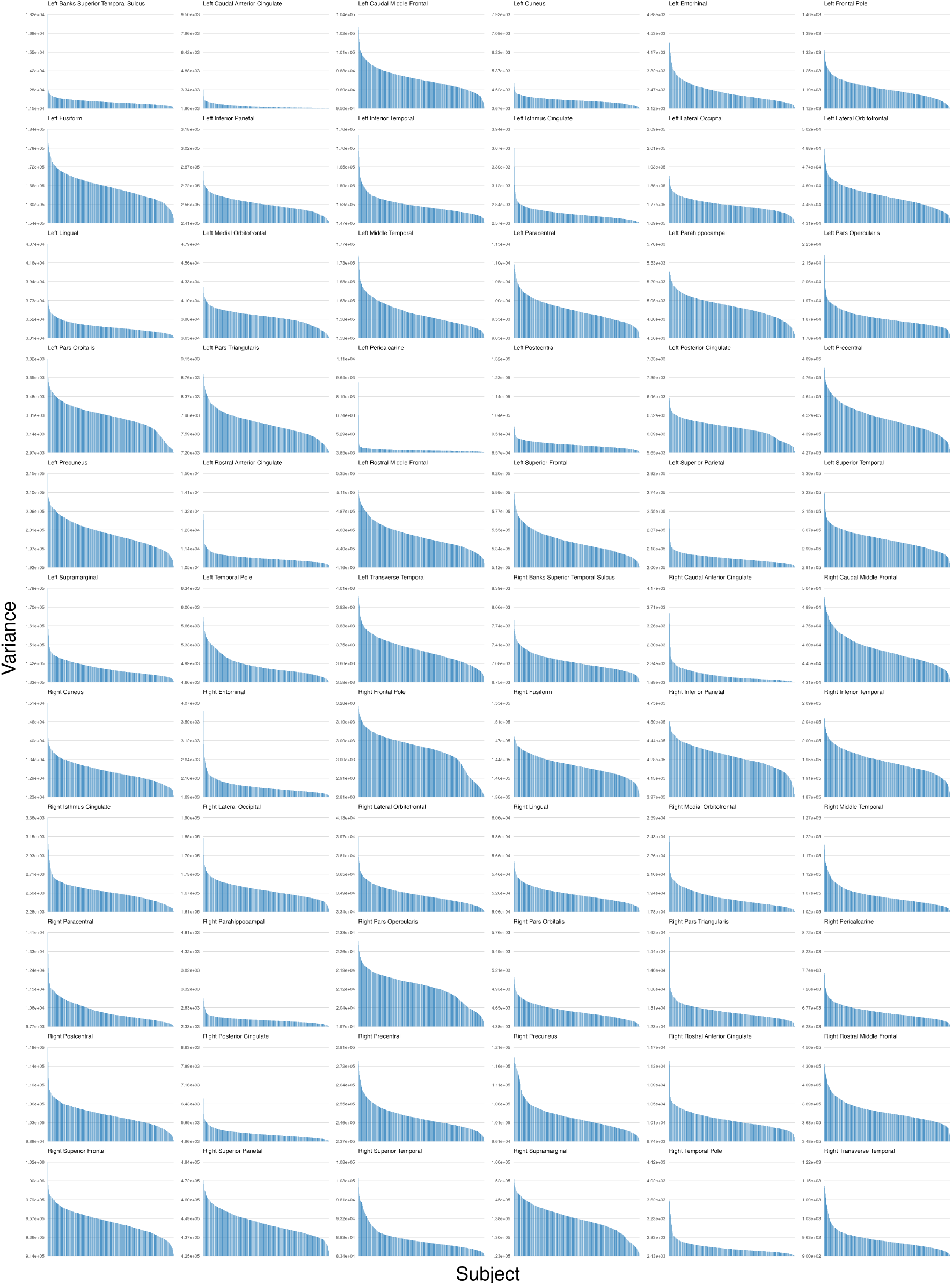
Posterior uncertainty for cortical-volume IDPs following Long-BARTharm harmonisation. Each bar represents the posterior variance for a subject–timepoint, with bars ordered from highest to lowest variance within each ROI.

## 4 Discussion

Although conventional neuroimaging harmonisation approaches have primarily addressed acquisition-related variability in cross-sectional data, longitudinal studies introduce additional methodological challenges, particularly the need to account for within-subject dependence while preserving biologically meaningful changes over time (Beer et al., 2020). In parallel, the accumulation of new observations creates a practical need for harmonisation procedures that can be applied without repeatedly fitting computationally demanding models. Moreover, uncertainty introduced during harmonisation is often not explicitly considered in subsequent statistical analyses (Reynolds et al., 2023). To address these challenges, we extend the original BARTharm methodology into a broader Bayesian nonparametric framework encompassing longitudinal harmonisation, posterior predictive harmonisation, and uncertainty-aware downstream inference.

Longitudinal neuroimaging data require harmonisation approaches that account for dependence between repeated observations while distinguishing acquisition-related variability from biological changes over time. Unlike cross-sectional measurements, repeated observations from the same individual are correlated, and failing to account for this dependence may affect estimates and inference for longitudinal biological effects (Beer et al., 2020). Long-BARTharm addresses these challenges by incorporating subject-specific random intercepts and slopes, allowing individuals to differ in their baseline measurements and linear rates of change while retaining BART’s flexibility to model nonlinear population-level temporal relationships and interactions. Building on the original BARTharm formulation, the longitudinal framework also incorporates continuous IQMs to represent acquisition-related variability beyond categorical scanner identifiers, while accommodating scanner-specific residual variance differences when scanner information is available (Prevot et al., 2025).

In both simulation settings, Long-BARTharm recovers the known scanner-free outcomes more accurately than LongComBat and leaves less association between the harmonised outcomes and the simulated scanner effect. This is consistent with cross-sectional work showing that IQMs provide information about the quality of individual scans beyond that available from scanner identifiers alone (Prevot et al., 2025). Additionally, in the longitudinal setting, a subject can return to the same scanner at each visit while the quality of their scans varies over time. LongComBat uses the scanner identifier to estimate scanner effects, so visits acquired on the same scanner share the same estimated shift and scale. Long-BARTharm allows the scanner contribution to vary with the IQMs measured at each visit, which also may help distinguish acquisition-related changes between visits from changes in the subject’s underlying trajectory. LongComBat does on average estimate the residual scale multipliers more accurately in the semi-synthetic setting. However, this advantage is concentrated among scanners with fewer than 10 scans, while Long-BARTharm gives more accurate estimates for scanners with at least 20 scans. Hence, the number of scans available per scanner seems to be important when interpreting the residual scale multipliers estimated by Long-BARTharm.

In the longitudinal ADNI analysis, Long-BARTharm reduced scanner-specific residual variance heterogeneity, with lower heterogeneity than LongComBat across most cortical regions. By combining flexible IQM-based modelling of acquisition-related mean effects with scanner-specific residual scale adjustments, Long-BARTharm provides a means of addressing distinct sources of technical variability within a longitudinal framework. Beyond reducing scanner-related variability, an important objective of longitudinal harmonisation is to preserve biologically meaningful changes over time. In the ADNI analysis, Long-BARTharm generally yielded stronger statistical evidence for longitudinal cortical-thickness associations than LongComBat, as reflected by smaller time-effect p-values across a greater number of cortical regions. Given the established association between ageing and cortical thinning (Di Biase et al., 2023), these findings are consistent with Long-BARTharm retaining biologically relevant longitudinal variation while reducing acquisition-related effects. By modelling acquisition-related variability through continuous IQMs while retaining time-dependent biological variation, Long-BARTharm may reduce residual technical noise that could otherwise obscure longitudinal associations in downstream analyses.

Posterior predictive harmonisation extends the practical applicability of BARTharm by allowing newly acquired observations to be harmonised using an existing fitted model, without repeating the computationally demanding MCMC estimation procedure. This is particularly relevant for large-scale or evolving neuroimaging studies, where additional participants or follow-up measurements may become available after the initial harmonisation analysis. Beyond reducing the need for repeated model fitting, this approach also provides flexibility in how newly harmonised observations are incorporated into downstream analyses. Specifically, these observations can either be analysed independently or combined with previously harmonised data, allowing researchers to investigate newly acquired measurements separately or as part of an expanding dataset. In both settings, an important consideration is whether posterior predictive harmonisation yields downstream results comparable to those obtained through full-data harmonisation, in which the model is fitted using all available observations. In the cross-sectional analysis, posterior predictive BARTharm yielded age-related associations broadly consistent with those obtained through full-data harmonisation when analyses were restricted to newly harmonised participants. Similarly, combining these observations with previously harmonised data produced comparable age-related associations. Notably, BARTharm retained its tendency to yield stronger statistical evidence for age-related associations than ComBat under both posterior predictive and full-data harmonisation. In the longitudinal analysis, posterior predictive Long-BARTharm was applied to additional follow-up observations from participants already represented during model fitting. When these observations were combined with previously harmonised measurements, the resulting longitudinal time-related associations were broadly consistent with those obtained through full-data harmonisation. Furthermore, the close agreement between posterior predictive and full-data harmonised cortical-volume measurements suggests that the fitted model can provide similar acquisition-related adjustments for subsequent observations. Together, these findings support the application of Long-BARTharm to evolving longitudinal datasets as additional follow-up measurements become available. Although these findings support the practical use of posterior predictive harmonisation, its applicability depends on the characteristics represented during model fitting. In particular, new observations should have biological covariates and IQMs within the ranges covered by the training data, since the fitted model may not reliably represent relationships outside these ranges. When scanner-specific residual scale adjustments are included, new observations must originate from scanner configurations represented during training, as posterior draws of the corresponding variance parameters are unavailable for previously unseen configurations. Categorical covariates must likewise have levels represented in the training data. An additional restriction applies to Long-BARTharm, where posterior predictive harmonisation is currently limited to follow-up observations from participants represented during model fitting, as the procedure reuses their previously estimated subject-specific random intercepts and slopes. Consequently, observations that do not satisfy these conditions may require updating or refitting the harmonisation model.

Harmonisation involves estimating acquisition-related effects, introducing uncertainty into the resulting harmonised measurements. Importantly, this uncertainty may vary across observations and brain regions, such that harmonised measurements are not necessarily estimated with equal precision. Our posterior variance analyses demonstrated this heterogeneity in both cross-sectional BARTharm and Long-BARTharm, revealing differences in harmonisation uncertainty across cortical regions and individual observations. Such heterogeneity is relevant to downstream statistical analyses, where harmonised measurements are typically treated as fixed quantities without explicitly accounting for uncertainty introduced during harmonisation. In particular, conventional unweighted regression does not distinguish observations according to their estimated harmonisation precision, raising the question of whether incorporating this information could influence downstream inference. To account for differences in harmonisation precision, we incorporate observation-specific inverse posterior-variance weights into downstream regression analyses. In both cross-sectional and longitudinal settings, uncertainty weighting generally yielded smaller p-values for age-related and time-related associations, respectively, compared with unweighted BARTharm. Notably, the estimated biological effect coefficients remained broadly similar between weighted and unweighted analyses, suggesting that the differences in statistical significance may reflect changes in the estimated precision of these associations rather than substantial changes in their magnitudes. These findings nevertheless require cautious interpretation. Smaller p-values following uncertainty weighting do not necessarily establish improved statistical power or more reliable inference, as the validity of the resulting statistical tests depends on the assumptions underlying the regression and weighting procedures. Moreover, posterior harmonisation variance quantifies uncertainty associated with the harmonised estimates but does not necessarily represent the total residual variance relevant to downstream regression. Thus, inverse posterior-variance weighting provides a means of incorporating harmonisation precision into subsequent analyses, rather than fully propagating harmonisation uncertainty or guaranteeing improved inferential performance.

Several practical considerations remain important when applying BARTharm. As discussed in the original cross-sectional framework (Prevot et al., 2025), IQM selection requires particular attention because some metrics may reflect genuine anatomical or disease-related variation rather than exclusively acquisition-related effects. This is especially relevant in longitudinal studies, where biological changes over time may influence tissue-derived IQMs. In our ADNI application, IQMs associated with tissue composition and intracranial volume fractions were excluded following an assessment of their relationships with biological and clinical characteristics. In addition, the longitudinal formulation assumes linear subject-specific deviations from the population-level trajectory, although the biological BART component can accommodate nonlinear population-level relationships with time. More complex individual trajectories may therefore require alternative random-effects structures. Finally, although posterior predictive harmonisation reduces the computational demands of processing additional observations, initial model fitting still requires MCMC sampling and remains more computationally intensive than conventional location-scale approaches. Nevertheless, because IDPs are modelled independently, computation can be parallelised across brain regions without requiring specialised hardware.

## 5 Conclusions

In this work, we extended the original BARTharm framework to address three complementary challenges in neuroimaging harmonisation: accommodating longitudinal measurements, harmonising newly acquired observations using previously fitted models, and incorporating harmonisation uncertainty into downstream statistical analyses. The proposed Long-BARTharm formulation accounts for subject-specific trajectories while retaining the flexibility of Bayesian Additive Regression Trees to model complex biological and acquisition-related effects. Across the evaluated synthetic, semi-synthetic, and real-data settings, Long-BARTharm demonstrated effective reduction of scanner-related variability while retaining biologically relevant longitudinal associations, with improvements over LongComBat in several evaluated settings. Beyond longitudinal harmonisation, we developed posterior predictive procedures for both cross-sectional BARTharm and Long-BARTharm, enabling new observations to be harmonised without repeating the full MCMC estimation procedure. The observed agreement with full-data harmonisation supports the applicability of these procedures to evolving neuroimaging datasets. Furthermore, the posterior distributions revealed heterogeneous harmonisation uncertainty across brain regions and individual observations, while incorporating this information through inverse-variance weighting influenced the statistical significance of downstream biological associations.

Overall, these developments broaden the applicability of BARTharm beyond conventional cross-sectional harmonisation, combining flexible acquisition-effect modelling with longitudinal inference, posterior prediction, and uncertainty-aware downstream analysis. The resulting framework provides additional capabilities for harmonising heterogeneous and evolving neuroimaging datasets while accounting for biological variation and the uncertainty associated with harmonised measurements.

## Supplementary Material

### 1 Long-BARTharm model and posterior inference

#### 1.1 Model specification

Ignoring the IDP subscript *v*, for subject *i*, visit *j*, scanner/site *s*(*i, j*), *T_ij_* = (1*, t_ij_*), *α_i_* = (*α_i_*_1_*, α_i_*_2_)*^⊤^* and random-effect component *d* = 1, 2, we model

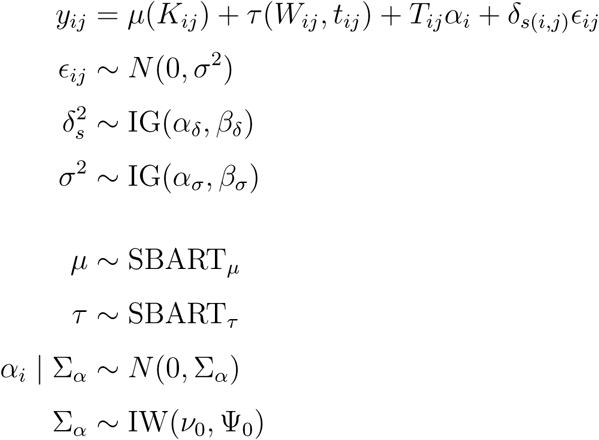

#### 1.2 Posterior updates

Here, *N* denotes the number of subjects, *n_i_* denotes the number of visits for subject *i*, *n_s_* denotes the number of observations (*i, j*) acquired at scanner/site *s*, and *n* = Σ*_i_ n_i_* = Σ*_s_ n_s_* denotes the total number of observations.

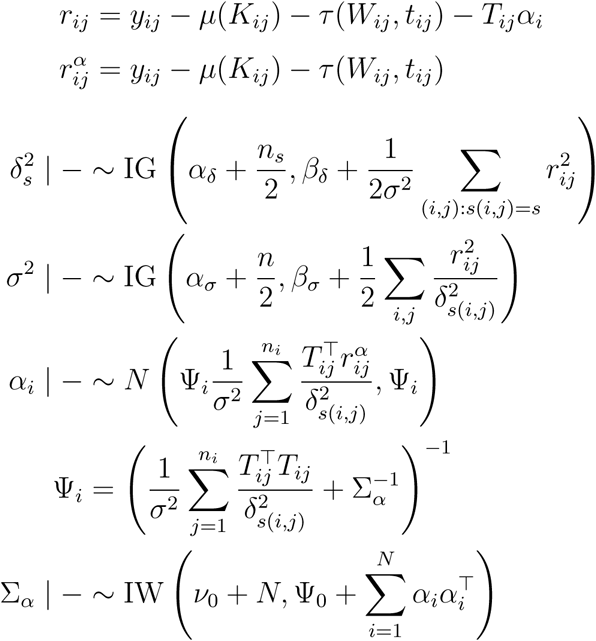

#### 1.3 Derivations of the full conditional distributions

**Global residual variance** *σ*^2^

Relevant conditional density and prior

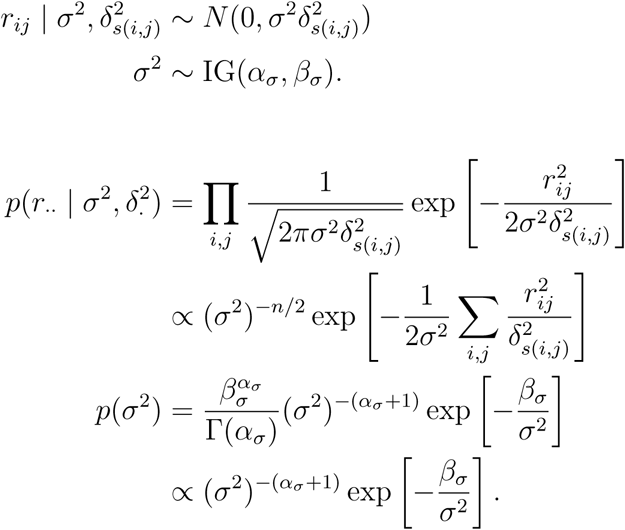

Conditional posterior distribution

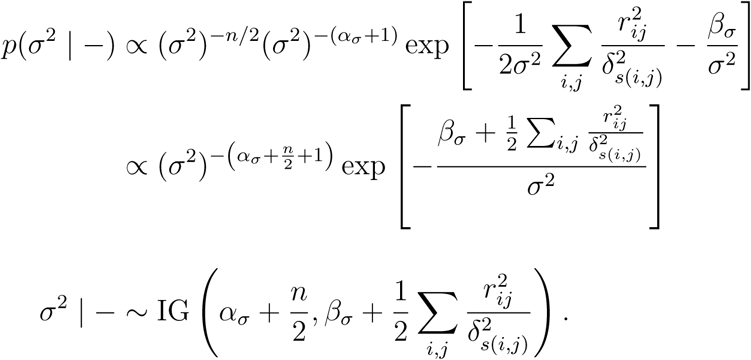

**Scanner-specific variance multipliers** *δ_s_*^2^

Relevant conditional density and prior

For all (*i, j*) such that *s*(*i, j*) = *s*

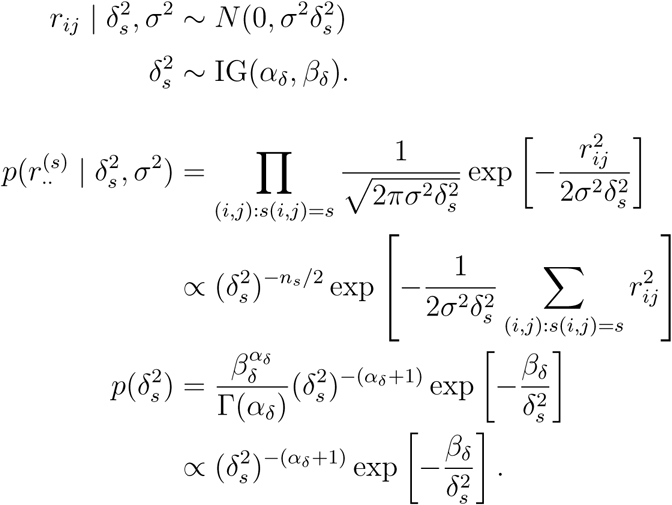

Conditional posterior distribution

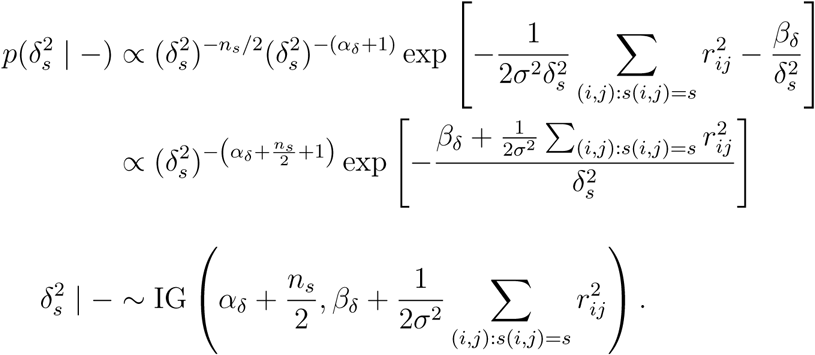

**Subject-specific random effects** *α_i_*

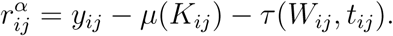

Relevant conditional density and prior

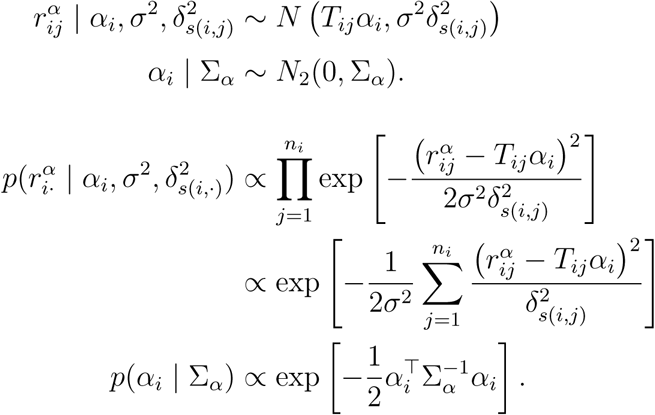

Conditional posterior distribution

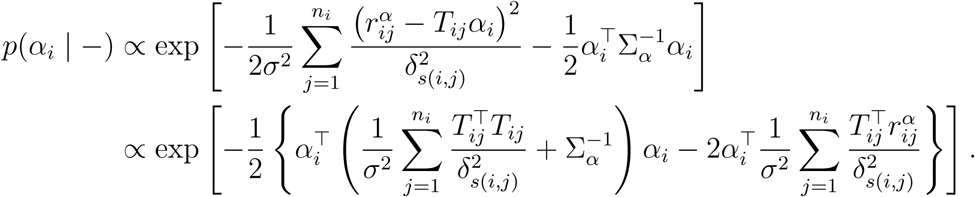

Completing the square, using

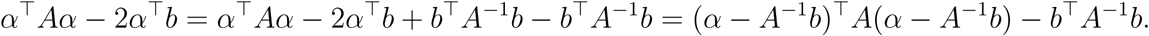

and defining 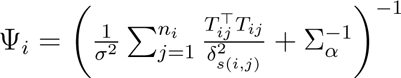,

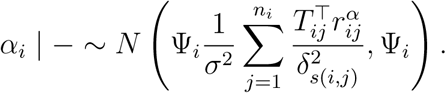

**Random-effects covariance** Σ*_α_*

Let *D* = 2 denote the dimension of the random-effects vector.

Relevant conditional density and prior

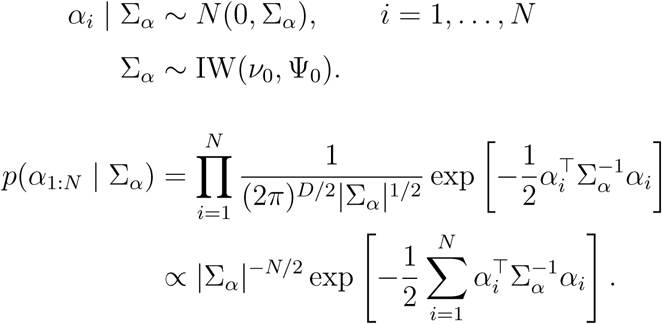

Using 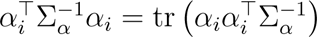,

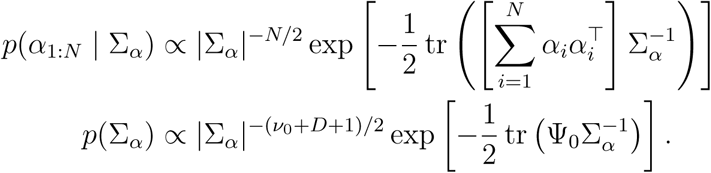

Conditional posterior distribution

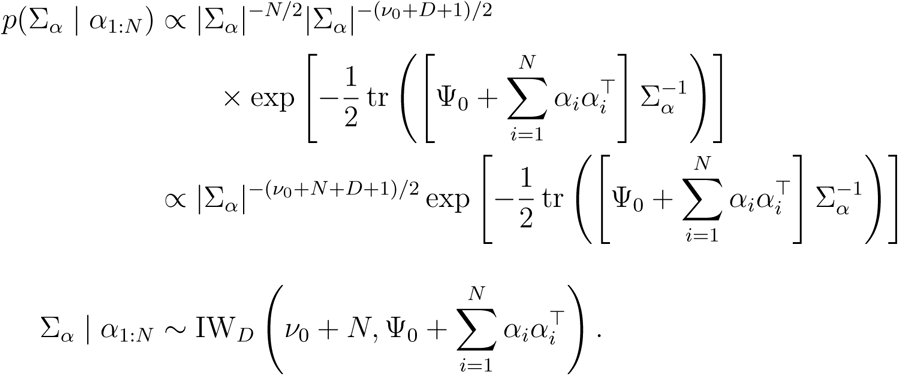

### 2 Real-data

Longitudinal harmonisation analyses compare unharmonised data, LongComBat, and Long-BARTharm, examining the preservation of biological associations, regional effect estimates, and scanner-related residual variance heterogeneity. Posterior predictive harmonisation and uncertainty weighting are further evaluated for both cross-sectional BARTharm and Long-BARTharm, with comparisons against unharmonised data and the corresponding ComBat methods. Posterior uncertainty across individual observations within cortical regions is also examined.

